# The genome of the zoonotic malaria parasite *Plasmodium simium* reveals adaptations to host-switching

**DOI:** 10.1101/841171

**Authors:** Tobias Mourier, Denise Anete Madureira de Alvarenga, Abhinav Kaushik, Anielle de Pina-Costa, Olga Douvropoulou, Qingtian Guan, Francisco J. Guzmán-Vega, Sarah Forrester, Filipe Vieira Santos de Abreu, Cesare Bianco Júnior, Julio Cesar de Souza Junior, Silvia Bahadian Moreira, Zelinda Maria Braga Hirano, Alcides Pissinatti, Maria de Fátima Ferreira-da-Cruz, Ricardo Lourenço de Oliveira, Stefan T. Arold, Daniel C. Jeffares, Patrícia Brasil, Cristiana Ferreira Alves de Brito, Richard Culleton, Cláudio Tadeu Daniel-Ribeiro, Arnab Pain

## Abstract

**Background:** *Plasmodium simium*, a malaria parasite of non-human primates (NHP) was recently shown to cause zoonotic infections in humans in Brazil. We sequenced the *P. simium* genome to investigate its evolutionary history and to identify any genetic adaptions that may underlie the ability of this parasite to switch between host species.

**Results:** Phylogenetic analyses based on whole genome sequences of *P. simium* from humans and NHPs reveals that *P. simium* is monophyletic within the broader diversity of South American *Plasmodium vivax*, suggesting *P. simium* first infected NHPs as a result of a host-switch of *P. vivax* from humans. The *P. simium* isolates show the closest relationship to Mexican *P. vivax* isolates. Analysis of erythrocyte invasion genes reveals differences between *P. vivax* and *P. simium*, including large deletions in the Duffy Binding Protein 1 (DBP1) and Reticulocyte Binding Protein 2a genes of *P. simium*. Analysis of *P. simium* isolated from NHPs and humans revealed a deletion of 38 amino acids in DBP1 present in all human-derived isolates, whereas NHP isolates were multi-allelic.

**Conclusions:** Analysis of the *P. simium* genome confirmed a close phylogenetic relationship between *P. simium* and *P. vivax*, and suggests a very recent American origin for *P. simium*. The presence of the DBP1 deletion in all human-derived isolates tested suggests that this deletion, in combination with other genetic changes in *P. simium*, may facilitate the invasion of human red blood cells and may explain, at least in part, the basis of the recent zoonotic infections.

## Background

There are currently eight species of malaria parasites known to cause disease in humans: *Plasmodium falciparum, Plasmodium vivax, Plasmodium malariae, Plasmodium ovale curtisi, Plasmodium ovale wallikeri, Plasmodium knowlesi, Plasmodium cynomolgi* and *Plasmodium simium*. The last three species are non-human primate (NHP) parasites that have recently been shown to infect humans [1–3].

As interventions against human parasites, particularly *P. falciparum* and *P. vivax*, reduce their prevalence, the importance of zoonotic malaria is becoming increasingly apparent. In countries moving towards malaria elimination, the presence of potentially zoonotic parasite populations in NHPs is a significant obstacle.

The propensity of malaria parasites to switch hosts and the consequences of this for human health are highlighted by the fact that both *P. vivax* and *P. falciparum* first arose as human pathogens following host switches from non-human great apes in Africa [4–6]. Contact between humans and the mosquitoes that feed on NHPs is increasing due to habitat destruction and human encroachment into NHP habitats [7]. Therefore, there is increasing danger of zoonotic transmission and the emergence of novel human malaria pathogens. Understanding how malaria parasites adapt to new hosts and new transmission environments allows the risks posed by novel zoonotic malaria outbreaks to be assessed.

The clinical epidemiology of zoonotic malaria varies according to the parasite species and the demographics of the human host population. Severe and fatal outcomes for people infected with *P. knowlesi* in Malaysia have been reported [8], whilst *P. cynomolgi* infection in the same region causes moderate or mild clinical symptoms [9]. Interestingly, both *P. knowlesi* and *P. cynomolgi* infections in the Mekong region appear less virulent than in Malaysia, and are often asymptomatic [3, 10]. This may be due to differences in the relative virulence of the circulating parasite strains and/or in the susceptibility of the local human populations. As NHP parasites have adapted to co-evolved with their natural hosts, it is impossible to predict their virulence in zoonotic infections. For example, the pathogenicity of *P. falciparum* has been attributed to its relatively recent emergence as a human pathogen [11], which is thought to have occurred following a single transfer from a gorilla in Africa [5].

Eighty-nine percent of malaria infections in Brazil are caused by *P. vivax* and over 99.9% of these cases occur in the Amazonian region. This region accounts for almost 60% of the area of Brazil and is home to 13% of the human population (https://www.ibge.gov.br/). Around 90% of the malaria cases registered outside the Amazonian region occur in the Atlantic Forest, a region of tropical forest that extends along the Brazil’s Atlantic coast. The infections are apparently mild and are caused by a vivax-like malaria parasite transmitted by *Anopheles (Kerteszia) cruzii*, a mosquito species that breeds in the leaf axils of bromeliad plants [12].

A malaria outbreak in the Atlantic Forest of Rio de Janeiro in 2015/2016 was shown to be caused by the NHP parasite *P. simium* [1]. Parasite DNA samples collected from both humans and NHPs in the same region had identical mitochondrial genome sequences, distinct from that of *P. vivax* isolates collected anywhere in the world but identical to that of a *P. simium* parasite isolated from a monkey in the same region in 1966, and to all subsequent *P. simium* isolates recovered from NHPs [13, 14].

Previously, it was thought that *P. vivax* became a human parasite following a host switch from macaques in Southeast Asia, due to its close phylogenetic relationship with a clade of parasites infecting macaques in this region, and due to the high genetic diversity among *P. vivax* isolates from Southeast Asia [15]. However, it is now considered likely that it became a human parasite following a host switch from non-human great apes in Africa [6]. It seems likely that *P. vivax* was introduced to the Americas by European colonisers following Columbus’ journey to the New World towards the end of the 15^th^ Century, since present-day South American *P. vivax* is closely related to a strain of the parasite present, historically, in Spain [16]. However, there is some evidence to suggest that *P. vivax* may also have been introduced to South America in pre-Columbian times [17], and this, together with the post-Columbian influx of infected people from various regions of the world associated with colonization, may have contributed to the extensive genetic diversity of present day *P. vivax* in Central and South America [17].

*Plasmodium simium*, a parasite of various species of Platyrrhini monkey whose range is restricted to the Atlantic Forest from Southeast and South of Brazil [18], is genetically, morphologically, and immunologically similar to *P. vivax* [1, 19–22]. Based on this similarity, it is likely that the *P. simium* parasite of NHPs in Brazil originated from *P. vivax* following a host switch from humans. The recent 2015/2016 outbreak of *P. simium* in the local human population of Rio de Janeiro’s Atlantic Forest raises questions about the degree of divergence between *P. vivax* and *P. simium*, and whether adaptation to NHPs has led to the evolution of a parasite with relevance to human health that differs from that of *P. vivax*.

It remains unproven whether the current outbreak of *P. simium* in the human population of Rio de Janeiro was the result of a single parasite transfer from a NHP to a human and its subsequent transfer between people, or whether multiple independent transfers have occurred with different NHP-derived parasites. Furthermore, the nature and degree of adaptation to NHP hosts and a sylvatic transmission cycle that has occurred in *P. simium* following its anthroponotic origin is relevant to understanding how malaria parasites adapt to new hosts. It is of interest to determine whether the current, human-infecting *P. simium* parasites have undergone recent changes at the genomic level which allow them to infect humans, as it has been previously suggested that *P. simium* lacks this ability [23].

In order to address these questions, and so to better understand the epidemiology and natural history of this emerging zoonotic parasite, we analysed whole genome sequences of *P. simium* parasites isolated from both humans and NHPs.

## Results

### Genome assembly and phylogeny

From a single *P. simium* sample collected from Rio de Janeiro state in 2016 [1] short read DNA sequences were obtained and assembled into a draft genome (see Supplementary Materials). The assembled genome consists of 2,192 scaffolds over 1kb in length with a combined size of 29 Mb (Table S1). Two scaffolds corresponding to the apicoplast and mitochondrial genomes were also identified (Figure S1). Gene content analysis showed a completeness of annotation comparable to previously published *Plasmodium* assemblies (Figure S2). Hence, although the fragmented nature of the chromosome assemblies prevents precise assessment of synteny conservation with other *Plasmodium* species, the set of gene components in the *P. simium* genome is relatively complete. A phylogenetic tree constructed from 3,181 of 1:1 orthologs of the annotated *P. simium* protein-coding genes with the orthologues of *P. vivax, P. cynomolgi, P. coatneyi, P. knowlesi, P. malariae, P. falciparum, P. reichenowi*, and *P. gallinaceum* confirmed that *P. simium* is very closely related to *P. vivax* (Figure S3). Analysis of gene families revealed a *P. simium* gene repertoire largely similar to *P. vivax* (Figure S4; Table S2) and no indication of a recent expansion of the *P. simium* PIR gene family (Figure S5).

### *P. simium-P. vivax* diversity analysis

To detect single nucleotide polymorphisms (SNPs) within the *P. vivax/P. simium* clade, short Illumina paired-end sequence reads were mapped onto the *P. vivax* P01 reference genome [24]. Reads were collected from eleven human-derived *P. simium* samples, two monkey *P. simium* samples, two *P. vivax* samples from the Brazilian Amazon, and a previously published *P. simium* CDC strain (originally isolated in 1966, see Methods). Data from a range of *P. vivax* strains representing a global distribution was retrieved from the literature [25]. Including only SNPs with a minimum depth of five reads, a total of 232,780 SNPs were called initially across 79 samples. Sixteen samples were subsequently removed, primarily due to low coverage, resulting in a total of 63 samples for further analysis (Table S3, Table S4). Since few SNP loci were covered across all samples, and to enable diversity analysis, we restricted all further analysis to the 124,968 SNPs for which data were available from at least 55 samples (Figure S6).

### *P. simium-P. vivax* population analysis

A Principal Component Analysis (PCA) plot constructed from these genome-wide SNP loci showed a clear separation between American and Asian *P. vivax* samples as well as a distinct grouping of *P. simium* samples (Figure S7). The latter observation suggests that both human and NHP *P. simium* isolates represent a single population that is genetically different from American *P. vivax* populations. A similar pattern is observed by performing a multidimensional scaling analysis of the SNP data (Figure S8). To enable a phylogenetic approach, we constructed an alignment from the 124,968 SNP sites. In the resulting phylogenetic tree, *P. vivax* strains generally clustered according to their geographical origin, with clear separation of the Asian and American samples (Figure 1A, a tree with sample IDs is available in Figure S9). *Plasmodium simium* samples clustered as a monophyletic group with Mexican *P. vivax* samples (Figure 1A), consistent with a recent American origin for *P. simium*. This phylogeny, based on genome-wide SNPs represents an “average” phylogeny across the genome and cannot be considered to reflect a true history of parasite ancestry due to the effects of recombination. It is possible that trees produced from individual genes might reveal different phylogenetic relationships. Yet, the association between *P. simium* samples and *P. vivax* samples from Mexico is also observed in a phylogenetic network (Figure S10).

**Figure 1.**
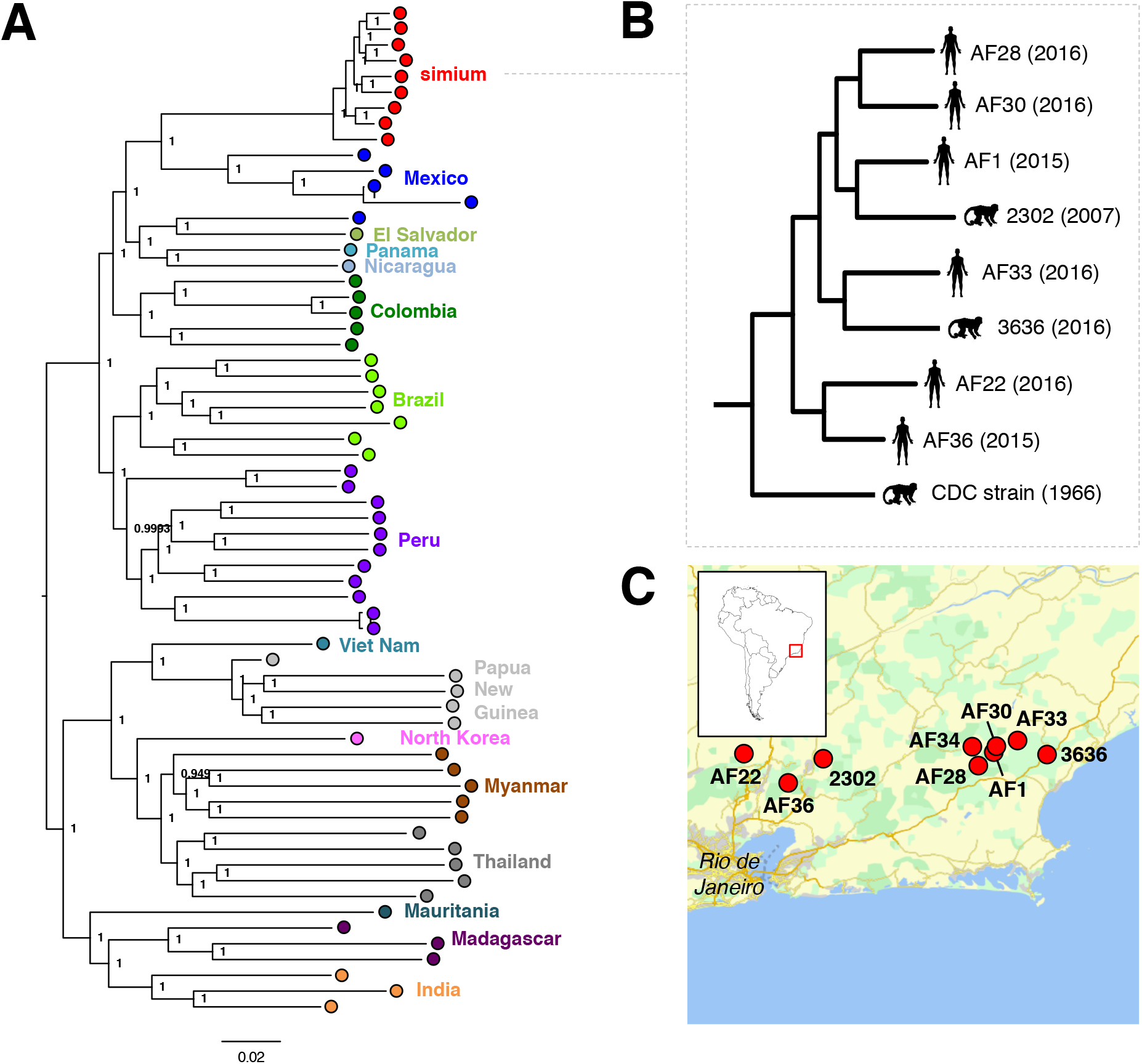
SNP phylogeny. A) Mid-point rooted maximum likelihood tree produced from 143,123 concatenated SNP positions with data from a minimum of 55 samples. The tree was produced using PhyML with the GTR evolutionary model. Branch support was evaluated with the Bayesian-like transformation of approximate likelihood ratio test (aBayes). Genetic distance is shown below the tree. *Plasmodium vivax* isolates are denoted as coloured circles by their country of sample origin. A tree with specific sample IDs is available in Figure S9B) Magnification of the *Plasmodium simium* clade (as in panel A). C) Map showing the geographic locations at which the *P. simium* samples were collected.

To examine whether the *P. simium* isolates we studied were part of a continuous population with local *P. vivax* parasites, we examined population ancestry with the ADMIXTURE program [26] (Figure S11). This analysis is consistent with the PCA and MDS analyses (Figure S7 & Figure S8) and the phylogenetic analysis of segregating SNPs (Figure 1), showing that *P. simium* forms a genetically distinct population separate from *P. vivax*. The absence of *P. simium-P. vivax* hybrids (indicating genetic introgression events) indicates that *P. simium* has undergone a period of independent evolution.

### Genetic diversity and population divergence

To characterise the *P. simium* population further, we estimated the nucleotide diversity in *P. simium* and *P. vivax* samples (see Materials and Methods). *Plasmodium simium* diversity (genome-median: 1.3 × 10^−4^) is approximately six times lower than the diversity observed when comparing all *P. vivax* samples (genome-median: 7.9×10^−4^) (Figure 2A). Diversity measured within coding sequences in *P. vivax* is consistent with previous reports [6]. The median nucleotide divergence between *P. simium* and *P. vivax* genomes of 8.7× 10^−4^ and the low diversity within *P. simium* suggest that the strains we examined are part of a relatively recent or isolated population.

**Figure 2.**
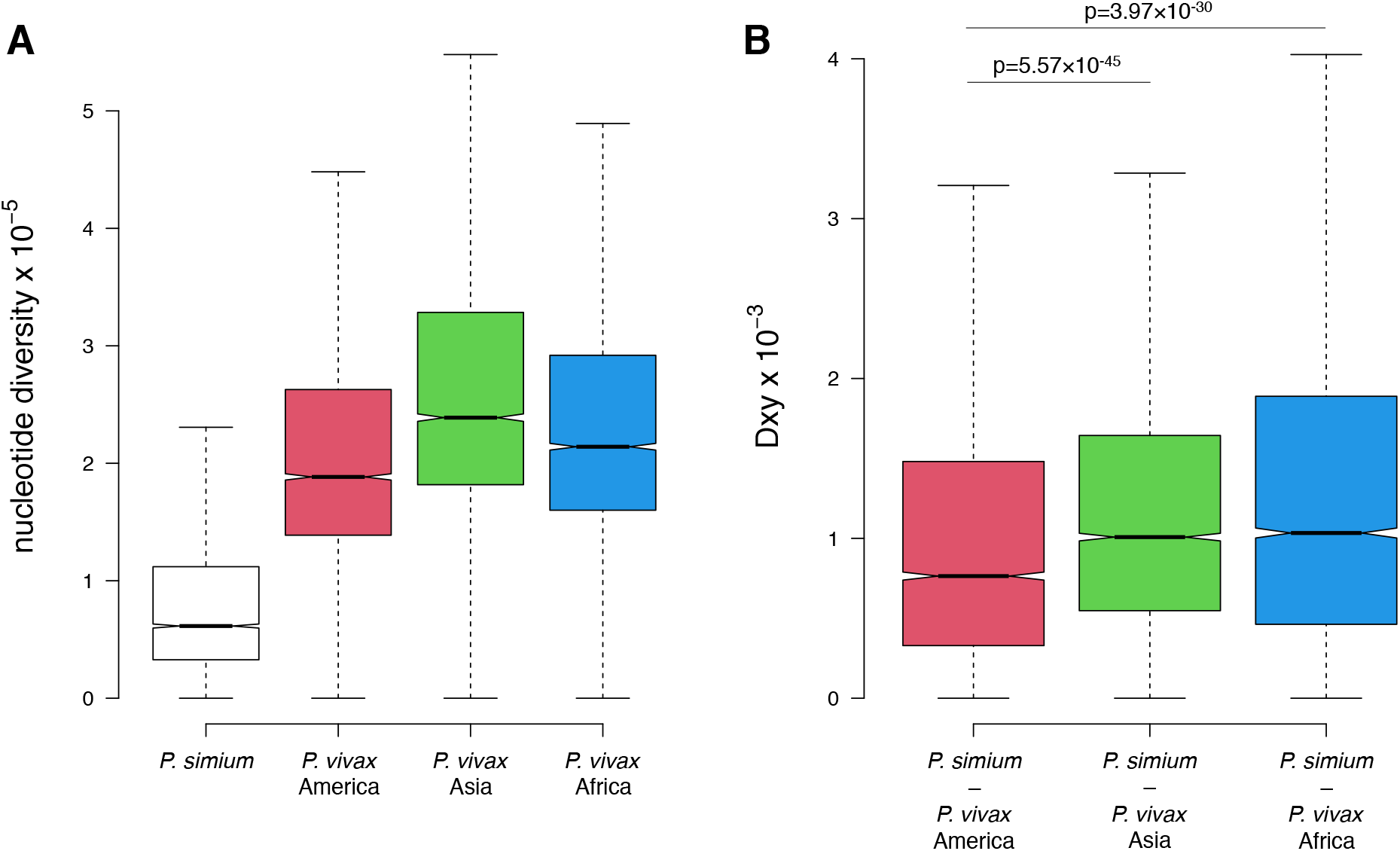
Nucleotide diversity and genetic distance between populations. A) Diversity within populations. Box plot showing the nucleotide diversity in 10 kb windows between *Plasmodium simium* samples (left), and *Plasmodium vivax* samples from America, Asia, and Africa. B) Genetic distance between populations. Box plot showing D_XY_ in 10 kb windows comparing *Plasmodium simium* samples to P. vivax samples from America, Asia, and Africa. P-values from one-sided Mann-Whitney tests for difference in medians are shown above boxes. For both plots, boxes denote 25th and 75th percentiles with all outliers removed.

The difference in diversity within *P. simium* and *P. vivax* populations will influence relative measures of population differentiation such as FST [27], and we therefore calculated D_XY_, an absolute measure of diversity that is independent of the levels of diversity within the two populations being compared [27, 28]. When comparing D_XY_ in 10kb windows across the genome, the diversity observed between *P. simium* samples and *P. vivax* samples from America is significantly lower than the diversity between *P. simium* and *P. vivax* populations from Asia and Africa (Figure 2B), supporting the origin of *P. simium* from American *P. vivax* populations.

The vast majority of genes have an D_XY_ of zero, consistent with a recent split between *P. simium* and *P. vivax* (Figure S12A). We noticed that genes showing the highest D_XY_ are from multi-gene families (Figure S12B). Although polymorphic antigens are expected to display high levels of diversity, this could also be explained by the uncertainties in SNP calling among multi-gene families with high levels of sequence redundancy.

### *P. simium* invadome components

Binding and red blood cell invasion is mediated by two key malaria gene families expressed in merozoites: the Duffy Binding Proteins (DBPs) and the Reticulocyte Binding Proteins (RBPs). In *P. vivax*, DBPs bind to the Duffy Antigen Receptor for Chemokines (DARC) [29, 30], which is present on both host normocytes and reticulocytes, and RBPs are known to restrict *P. vivax* to binding and invading reticulocytes [31–33]. Two DBPs, DBP1 and DBP2, are present in *P. vivax* P01 (Table S5). Recently, the reported protein structure of *P. vivax* RBP2b revealed the conservation of residues involved in the invasion complex formation [33]. RBPs can be divided into three subfamilies, RBP1, RBP2, and RBP3 [34]. The *P. vivax* P01 genome encodes 11 RBPs (including the reticulocyte binding surface protein, RBSA), of which three are pseudogenes (Table S5).

The *P. vivax* DBP and RBP protein sequences were used to search for *P. simium* orthologues, resulting in the detection of the two DBP proteins and RBP1a, RBP1b, RBP2a, RBP2b, and RBP3, and a failure to detect RBP2c and RBP2d (Figure 3; Table S5; Figure S13; Figure S14) across all sequenced *P. simium* samples. As in *P. vivax* genomes, the *P. simium* RBP3 is a pseudogene [35], indicating that conversion to a pseudogene happened prior to the split between *P. vivax* and *P. simium*.

**Figure 3.**
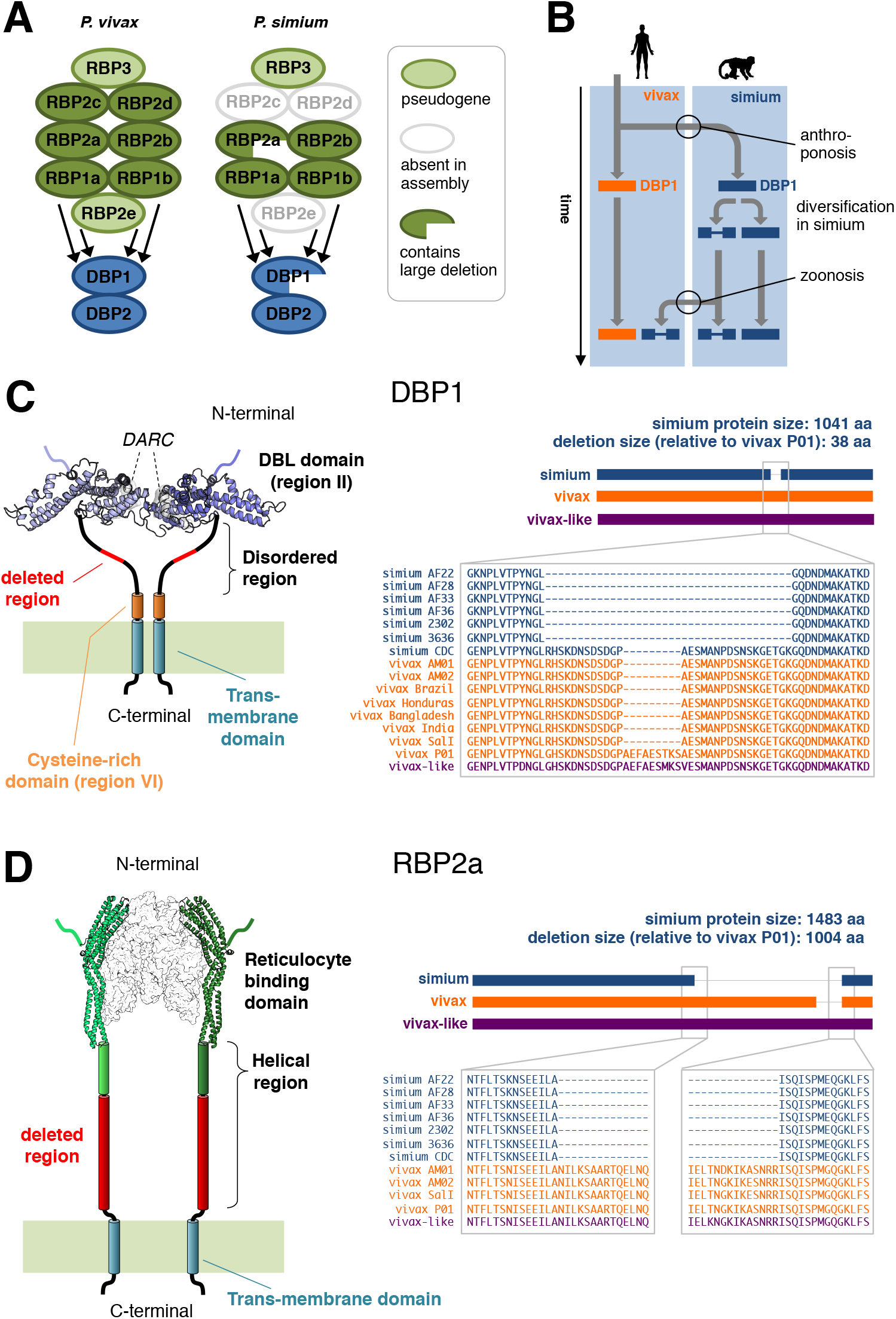
Invadome deletions. A) Overview of the invadome gene groups, Reticulocyte Binding Proteins (RBPs) and Duffy Binding Proteins (DBPs) in *Plasmodium vivax* and *Plasmodium simium*. The *P. vivax* genome harbours two RBP2d genes, one of which is a pseudogene (Table S5). B) Schematic depiction of the hypothesized scenario in which the DBP1 deletion – along with other accumulated genetic changes in *P. simium* – is a prerequisite for the recently observed zoonosis. C) Left: Structural rendering of DBP1, showing known structural domains and motifs. The two fragment molecules from the human DARC receptor are shown in grey. The 3dimensional structure of the DBL-DARC complex was modelled based on the *P. vivax* crystallographic model (PDB 4nuv). The region deleted in sequences from human-infecting *P. simium*, as compared to *P. vivax* P01, is highlighted in red. Right: Details of DBP1 protein alignments. A full alignment is available in Figure S20. D) Similar to panel C) but for RBP2a. The complex between the reticulocyte binding domain and the human receptor was modelled based on the cryoEM structure of the complex between the *P. vivax* RBP2b and the human transferrin receptor TfR1 (PDB 6d05). A full alignment is available in Figure S28.

To determine whether the apparent absences of individual RBP genes in *P. simium* was due to incomplete genome assembly, we examined the coverage of *P. simium* reads mapped onto *P. vivax* RBP gene loci. No *P. simium* coverage was observed at the RBP2c, RBP2d, and RBP2e genes in *P. simium* samples, including the previously published CDC strain (Figure S15).

Coverage of mapped reads across invadome-associated gene loci revealed no apparent elevated coverage in genes compared to their flanking genomic regions, which would have been expected if the *P. simium* genome contained multiple (duplicated) copies of non-assembled invasion genes (Figure S16).

### Structural variation in *P. simium* Duffy Binding Protein 1

The *P. simium* assembly revealed that the invasion gene DBP1 contains a large deletion within its coding sequence (Figure 3; Figure S17). Intriguingly, the previously published *P. simium* CDC strain DBP1 (GenBank accession: ACB42432) [36] does not contain this deletion (‘simium CDC’ in Figure 3C). A haplotype network confirms that this previously published DBP1 gene is indeed a *P. simium* sequence (Figure S18), and the SNP analyses consistently assign the CDC strain to the *P. simium* cluster (Figure 1, Figure S7, and Figure S8). Compared to the *P. vivax* P01 reference genome the SalI reference harbours a 27 base pair deletion in DBP1, in contrast to the 115 bp deletion observed in all *P. simium* samples isolated from humans (Figure 3). The 27 base pair deletion is also present in most *P. vivax* isolates (Figure S19). Additional deletion patterns exist among isolates, and in a few cases multiple versions are detected within samples (Figure S19).

The presence of repetitive sequences within the DBP1 gene could potentially result in aberrant assembly across the DBP1 locus, which may appear as an apparent deletion in subsequent bioinformatic analysis. We tested this possibility and showed that the DBP1 gene does not harbour any noticeable degree of repetitiveness at the nucleotide level (Figure S20). Several read mapping analyses confirmed that the *P. simium-specific* 115 bp deletion was not an assembly artefact (Figure S21-S23).

We next designed primers for PCR amplification of a nucleotide sequence that spans the deleted region in the *P. simium* DBP1 gene and tested the occurrence of these deletion events in a range of *P. vivax* and *P. simium* field samples from Brazil. All *P. vivax* samples tested by PCR produced bands consistent with absence of the deletion whereas all samples from human-infecting *P. simium* produced bands consistent with the presence of the precise 115 bp deletion (Figure S24, top & middle). Interestingly, NHP-infecting *P. simium* isolates contained a mix of sequences with and without deletions (Figure S24, bottom). PCR results are summarized in Table S6. Under the simplified assumption that the sampled NHP *P. simium* samples reflect the DBP1 allele frequency in the entire *P. simium* population (4 with deletion in DBP1, 8 without the deletion; Table S6), the probability of randomly sampling 15 human *P. simium* samples all having the DBP1 deletion is 6.97× 10^−8^ using a binomial distribution. If the *P. simium-specific* deletion in DBP1 is a prerequisite for the ability to infect humans, this suggests that only a subset of NHP-infecting *P. simium* parasites currently possess the ability to infect humans.

A large, 3012 bp additional deletion was observed in the *P. simium* RBP2a gene, the presence of which was also supported by read mapping (Figure 3, Figure S25-S28). PCR-based genotyping results confirmed the presence of this deletion in all *P. simium* isolates irrespective of their host (Figure S29, Table S6).

To test for the presence of insertions and deletions (indels) in other protein-coding genes, we curated a set of indels by comparing short indels predicted by the DELLY software [37] to *P. simium* assemblies. This set consisted of 244 indels in 222 genes (Table S7). Indel sizes were almost exclusively integers of three (hence conserving reading frames) and indels were predominantly found in low-complexity regions and in genes encoding long proteins (Figure S30). Notably, short indels were recorded in 138 genes with known functional annotations with an overrepresentation of DNA-binding function (GO:0005488 ‘binding’, p=0.0011; GO:0043565 ‘sequence-specific DNA binding’, p=0.0062).

### Potential structural implications of the deletions in DBP1 and RBP2a

We next investigated whether the observed deletions render DBP1 and RBP2a nonfunctional. DBP1 contains a large extracellular region, which includes the N-terminal DBL region which mediates the association with DARC in *P. vivax* [38], followed by a largely disordered region and a cysteine-rich domain (Figure 3C). DBP1 has a single-pass transmembrane helix and a short cytoplasmic tail. The deletion observed in the human-infecting *P. simium* only affects the disordered region, leaving the flanking domains intact. We produced homology models of the DBL domains from the *P. vivax* strain P01, the human-infecting *P. simium* strain AF22, and the *P. simium* CDC strain, based on the crystal structure of the > 96% identical DBL domain of *P. vivax* bound to DARC (PDB ID 4nuv). Whereas no significant substitutions were found in the DBL domain between both *P. simium* sequences, our analysis showed that residue substitutions between *P. simium* and *P. vivax* DBL domains cluster in proximity of the DARC binding site (Figure S31). Based on our models, these substitutions are unlikely to negatively affect the association with DARC, supporting the idea that the DBL domains of both *P. simium* would be capable of binding to human DARC. Hence, the human-infecting *P. simium* DBL1 probably retains the capacity to bind to human DARC, but has the interacting domain positioned closer to the membrane than in the NHP-infecting CDC strain.

The deletion we detected in the *P. simium* RBP2a was larger, resulting in the loss of 1003 amino acid residues. These residues are predicted to form a mostly α-helical extracellular stem-like structure that positions the reticulocyte binding domain away from the membrane (Figure 3D). However, given that the deletion affects neither the transmembrane region, nor the receptor-binding domain, our analysis suggests the resulting truncated RBP2a protein could still associate with the human receptor, but that the binding event would occur closer to the merozoite membrane.

## Discussion

We present the genome of *Plasmodium simium*, the eighth malaria parasite species known to infect humans in nature. In recent evolutionary time, *P. simium* has undergone both anthroponosis and zoonosis making it unique for the study of the genetics underlying host-switching in malaria parasites. Analysis of its genome confirmed a close phylogenetic relationship between *P. simium* and *P. vivax*, and further analyses on single nucleotide divergence support a very recent American origin for *P. simium*.

Two proteins involved in host invasion, DBP1 and RBP2a, were found to harbour extensive deletions in *P. simium* compared to *P. vivax*. Interestingly, experimental analysis of *P. simium* samples revealed that isolates from human hosts all carried the DBP1 deletion, whereas isolates from NHPs displayed both absence and presence of the deletion. This DBP1 deletion is not present in the *P. simium* isolated from a brown howler monkey in the 1960s, which was previously shown to be incapable of infecting humans [23], although some degree of laboratory adaptation of this parasite may have affected its genome. However, this deletion is also absent in *P. vivax*, so cannot in itself explain the ability of *P. simium* to infect humans in the current outbreak. It is possible, however, that this deletion is required for *P. simium* to invade human red blood cells given the alterations that have occurred elsewhere in its invadome following adaptation to non-human primates since the split between *P. simium* and its human-infecting *P. vivax* ancestor (Figure 3B).

Our data is consistent with the hypothesis that the DBP1 deletion is required for efficient invasion of human RBCs by the *P. simium* parasites present in the Atlantic Forest. However, another study from Espírito Santo, suggests that humans may be infected with *P. simium* strains that carry DBP1 without this deletion [39]. This discrepancy may be caused by genetic differences in the *P. simium* strains circulating in each region. Further, the distinction between *P. simium* and *P. vivax* infections based on epidemiological data may be confounded by human migration between the Amazonas and Espírito Santo [40].

Invadome proteins are obvious candidates for genetic factors underlying host-specificity; an inactivating mutation in a *P. falciparum* erythrocyte binding antigen has recently been shown to underlie host-specificity [41]. Traditionally, functional studies on invadome proteins have focused on domains known to bind or interact directly with the host. Although the *P. simium*-specific DBP1 and RBP2a deletions reported here do not cover known structural motifs, these deletions could nevertheless affect host cell recognition as disordered protein regions have known roles in cellular regulation and signal transduction [42]. Further, a shorter, less flexible linker between the merozoite membrane and the receptor-binding DBP1 domain may favour a more rigid and better oriented positioning of the dimeric DBP1, enhancing its capacity to engage the human receptor.

Phylogenetic analyses of the *P. simium* clade give the geographical location of its most closely related *P. vivax* strain as Mexico, and not Brazil. In imported populations, the relationship between geographical and genetic proximity may be weak. Multiple introductions of diverse strains from founder populations may occur independently over large distances, so that two closely related strains may be introduced in distantly located regions. It may be postulated that there occurred the introduction of strains of *P. vivax* to Mexico from the Old World that were closely related (due to similar regions of origin) to strains introduced to the Atlantic Forest which then went on to become *P. simium* in New World monkeys. Strains from a different point of origin were introduced to the Amazonian region of Brazil. This hypothesis necessitates reproductive isolation of the *P. simium* clade from the Brazilian *P. vivax* parasites following their initial introduction; an isolation that would be facilitated, presumably, by their separate host ranges or via adaptation to different vectors.

Due to uncertainties regarding the number of individual genomes that were transferred during the original host switch from human to NHPs that resulted in the formation of the *P. simium* clade, it is impossible to perform dating analyses to determine a time for the split between *P. vivax* and *P. simium* with which we can be confident. The phylogeny shown in Figure 1 is consistent with the hypothesis that all present-day *P. vivax/P. simium* originated from a now extinct, or as yet unsampled, Old World population. The most parsimonious explanation for this is that today’s New World *P. vivax/P. simium* originated from European *P. vivax*, which was itself a remnant of the original Eurasian/African *P. vivax* driven to extinction (or near extinction) in Africa by the evolution of the Duffy negative condition in the local human populations, and from Europe by malaria eradication programmes in the latter half of the twentieth century. This hypothesis is supported by the evidence of a close relationship between historical Spanish *P. vivax* and South American strains of the parasite [16], and by previous analyses of the mitochondrial genome [43]. Therefore, we postulate that the host switch between humans and non-human primates that eventually led to establishment of *P. simium* in howler monkeys must have occurred subsequent to the European colonisation of the Americas, within the last 600 years.

We find no evidence from the nuclear genome, the mitochondrial genome or the apicoplast genome that any of the *P. vivax /P. simium* strains from the New World considered in our analyses are more closely related to Old World parasites than they are to each other, as previously contended [44].

Given the limited genetic diversity amongst the *P. simium* isolates considered here compared to that of *P. vivax*, it is almost certain that the original host switch occurred from humans to NHPs, and not the other way around [22]. Similarly, the larger amount of genetic diversity in the current NHP-infecting *P. simium* compared to those *P. simium* strains isolated from humans (as indicated by the higher degree of DBP1 polymorphism in the NHP-infecting *P. simium* compared to the strains infecting humans), suggests that humans are being infected from a pool of NHP parasites in a true zoonotic manner, as opposed to the sharing of a common parasite pool between humans and NHPs.

The biological definition of a species is a group of organisms that can exchange genetic material and produce viable offspring. We have no way of knowing whether this is the case for *P. vivax* and *P. simium*, and genetic crossing experiments would be required to resolve this question. Our phylogenetic analyses, however, clearly show *P. simium* forming a clade on its own within the broader diversity of *P. vivax*, and this strongly suggests, given what we know about its biology, that allopatric speciation has been/is occurring.

*Plasmodium simium* is currently recognised as a species separate from *P. vivax*; it has been well characterised and described in the literature, and there is a type specimen available, with which all the strains sequenced here cluster in one monophyletic group. Therefore, we cannot at present overturn the species status of *P. simium* in the absence of conclusive proof from crossing experiments.

## Conclusions

The recent outbreak of human malaria in the Atlantic Forest of Rio de Janeiro underlines the impact of zoonotic events on human health. Non-human primate malaria parasites must be considered a reservoir of potential infectious human parasites relevant to any malaria eradication strategy. Little is known about the genetic basis for zoonoses, yet the genome sequence of *P. simium* suggests a deletion within the DBP1 gene is a possible facilitator of zoonotic transfer. The genome of *P. simium* will thus form an important resource for the future functional characterizations of the mechanisms underlying zoonotic malaria.

## Methods

### Sample Collection and Preparation

Human and primate samples of *P. simium* were collected and prepared as part of a previous study [1, 14]. Additionally, two *P. vivax* samples from the Amazon area of Brazil were also collected from human patients (Table S3). All participants provided informed written consent. The *P. simium* CDC (Howler) strain (Catalog No. MRA-353) from ATCC was obtained *via* the BEI Resources Repository in NIAID-NIH (https://www.beiresources.org/).

### DNA extraction and sequencing

DNA was extracted as previously described[1]. Genomic DNA for each sample was quantified using the Qubit® 2.0 Fluorometer and was used for library preparation. DNA for intact samples was sheared using a Covaris E220 DNA sonicator to fragments of 500bp. The DNA libraries for intact samples were made using the TruSeq Nano DNA Library Prep kit (Illumina), whereas the DNA libraries for degraded samples were made using Ovation Ultralow Library System V2 kit (Nugen), according to the manufacturers’ instructions. The amplified libraries were stored at −20 °C. The pooled libraries were sequenced in an Illumina HiSeq4000 instrument (2 x 150 bp PE reads) (Illumina). A PhiX control library was applied to the sequencing run as a base balanced sequence for the calibration of the instrument so that each base type is captured during the entire run. Samples AF22, AF26, AF36 were additionally sequenced and scaffolded by PacBio RS II platform (Pacific Biosciences, California, US) using a SMRT library. Genomic DNA from the *P. vivax* samples was extracted from filter paper as previously described [45].

### Illumina reads preparation and mapping

FastQC v 0.11.6 (http://www.bioinformatics.babraham.ac.uk/projects/fastqc) was used to evaluate the quality of Illumina reads. Illumina adapters were removed, and reads were trimmed using the trimmomatic v0.33 [46] software with the following conditions:

*LEADING:20 TRAILING:20 SLIDINGWINDOW:4:20 MINLEN:36*

To exclude human reads from our analysis, trimmed reads were mapped against the human reference genome (v. hg38) and the *Plasmodium vivax* strain P01 reference genome (v. 36) from PlasmoDB (www.plasmodb.org) with bowtie2 (v 2.3.3.1) [47]. Reads mapping against the human genome were removed from further analysis.

### Genome Assembly

*Plasmodium simium* sample AF22 was selected for genome assembly based on read quality and coverage. After removal of human contaminants, Illumina reads were assembled into contigs using the Spades (v 3.70) assembler [48]. Contigs assembled into scaffolds running SSPACE (v 3.0) [49] for 15 rounds and gaps filled with Gapfiller (v 1.10) [50]. Scaffolds were subsequently corrected with Illumina reads using the Pilon (v 1.22) software [51]. Blobtools (v 1.0) (DOI: 10.5281/zenodo.845347) [52] was used to remove any residual contaminant scaffolds. Genome size and GC content was in line with that of *P. vivax* (Table S1). Genomic scaffolds representing the mitochondrial and apicoplast genome were identified through blastn searches against the corresponding *P. falciparum* and *P. vivax* sequences (Figure S1). The *P. simium* mitochondrial genome was aligned against a range of previously published *P. vivax* and *P. simium* mitochondrial genomes [53, 54]. A gap-filled region in the alignment where the distal parts of the *P. simium* scaffold were merged was manually deleted. A minimum spanning haplotype network was produced using PopART [55, 56] confirming the authenticity of the *P. simium* mitochondrial genome (Figure S32).

### Genome Annotation

Two approaches were used to annotate the reference *P. simium* AF22 genome. Firstly, the Maker pipeline (v 2.31.8) [57] was run for two rounds, using ESTs and protein evidence from *P. vivax* and *P. cynomolgi* strain B and *P. falciparum* to generate Augustus gene models. Secondly, a separate annotation was produced using the Companion web server [58]. Companion was run using the *P. vivax* P01 reference assembly and default parameters. Basic annotation statistics are provided in Table S1. The relatively low number of genes (5966) is due to the fragmented and incomplete nature of the *P. simium* assembly (Table S1). Gene content was estimated using BUSCO [59, 60] (v3.0) revealing a gene annotation completeness comparable to other *Plasmodium* genome assemblies (Figure S2).

### PlasmoDB Genome References and Annotations

Genome FASTA files, as well as annotated protein and CDS files were obtained from PlasmoDB [61] for the following species: *P. gallinaceum* 8A, *P. cynomolgi* B and M, *P knowlesi* H, *P. falciparum* 3D7, *P. reichenowi* G01, *P. malariae* UG01, *P. ovale curtisi* GH01, *P. coatneyi* Hackeri, *P. vivax* P01 and *P. vivax* SalI. For each species, PlasmoDB version 36 was used.

### Orthologous group determination

Amino-acid sequences-based phylogenetic trees were prepared using protein sequences from the *P. simium* annotation, as well as the protein annotations from 10 malaria species downloaded from PlasmoDB: *P. vivax* P01, *P. cynomolgi* B, *P. knowlesi* H, *P. vivax-like* Pvl01, *P. coatneyi* Hackeri, *P. falciparum* 3D7, *P. gallinaceum* 8A, *P. malariae* UG01, *P. ovale* curtisi GH01, and *P. reichenowi* G01. *P. vivax*-like from PlasmoDB version 43, all other annotations from version 41. A total of 3181 1:1 orthologous genes were identified using the Proteinortho (v 6.0.3) software [62]. Approximately 88% of the predicted genes in *P. simium* have orthologues in *P. vivax* P01 (Figure S33).

### Short indels in genes

Shorter indels (<500 bp) were detected from soft-clipping information in read mapping using the ‘-i’ option in DELLY [37] (v 0.7.9). Predicted indels in protein-coding genes were then compared to independent assemblies of *P. simium* samples AF22, AF28, AF33, AF36, 2302 and 3636 (Table S3) and indels present in both DELLY predictions and all assemblies were kept (Table S7). Indel boundaries shifted by a maximum of two amino acid positions were allowed between predictions and assemblies. As assemblies were not complete, data for any gene was only required to be present in five of the six *P. simium* assemblies used.

### Protein phylogeny

Protein sequences were aligned using mafft (v 7.222) [63] and alignments were subsequently trimmed with trimAl (v 1.2rev59) [64] using the heuristic ‘automated1’ method to select the best trimming procedure. Trimmed alignments were concatenated and a phylogenetic tree was constructed using RAxML (v 8.2.3) [65] with the PROTGAMMALG model.

### SNP calling and analysis

Short sequence reads from 15 *P. simium* samples (13 derived from humans and two from NHPs) and two *P. vivax* samples, all from this study (Table S3), were aligned against a combined human (hg38) and *P. vivax* (strain P01, version 39) genome using NextGenMap (v0.5.5) [66]. This was similarly done for 30 previously published *P. vivax* strains [25] and the Sal1 reference. These data sets were downloaded from ENA (https://www.ebi.ac.uk/ena) (Table S4). Duplicate reads were removed using samtools (v 1.9) [67] and the filtered reads were realigned using IndelRealigner from the GATK package (v 4.0.11) [68]. SNPs were called independently with GATK HaplotypeCaller and freebayes (v 1.2.0) [69], keeping only SNPs with a QUAL score above 30. The final SNP set was determined from the inter-section between GATK and freebayes. Allele frequencies and mean coverage across SNP sites are shown in Figure S34. A PCA plot was constructed using plink (v 1.90) [70], and admixture analysis was done with Admixture (v 1.3.0) [26].

### SNP phylogeny

Alleles from SNP positions with data in 55 samples were retrieved, concatenated, and aligned using mafft [63]. Tree was produced by PhyML [71, 72] with the GTR substitution model selected by SMS [73]. Branch support was evaluated with the Bayesian-like transformation of approximate likelihood ratio test, aBayes [74]. A phylogenetic network was made in SplitsTree [75] using the NeighborNet network [76].

### Nucleotide diversity

Conventional tools calculating nucleotide diversity directly from the variant call files assume that samples are aligned across the entire reference sequence. But as read coverage across the reference genome was highly uneven between samples (Figure S34), adjustment for this was required. Coverage across the reference genome was thus calculated for each sample using samtools mpileup (v 1.9) [67]. For each comparison between two samples, the nucleotide divergence was calculated as number of detected bi-allelic SNPs per nucleotide with read coverage of at least 5X in both samples.

### Population divergence

To account for the differences in read coverage between samples we used the pixy software [77], which produces unbiased estimates of D_XY_ in the presence of missing data. Population divergence (D_XY_) was calculated from this all-site VCF using pixy version 1.0.4.beta1, either in 10 kb windows or in gene coordinate windows.

### Gene sequence deletions

Exploratory Neighbour-Joining phylogenies were produced with CLUSTALW [78, 79] and visualised with FigTree (https://github.com/rambaut/figtree/) after alignment with mafft [78]. Pacbio reads were aligned using Blasr (v 5.3.2) [80], short Illumina reads using NextGenMap (v0.5.5) [66]. Dotplots were produced with FlexiDot (v1.05) [81].

### Gene families and groups

Exported protein-coding gene sets were compiled from the literature [82–84]. Invasion genes were retrieved from Hu *et al*. (2010) [85]. Gene families were assessed in seven *Plasmodium* genomes (*P. simium, P. vivax* SalI, *P. vivax* P01, *P. vivax-like* Pvl01, *P. cynomolgi* M, *P. cynomolgi* B, and *P. knowlesi* H) using the following pipeline: for all genomes, annotated genes were collected for each gene family. These ‘seed’ sequences were used to search all proteins from all genomes using BLASTP and best hits for all proteins were recorded. For each gene family ‘seed’ sequences were then aligned with mafft [63], trimmed with trimAl [64], and HMM models were then built using HMMer (http://hmmer.org/).

For PIR/VIR and PHIST genes, models were built for each genome independently, for all other gene families a single model was built from all genomes. These models were then used to search all proteins in all genomes. All proteins with best BLASTP hit to a ‘seed’ sequence from a given genome were sorted according to their bit score. The lowest 5% of hits were discarded and remaining proteins with best hits to a ‘seed’ sequence were assigned one ‘significant’ hit. As all proteins were searched against ‘seeds’ from the six annotated genomes (*P. simium* excluded), a maximum of six ‘significant’ BLAST hits could be obtained. Similarly, for each HMM model the bottom 25% hits were discarded and remaining hits were considered ‘significant’. The final set of gene families consisted of previously annotated genes and un-annotated genes with at least two ‘significant’ hits (either BLASTP or HMM).

### PCR amplification of DBP1 and RBP2a genes

PCR primers were initially designed from alignments between *P. vivax* and *P. simium* sequences and tested using Primer-BLAST [86] and PlasmoDB (www.plasmodb.org). For DBP1, the reaction was performed in 10 μL volumes containing 0.5 μM of each oligonucleotide primer, 1 μL DNA and 5 μL of Master Mix 2x (Promega) (0.3 units of Taq Polymerase, 200 μM each deoxyribonucleotide triphosphates and 1.5 mM MgCl2). Samples were run with the following settings: 2 minutes of activation at 95°C, followed by 35 cycles with 30 seconds denaturation at 95°C, 30 seconds annealing at 57°C (ΔT=−0.2 °C from 2nd cycle) and 1 minute extension at 72°C, then 5 minutes final extension at 72°C.

For the RBP2a PCR, the reaction was performed in 10 μL volumes containing 0.5 μM of each oligonucleotide primer, 1 μL DNA, 0.1 μL PlatinumTaq DNA Polymerase High Fidelity (Invitrogen, 5U/μL), 0.2 mM each deoxyribonucleotide triphosphates and 2 mM MgSO_4_. The PCR assays were performed with the following cycling parameters: an initial denaturation at 94°C for 1.5 min followed by 40 cycles of denaturation at 94°C for 15 sec, annealing at 65°C for 30 sec (ΔT=−0.2 °C from 2nd cycle) and extension at 68°C for 3.5 min. All Genotyping assays were performed in the thermocycler Veriti 96 wells, Applied Biosystems, and the amplified fragments were visualized by electrophoresis on agarose gels (2% for DBP1 and 1% for RBP2a) in 1x TAE buffer (40 mM Tris-acetate, 1 mM EDTA) with 5 μg/ mL ethidium bromide (Invitrogen) in a horizontal system (Bio-Rad) at 100 V for 30 min. Gels were examined with a UV transilluminator (UVP - Bio-Doc System).

To prevent cross-contamination, the DNA extraction and mix preparation were performed in “parasite DNA-free rooms” distinct from each other. Furthermore, each of these separate areas has different sets of pipettes and all procedures were performed using filtered pipette tips. DNA extraction was performed twice on different days. Positive (DNA extracted from blood from patients with known *P. vivax* infection) and negative (no DNA and DNA extracted from individuals who have never traveled to malaria-endemic areas) controls were used in each round of amplification. DNA extracted from blood of a patient with high parasitemia for *P*. *vivax* and DNA of *P*. *simium* of a non-human primate with an acute infection and parasitemia confirmed by optical microscopy served as positive controls in the PCR assays. Primer sequences are provided in Figure S24 and S29.

### Structural modelling of DBP1 and RBP2a genes

RaptorX [87] was used for prediction of secondary structure and protein disorder. Homology models for the DBL domain of *P. vivax* P01 strain, *P. simium* AF22, and the previously published CDC *P. simium* strain were produced by SWISS-MODEL [88], using the crystallographic structure of the DBL domain from *Plasmodium vivax* DBP bound to the ectodomain of the human DARC receptor (PDB ID 4nuv), with an identity of 98%, 96% and 96% for *P. vivax, P. simium* AF22 and *P. simium* CDC, respectively. QMEAN values were −2.27, −2.04 and −2.03, respectively. The homology model for the reticulocyte binding protein 2 (RBP2a) of *P. vivax* strain P01 was produced based on the cryoEM structure of the complex between the *P. vivax* RBP2b and the human transferrin receptor TfR1 (PDB ID 6d05)[33], with an identity of 31% and QMEAN value of −2.46. The visualization and structural analysis of the produced models was done with PyMOL (https://pymol.org/2/).

## Declarations

### Ethics approval and consent to participate

This study was conducted under the approved project by the IRB commitees in Fiocruz (CAAE Fiocruz: 88554718.0.0000.5262) and in KAUST (19IBEC12).

### Consent for publication

Not applicable

### Availability of data and materials

The reference genome assembly and short sequence reads have been uploaded to European Nucleotide Archive (https://www.ebi.ac.uk/ena/) under the Study accession number PRJEB34061.

### Authors’ contributions

CTDR, PB, AdPC, RLdO, RC and AP conceived the study. CTDR, PB, CFAdB, MdFFdC, RC and AP supervised and co-ordinated the study. AdPC, FVSdA, DAMA, CBJ, JCdSJ and ZMBH collected materials. OD, QG, AdPC, CFAdB, MdFFdC, FVSdA, DAMA, CBJ, JCsSJ and ZMBH conducted wet-lab experiments. TM, AK, SF, DCJ, FJGV, SA, CTDR, PB, RLdO, CFAdB, MdFFdC, FVSdA, DAMA, CBJ, JCdSJ, ZMBH, RC and AP analysed and interpreted the data. TM, RC, AP, CTDR, PB, CFAdB and RLdO drafted and edited the manuscript. All authors read and approved the final manuscript.

## Acknowledgements

We thank Prof. Xin-zhuan Su at the National Institute of Allergy and Infectious Diseases, NIH, for invaluable help in obtaining the parasite gDNA from the BEI Resources; Sidnei Silva and Graziela Zanini, for assistance on the parasitological diagnosis of the human samples; Aline Lavigne and Larissa Gomes for undertaking the PCR for *P vivax;* Alcides Pissinatti and Silvia Bahadian Moreira for the facilities provided at the Primate Centre of Rio de Janeiro; Orzinete Rodrigues Soares for non-human primates’ blood slides’; Marcelo Quintela, Waldemir Paixão Vargas, Carlos Alberto C. da Silva, Alexandre B. de Souza, Vicente Klonowski, Romenique L. Araújo, Luis R. Nogueira, Fernando Barreto, Ana L. Quijada, Luiz P.P. Silva, Gelson Medeiros, Adilson B. Ramos, Marcilene B. Ramos, Carlos A.A. Júnior, Paulo G. Barbosa, Sérgio F. Fragoso, Adilson R. Silva, Cecília Cronemberger, Marcelo Rheingantz, Leonardo Nascimento and João Marins for the field support; *Grupo Técnico de Vigilância de Arboviroses* (GT-Arbo – Brazilian Ministry of Health) for field and material supports; and Cassio Leonel Peterka from The Brazilian Ministry of Health for malaria epidemiological data. The following reagent was obtained through BEI Resources, NIAID, NIH: *Plasmodium simium*, Strain Howler, MRA-353, contributed by William E. Collins. We thank Kieran Samuk for assistance with pixy, Richard Carter for invaluable discussion and comment on previous versions of this manuscript, and Tony Holder for critical reading of the manuscript.

## References

1. Brasil P, Zalis MG, de Pina-Costa A, Siqueira AM, Junior CB, Silva S, Areas ALL, Pelajo-Machado M, de Alvarenga DAM, da Silva Santelli ACF et al:Outbreak of human malaria caused by Plasmodium simium in the Atlantic Forest in Rio de Janeiro: a molecular epidemiological investigation. Lancet Glob Health 2017, 5(10):e1038–e1046.

2. Cox-Singh J, Davis TME, Lee K-S, Shamsul SSG, Matusop A, Ratnam S, Rahman HA, Conway DJ, Singh B: Plasmodium knowlesi malaria in humans is widely distributed and potentially life threatening. Clinical infectious diseases: an official publication of the Infectious Diseases Society of America 2008, 46(2): 165–171.

3. Imwong M, Madmanee W, Suwannasin K, Kunasol C, Peto TJ, Tripura R, von Seidlein L, Nguon C, Davoeung C, Day NPJ et al: Asymptomatic Natural Human Infections With the Simian Malaria Parasites Plasmodium cynomolgi and Plasmodium knowlesi. J Infect Dis 2019, 219(5):695–702.

4. Loy DE, Liu W, Li Y, Learn GH, Plenderleith LJ, Sundararaman SA, Sharp PM, Hahn BH: Out of Africa: origins and evolution of the human malaria parasites Plasmodium falciparum and Plasmodium vivax. Int J Parasitol 2017, 47(2-3):87–97.

5. Liu W, Li Y, Learn GH, Rudicell RS, Robertson JD, Keele BF, Ndjango JB, Sanz CM, Morgan DB, Locatelli S et al:Origin of the human malaria parasite Plasmodium falciparum in gorillas. Nature 2010, 467(7314):420–425.

6. Liu W, Li Y, Shaw KS, Learn GH, Plenderleith LJ, Malenke JA, Sundararaman SA, Ramirez MA, Crystal PA, Smith AG et al:African origin of the malaria parasite Plasmodium vivax. Nature communications 2014, 5:3346.

7. Marques GR, Condino ML, Serpa LL, Cursino TV: [Epidemiological aspects of autochthonous malaria in the Atlantic forest area of the northern coast of the State of Sao Paulo, 1985-2006]. Rev Soc Bras Med Trop 2008, 41(4):386–389.

8. Cox-Singh J, Davis TM, Lee KS, Shamsul SS, Matusop A, Ratnam S, Rahman HA, Conway DJ, Singh B: Plasmodium knowlesi malaria in humans is widely distributed and potentially life threatening. Clin Infect Dis 2008, 46(2):165–171.

9. Ta TH, Hisam S, Lanza M, Jiram AI, Ismail N, Rubio JM: First case of a naturally acquired human infection with Plasmodium cynomolgi. Malar J 2014, 13:68.

10. Marchand RP, Culleton R, Maeno Y, Quang NT, Nakazawa S: Co-infections of Plasmodium knowlesi, P. falciparum, and P. vivax among Humans and Anopheles dirus Mosquitoes, Southern Vietnam. Emerg Infect Dis 2011, 17(7):1232–1239.

11. Otto TD, Gilabert A, Crellen T, Bohme U, Arnathau C, Sanders M, Oyola SO, Okouga AP, Boundenga L, Willaume E et al: Genomes of all known members of a Plasmodium subgenus reveal paths to virulent human malaria. Nat Microbiol 2018, 3(6):687–697.

12. de Pina-Costa A, Brasil P, Di Santi SM, de Araujo MP, Suarez-Mutis MC, Santelli AC, Oliveira-Ferreira J, Lourenco-de-Oliveira R, Daniel-Ribeiro CT: Malaria in Brazil: what happens outside the Amazonian endemic region. Mem Inst Oswaldo Cruz 2014, 109(5):618–633.

13. Collins WE, Contacos PG, Guinn EG: Observations on the sporogonic cycle and transmission of Plasmodium simium Da Fonseca. J Parasitol 1969, 55(4):814–816.

14. de Alvarenga DAM, Culleton R, de Pina-Costa A, Rodrigues DF, Bianco C, Jr., Silva S, Nunes AJD, de Souza JC, Jr., Hirano ZMB, Moreira SB et al:An assay for the identification of Plasmodium simium infection for diagnosis of zoonotic malaria in the Brazilian Atlantic Forest. Scientific reports 2018, 8(1):86.

15. Escalante AA, Cornejo OE, Freeland DE, Poe AC, Durrego E, Collins WE, Lal AA: A monkey’s tale: the origin of Plasmodium vivax as a human malaria parasite. Proc Natl Acad Sci USA 2005, 102(6):1980–1985.

16. Gelabert P, Sandoval-Velasco M, Olalde I, Fregel R, Rieux A, Escosa R, Aranda C, Paaijmans K, Mueller I, Gilbert MT et al: Mitochondrial DNA from the eradicated European Plasmodium vivax and P. falciparum from 70-year-old slides from the Ebro Delta in Spain. Proceedings of the National Academy of Sciences of the United States of America 2016, 113(41): 11495–11500.

17. Rodrigues PT, Valdivia HO, de Oliveira TC, Alves JMP, Duarte A, Cerutti-Junior C, Buery JC, Brito CFA, de Souza JC, Jr., Hirano ZMB et al:Human migration and the spread of malaria parasites to the New World. Scientific reports 2018, 8(1):1993.

18. Deane LM: Simian malaria in Brazil. Mem Inst Oswaldo Cruz 1992, 87 Suppl 3:1–20.

19. Fonseca F: [Plasmodium of a primate of Brazil]. Mem Inst Oswaldo Cruz 1951, 49:543–553.

20. Leclerc MC, Durand P, Gauthier C, Patot S, Billotte N, Menegon M, Severini C, Ayala FJ, Renaud F: Meager genetic variability of the human malaria agent Plasmodium vivax. Proceedings of the National Academy of Sciences of the United States of America 2004, 101(40):14455–14460.

21. Duarte AM, Porto MA, Curado I, Malafronte RS, Hoffmann EH, de Oliveira SG, da Silva AM, Kloetzel JK, Gomes Ade C: Widespread occurrence of antibodies against circumsporozoite protein and against blood forms of Plasmodium vivax, P. falciparum and P. malariae in Brazilian wild monkeys. J Med Primatol 2006, 35(2):87–96.

22. Tazi L, Ayala FJ: Unresolved direction of host transfer of Plasmodium vivax v. P. simium and P. malariae v. P. brasilianum. Infect Genet Evol 2011, 11(1):209–221.

23. Coatney GR, Collins WE, Warren M, Contacos PG: Plasmodium simium. In: The Primate Malarias. National Institutes of Health; 1971.

24. Auburn S, Bohme U, Steinbiss S, Trimarsanto H, Hostetler J, Sanders M, Gao Q, Nosten F, Newbold CI, Berriman M et al: A new Plasmodium vivax reference sequence with improved assembly of the subtelomeres reveals an abundance of pir genes. Wellcome Open Res 2016, 1:4.

25. Hupalo DN, Luo Z, Melnikov A, Sutton PL, Rogov P, Escalante A, Vallejo AF, Herrera S, Arevalo-Herrera M, Fan Q et al: Population genomics studies identify signatures of global dispersal and drug resistance in Plasmodium vivax. Nat Genet 2016, 48(8):953–958.

26. Alexander DH, Novembre J, Lange K: Fast model-based estimation of ancestry in unrelated individuals. Genome research 2009, 19(9):1655–1664.

27. Cruickshank TE, Hahn MW: Reanalysis suggests that genomic islands of speciation are due to reduced diversity, not reduced gene flow. Mol Ecol 2014, 23(13):3133–3157.

28. Nei M, Li WH: Mathematical model for studying genetic variation in terms of restriction endonucleases. Proceedings of the National Academy of Sciences of the United States of America 1979, 76(10):5269–5273.

29. Kanjee U, Rangel GW, Clark MA, Duraisingh MT: Molecular and cellular interactions defining the tropism of Plasmodium vivax for reticulocytes. Curr Opin Microbiol 2018, 46:109–115.

30. Miller LH, McAuliffe FM, Mason SJ: Erythrocyte receptors for malaria merozoites. Am J Trop Med Hyg 1977, 26(6 Pt 2):204–208.

31. Iyer J, Gruner AC, Renia L, Snounou G, Preiser PR: Invasion of host cells by malaria parasites: a tale of two protein families. Mol Microbiol 2007, 65(2):231–249.

32. Chan LJ, Dietrich MH, Nguitragool W, Tham WH: Plasmodium vivax Reticulocyte Binding Proteins for invasion into reticulocytes. Cell Microbiol 2019:e13110.

33. Gruszczyk J, Huang RK, Chan LJ, Menant S, Hong C, Murphy JM, Mok YF, Griffin MDW, Pearson RD, Wong W et al: Cryo-EM structure of an essential Plasmodium vivax invasion complex. Nature 2018, 559(7712):135–139.

34. Carlton JM, Adams JH, Silva JC, Bidwell SL, Lorenzi H, Caler E, Crabtree J, Angiuoli SV, Merino EF, Amedeo P et al: Comparative genomics of the neglected human malaria parasite Plasmodium vivax. Nature 2008, 455(7214):757–763.

35. Gilabert A, Otto TD, Rutledge GG, Franzon B, Ollomo B, Arnathau C, Durand P, Moukodoum ND, Okouga AP, Ngoubangoye B et al: Plasmodium vivax-like genome sequences shed new insights into Plasmodium vivax biology and evolution. PLoS Biol 2018, 16(8):e2006035.

36. Ntumngia FB, McHenry AM, Barnwell JW, Cole-Tobian J, King CL, Adams JH: Genetic variation among Plasmodium vivax isolates adapted to non-human primates and the implication for vaccine development. Am J Trop Med Hyg 2009, 80(2):218–227.

37. Rausch T, Zichner T, Schlattl A, Stutz AM, Benes V, Korbel JO: DELLY: structural variant discovery by integrated paired-end and split-read analysis. Bioinformatics 2012, 28(18):i333–i339.

38. Batchelor JD, Zahm JA, Tolia NH: Dimerization of Plasmodium vivax DBP is induced upon receptor binding and drives recognition of DARC. Nat Struct Mol Biol 2011, 18(8):908–914.

39. de Oliveira TC, Rodrigues PT, Early AM, Duarte A, Buery JC, Bueno MG, Catao-Dias JL, Cerutti C, Jr., Rona LDP, Neafsey DE et al:Plasmodium simium: population genomics reveals the origin of a reverse zoonosis. J Infect Dis 2021.

40. Meneguzzi VC, Santos CB, Pinto Ide S, Feitoza LR, Feitoza HN, Falqueto A: Use of geoprocessing to define malaria risk areas and evaluation of the vectorial importance of anopheline mosquitoes (Diptera: Culicidae) in Espirito Santo, Brazil. Mem Inst Oswaldo Cruz 2009, 104(4):570–575.

41. Proto WR, Siegel SV, Dankwa S, Liu W, Kemp A, Marsden S, Zenonos ZA, Unwin S, Sharp PM, Wright GJ et al: Adaptation of Plasmodium falciparum to humans involved the loss of an ape-specific erythrocyte invasion ligand. Nature communications 2019, 10(1):4512.

42. Wright PE, Dyson HJ: Intrinsically disordered proteins in cellular signalling and regulation. Nature reviews Molecular cell biology 2015, 16(1): 18–29.

43. Culleton R, Carter R: African Plasmodium vivax: distribution and origins. Int J Parasitol 2012, 42(12):1091–1097.

44. Li J, Collins WE, Wirtz RA, Rathore D, Lal A, McCutchan TF: Geographic subdivision of the range of the malaria parasite Plasmodium vivax. Emerging infectious diseases 2001, 7(1):35–42.

45. Choi EH, Lee SK, Ihm C, Sohn YH: Rapid DNA extraction from dried blood spots on filter paper: potential applications in biobanking. Osong Public Health Res Perspect 2014, 5(6):351–357.

46. Bolger AM, Lohse M, Usadel B: Trimmomatic: a flexible trimmer for Illumina sequence data. Bioinformatics 2014, 30(15):2114–2120.

47. Langmead B, Salzberg SL: Fast gapped-read alignment with Bowtie 2. Nature methods 2012, 9(4):357–359.

48. Bankevich A, Nurk S, Antipov D, Gurevich AA, Dvorkin M, Kulikov AS, Lesin VM, Nikolenko SI, Pham S, Prjibelski AD et al:SPAdes: a new genome assembly algorithm and its applications to single-cell sequencing. J Comput Biol 2012, 19(5):455–477.

49. Boetzer M, Henkel CV, Jansen HJ, Butler D, Pirovano W: Scaffolding pre-assembled contigs using SSPACE. Bioinformatics 2011, 27(4):578–579.

50. Boetzer M, Pirovano W: Toward almost closed genomes with GapFiller. Genome biology 2012, 13(6):R56.

51. Walker BJ, Abeel T, Shea T, Priest M, Abouelliel A, Sakthikumar S, Cuomo CA, Zeng Q, Wortman J, Young SK et al:Pilon: an integrated tool for comprehensive microbial variant detection and genome assembly improvement. PloS one 2014, 9(11):e112963.

52. Laetsch D, Blaxter M: BlobTools: Interrogation of genome assemblies [version 1; referees: 2 approved with reservations]. F1000Research 2017, 6(1287).

53. Jongwutiwes S, Putaporntip C, Iwasaki T, Ferreira MU, Kanbara H, Hughes AL: Mitochondrial genome sequences support ancient population expansion in Plasmodium vivax. Molecular biology and evolution 2005, 22(8):1733–1739.

54. Rodrigues PT, Alves JM, Santamaria AM, Calzada JE, Xayavong M, Parise M, da Silva AJ, Ferreira MU: Using mitochondrial genome sequences to track the origin of imported Plasmodium vivax infections diagnosed in the United States. Am J Trop Med Hyg 2014, 90(6):1102–1108.

55. Bandelt HJ, Forster P, Rohl A: Median-joining networks for inferring intraspecific phylogenies. Molecular biology and evolution 1999, 16(1):37–48.

56. Leigh JW, Bryant D: POPART: full-feature software for haplotype network construction. Methods Ecol Evol 2015, 6(9): 1110–1116.

57. Holt C, Yandell M: MAKER2: an annotation pipeline and genome-database management tool for second-generation genome projects. BMC Bioinformatics 2011, 12(1):491.

58. Steinbiss S, Silva-Franco F, Brunk B, Foth B, Hertz-Fowler C, Berriman M, Otto TD: Companion: a web server for annotation and analysis of parasite genomes. Nucleic acids research 2016, 44(W1):W29–34.

59. Waterhouse RM, Seppey M, Simao FA, Manni M, Ioannidis P, Klioutchnikov G, Kriventseva EV, Zdobnov EM: BUSCO applications from quality assessments to gene prediction and phylogenomics. Molecular biology and evolution 2017.

60. Simao FA, Waterhouse RM, Ioannidis P, Kriventseva EV, Zdobnov EM: BUSCO: assessing genome assembly and annotation completeness with single-copy orthologs. Bioinformatics 2015, 31(19):3210–3212.

61. Aurrecoechea C, Brestelli J, Brunk BP, Dommer J, Fischer S, Gajria B, Gao X, Gingle A, Grant G, Harb OS et al: PlasmoDB: a functional genomic database for malaria parasites. Nucleic acids research 2009, 37(Database issue):D539–543.

62. Lechner M, Findeiss S, Steiner L, Marz M, Stadler PF, Prohaska SJ: Proteinortho: detection of (co-)orthologs in large-scale analysis. BMC Bioinformatics 2011, 12:124.

63. Katoh K, Standley DM: MAFFT multiple sequence alignment software version 7: improvements in performance and usability. Molecular biology and evolution 2013, 30(4):772–780.

64. Capella-Gutierrez S, Silla-Martinez JM, Gabaldon T: trimAl: a tool for automated alignment trimming in large-scale phylogenetic analyses. Bioinformatics 2009, 25(15):1972–1973.

65. Stamatakis A: RAxML version 8: a tool for phylogenetic analysis and post-analysis of large phylogenies. Bioinformatics 2014, 30(9):1312–1313.

66. Sedlazeck FJ, Rescheneder P, von Haeseler A: NextGenMap: fast and accurate read mapping in highly polymorphic genomes. Bioinformatics 2013, 29(21):2790–2791.

67. Li H, Handsaker B, Wysoker A, Fennell T, Ruan J, Homer N, Marth G, Abecasis G, Durbin R: The Sequence Alignment/Map format and SAMtools. Bioinformatics 2009, 25(16):2078–2079.

68. McKenna A, Hanna M, Banks E, Sivachenko A, Cibulskis K, Kernytsky A, Garimella K, Altshuler D, Gabriel S, Daly M et al: The Genome Analysis Toolkit: a MapReduce framework for analyzing next-generation DNA sequencing data. Genome research 2010, 20(9):1297–1303.

69. Garrison E, Marth G: Haplotype-based variant detection from short-read sequencing. arXiv 2012:1207.3907.

70. Chang CC, Chow CC, Tellier LC, Vattikuti S, Purcell SM, Lee JJ: Second-generation PLINK: rising to the challenge of larger and richer datasets. Gigascience 2015, 4:7.

71. Guindon S, Dufayard JF, Lefort V, Anisimova M, Hordijk W, Gascuel O: New algorithms and methods to estimate maximum-likelihood phylogenies: assessing the performance of PhyML 3.0. Syst Biol 2010, 59(3):307–321.

72. Guindon S, Gascuel O: A simple, fast, and accurate algorithm to estimate large phylogenies by maximum likelihood. Syst Biol 2003, 52(5):696–704.

73. Lefort V, Longueville JE, Gascuel O: SMS: Smart Model Selection in PhyML. Molecular biology and evolution 2017, 34(9):2422–2424.

74. Anisimova M, Gil M, Dufayard JF, Dessimoz C, Gascuel O: Survey of branch support methods demonstrates accuracy, power, and robustness of fast likelihood-based approximation schemes. Syst Biol 2011, 60(5):685–699.

75. Huson DH, Bryant D: Application of phylogenetic networks in evolutionary studies. Molecular biology and evolution 2006, 23(2):254–267.

76. Bryant D, Moulton V: Neighbor-net: an agglomerative method for the construction of phylogenetic networks. Molecular biology and evolution 2004, 21(2):255–265.

77. Korunes KL, Samuk K: pixy: Unbiased estimation of nucleotide diversity and divergence in the presence of missing data. Mol Ecol Resour 2021, 21(4): 1359–1368.

78. Larkin MA, Blackshields G, Brown NP, Chenna R, McGettigan PA, McWilliam H, Valentin F, Wallace IM, Wilm A, Lopez R et al: Clustal W and Clustal X version 2.0. Bioinformatics 2007, 23(21):2947–2948.

79. Thompson JD, Higgins DG, Gibson TJ: CLUSTAL W: improving the sensitivity of progressive multiple sequence alignment through sequence weighting, position-specific gap penalties and weight matrix choice. Nucleic acids research 1994, 22(22):4673–4680.

80. Chaisson MJ, Tesler G: Mapping single molecule sequencing reads using basic local alignment with successive refinement (BLASR): application and theory. BMC Bioinformatics 2012, 13:238.

81. Seibt KM, Schmidt T, Heitkam T: FlexiDot: highly customizable, ambiguity-aware dotplots for visual sequence analyses. Bioinformatics 2018, 34(20):3575–3577.

82. van Ooij C, Tamez P, Bhattacharjee S, Hiller NL, Harrison T, Liolios K, Kooij T, Ramesar J, Balu B, Adams J et al: The malaria secretome: from algorithms to essential function in blood stage infection. PLoS pathogens 2008, 4(6):e1000084.

83. Boddey JA, Carvalho TG, Hodder AN, Sargeant TJ, Sleebs BE, Marapana D, Lopaticki S, Nebl T, Cowman AF: Role of plasmepsin V in export of diverse protein families from the Plasmodium falciparum exportome. Traffic 2013, 14(5):532–550.

84. Schulze J, Kwiatkowski M, Borner J, Schluter H, Bruchhaus I, Burmester T, Spielmann T, Pick C: The Plasmodium falciparum exportome contains non-canonical PEXEL/HT proteins. Mol Microbiol 2015, 97(2):301–314.

85. Hu G, Cabrera A, Kono M, Mok S, Chaal BK, Haase S, Engelberg K, Cheemadan S, Spielmann T, Preiser PR et al: Transcriptional profiling of growth perturbations of the human malaria parasite Plasmodium falciparum. Nat Biotechnol 2010, 28(1):91–98.

86. Ye J, Coulouris G, Zaretskaya I, Cutcutache I, Rozen S, Madden TL: Primer-BLAST: a tool to design target-specific primers for polymerase chain reaction. BMC Bioinformatics 2012, 13:134.

87. Wang S, Li W, Liu S, Xu J: RaptorX-Property: a web server for protein structure property prediction. Nucleic acids research 2016, 44(W1):W430–435.

88. Waterhouse A, Bertoni M, Bienert S, Studer G, Tauriello G, Gumienny R, Heer FT, de Beer TAP, Rempfer C, Bordoli L et al: SWISS-MODEL: homology modelling of protein structures and complexes. Nucleic acids research 2018, 46(W1):W296–W303.

