## Supplementary Figures for "The genome of the zoonotic malaria parasite *Plasmodium simium* reveals adaptations to host-switching"

Supplementary Figure Legends

**Figure S1 - Apicoplast and mitochondrial genomes**

Circos plots of the assembled apicoplast (top) and mitochondrial (bottom) genomes. The presence of protein-coding genes (grey), ribosomal-RNAs (green), and transfer-RNAs (red) is denoted by bars. For each gene category, outermost circle denotes genes on the forward strand, and the innermost circle denotes genes on the reverse strand. The two circular representations nearest to the center represent GC-skew and GC-content, respectively.

**Figure S2 – Busco assembly assessment**

Gene content in *Plasmodium simium* and other *Plasmodium* genome assemblies. BUSCO was run using the eukaryota odb9 data set containing 303 BUSCO groups.

**Figure S3 - Protein phylogeny**

Maximum likelihood phylogeny based on 3,204 concatenated *Plasmodium simium* protein-coding genes with 1:1 orthologs across a selection of *Plasmodium* species (see Materials and Methods). Tree was constructed using RaxML with the GTRGAMMA model and *Plasmodium gallinaceum* as outgroup. Branch support from 100 bootstrap replicates.

**Figure S4 – Gene families**

The number of gene family members among *Plasmodium* genomes. Diameters of circles are proportional to the $\log_{10}$-transformed number of family members, as shown on the right.

**Figure S5 – Pir/Vir gene clustering**

Clustering of PIR protein sequences based on BLASTP similarity. Network threshold is a bit-score of 40 and visualized using the edge-weighted spring embedded layout in cytoscape [3].

**Figure S6 - SNP counts**

Top: Number of high-confidence SNPs detected in each sample. *Plasmodium simium* samples shown in dark red, and *Plasmodium vivax* samples in orange (see also

Supplementary Tables S3 & S4). Bottom: For a given SNP loci data is not available from all samples. The bar chart shows how many SNP loci (left y-axis) that have data (coverage) in a given number of samples (x-axis). The cumulative fraction of SNPs is shown as red line (right x-axis).

**Figure S7 - PCA plot**

A) Plot of the first two dimensions from a principal component analysis of 124,968 SNPs. *Plasmodium vivax* samples are denoted by their geographic origin. B) Magnification of American vivax samples. C) Cumulative contribution (in percent) of each eigenvector. The separate clustering of *Plasmodium simium* and American *P. vivax* samples was also observed when plotting 2nd versus 3rd dimensions, and 3rd versus 4th dimensions (not shown).

**Figure S8 - MDS plot**

Plot of Multidimensional Scaling of *Plasmodium simium* and *Plasmodium vivax* samples.

**Figure S9 - SNP phylogeny**

Phylogenetic tree constructed from SNP sites. Identical to Figure 1, but with sample IDs shown at terminal branches.

**Figure S10 - Phylogenetic network**

Network from nucleotide diversity distances produced in SplitsTree [1] using the NeighborNet network [2]. A magnification of the central hubs (red rectangle) is shown at the bottom right.

**Figure S11 - Admixture**

Q-estimates from unsupervised ADMIXTURE clustering analysis at K from 2 to 10. *Plasmodium vivax* samples are ordered by geographic origin. The lowest cross-validation error was observed for K=3.

**Figure S12 - Gene $D_{XY}$ diversity**

A) For different intervals of average $D_{XY}$ values (x-axis), the number of genes with thee values are plotted (y-axis, $log_{10}$-scaled).

B) Box plot showing the distributions of $D_{XY}$ values for members of selected gene families. The right-most, yellow box shows the DXY values for all genes that not member of a gene family.

**Figure S13 - DBP phylogeny**

Neighbor-Joining tree of *Plasmodium* DBP protein sequences. Sequences were aligned using mafft and tree produced with CLUSTALW. Support from 1000 bootstrap replicates. Tree visualized with FigTree, genetic distance shown below tree. *Plasmodium simium* and *Plasmodium vivax* sequences derived from this study are highlighted in red and blue, respectively. Remaining sequences are suffixed by their genome of origin (PvivP; *P. vivax* P01, PvivS; *P. vivax* SalI, PcynM; *P. cynomolgi* M, PcynB; *P. cynomolgi* B, PknoH; *P. knowlesi* H).

**Figure S14 - RBP phylogeny**

Neighbor-Joining tree of Plasmodium RBP protein sequences. Sequences were aligned using mafft and tree produced with CLUSTALW. Support from 1000 bootstrap replicates. Tree visualized with FigTree, genetic distance shown below tree. *Plasmodium simium* and *Plasmodium vivax* sequences derived from this study are highlighted in red and blue, respectively. Remaining sequences are suffixed by their genome of origin (PvivP; *P. vivax* P01, PvivS; *P. vivax* SalI, PcynM; *P. cynomolgi* M, PcynB; *P. cynomolgi* B, PknoH; *P. knowlesi* H).

**Figure S15 - Coverage across RBP gene loci**

Average read coverage across gene loci when mapping human *Plasmodium simium* (top) and *Plasmodium vivax* (bottom) reads onto the *P. vivax* P01 genome. The RBPs genes are shown above plots with the *P. vivax* P01 gene identifiers below plots. Genes absent in *P. simium* is highlighted in grey. Note that the *P. vivax* P01 genome contains two annotated RBP1a and RBP2d genes, respectively. The average read coverage per gene is shown as dots for each individual sample, and the combined distributions are outlined by boxes. A single *P. simium* sample, AF22, has a typical coverage of around 200X but is omitted from this representation for clarity. The coverage of the *P. simium* CDC strain is highlighted as red dots.

**Figure S16 – coverage across flanking regions**

The average read coverage across *Plasmodium vivax* DBP and RBP genes is compared to the average read coverage across flanking genomic regions. The $\log_2$ ratio between coverage at flanking region and coverage at gene was calculated for four regions: 10kb-5kb upstream of gene (UP2), 5kb-0kb upstream of gene (UP1), 0kb-5kb downstream of gene (DOWN1), 5kb-10kb downstream of gene (DOWN2). Boxplot denotes the range of ratios for *Plasmodium simium* samples (red boxes), American *Plasmodium vivax* samples (light blue), and remaining *Plasmodium vivax* samples (blue). Up- and downstream are defined based on genome coordinates irrespective of gene orientation. After Bonferroni correction, only the 'UP1' region at RBP3 for *P. simium* samples (p=0.011, denoted by asterisk) had a probability below 0.05 of the mean being above zero assuming a normal distribution.

**Figure S17 - DBP1 alignment**

Complete alignment of DBP1 protein sequences. The presence of the DBL domain (Pfam: PF0311) and the trans-membrane domain is indicated.

**Figure S18 - Haplotype network of DBP1 sequences**

Haplotype network (minimum spanning network) produced using PopART [4,5]. Numbers of mutations are indicated by hatch marks on edges. *Plasmodium simium* samples are shown in red, *Plasmodium vivax* in green, and *P. vivax-like* in purple. The previously published *P. simium* CDC strain sequence (ACB42432) is shown in black and bold.

**Figure S19 - Read support for DBP1 deletion patterns**

Among *Plasmodium vivax* and *Plasmodium simium* samples multiple deletion patterns in the DBP1 gene are observed (top). Deletions found in *P. vivax* samples are arbitrarily denoted vivax 1-4. For each sample, the number of reads supporting a given deletion is shown (bottom). Samples with reads supporting multiple deletion forms are indicated by asterisks.

**Figure S20 - Dotplot**

Similarity DNA dot plots of DBP1 genes. Top plots show *Plasmodium simium* and bottom plots show *Plasmodium vivax* DBP1 genes. Introns are denoted by grey rectangles and the deleted region (or site of deleted region in *P. simium*) is highlighted in red.

**Figure S21 - Read coverage across the DBP1 deletion in *Plasmodium simium* samples**

Artemis representation of read coverage across the *Plasmodium simium* DBP1 deletion. Reads from *Plasmodium vivax* samples AM01 & AM02 (two top panels) and *P. simium* AF22 & AF36 samples (two bottom panels) were mapped onto the *P. simium* AF22 assembly. The site of deletion is indicated by vertical red arrows. Note the lack of *P. vivax* reads spanning the deletion site.

**Figure S22 - read coverage across the DBP1 deletion in *Plasmodium vivax* samples**

Artemis representation of read coverage across the *Plasmodium simium* DBP1 deletion. Reads from *P. simium* AF22 & AF36 samples were mapped onto the *Plasmodium vivax* P01 genome. The site of deletion is indicated by red squares. Note the lack of *P. simium* coverage across the deleted region.

**Figure S23 - PacBio read coverage across the DBP1 deletion in *Plasmodium simium***

Schematic depiction of PacBio reads mapping across the deletion in DBP1 gene. X-axis denotes genomic positions. Black line at the center of plot shows the extent of exons. Reads mapping on the forward strand are shown above exons, reverse strand reads underneath exons. Reads are colored according to the sample from which they are derived. The site of deletion is indicated by arrows and thin vertical orange lines.

**Figure S24 – DBP1 PCR**

Top: Schematic overview of DBP1 deletion PCR approach. Gel images shown for human *Plasmodium vivax* samples (top gel image), human *Plasmodium simium* samples (middle image), and non-human primate (NHP) *P. simium* samples (bottom image). Expected band sizes with and without deletion event are indicated by red triangles. Bottom: Primer sequences.

**Figure S25 - RBP2a alignment**

Complete alignment of RBP2a protein sequences

**Figure S26 - read coverage across the RBP2a deletion in *Plasmodium simium* samples**

Artemis representation of read coverage across the *Plasmodium simium* RBP2a deletion. Reads from vivax samples AM01 & AM02 (two top panels) and s *P. simium* AF22 & AF36 samples (two bottom panels) were mapped onto the *P. simium* AF22 assembly. The site of deletion is indicated by vertical red arrows. Note the lack of *Plasmodium vivax* reads spanning the deletion site.

**Figure S27 - read coverage across the RBP2a deletion in *Plasmodium vivax* samples**

Artemis representation of read coverage across the *Plasmodium simium* RBP2a deletion. Reads from *P. simium* AF22 & AF36 samples were mapped onto the *P. vivax* P01 genome. The site of deletion is indicated by red squares. Note the lack of *P. simium* coverage across the deleted region.

**Figure S28 - PacBio read coverage across RBP2a deletion in *Plasmodium simium***

Schematic depiction of PacBio reads mapping across deletions in the RBP2a gene. X-axis denotes genomic positions. Black line at the center of plots shows the extent of exons. Reads mapping on the forward strand are shown above exons, reverse strand reads underneath exons. Reads are colored according to the sample from which they are derived. Sites of deletions are indicated by arrows and thin vertical orange lines.

**Figure S29 – RBP2a PCR**

Top: Schematic overview of RBP2a deletion PCR approach. Gel images shown for *Plasmodium vivax* and *Plasmodium simium* samples. Expected band sizes with and without deletion event are indicated by red triangles. Bottom: Primer sequences.

**Figure S30 – Short indels**

A) Pie chart showing the percentage of indels being integers of 1-3 base pairs. B) The percentage of insertions (left bar) and deletions (middle) overlapping low-complexity regions in proteins. The percentage of all proteins consisting of low-complexity sequences is shown on the right. Low-complexity annotation downloaded from

PlasmoDB. C) Size distributions of genes with and without indels. Genes with indels are listed in Supplementary Table S7.

**Figure S31 – DBP1 protein structures**

*Plasmodium simium* DBP1 is predicted to associate with human DARC. Left: Top-view of modelled DBP1 from *Plasmodium vivax* strain P01, human-infecting *P. simium* AF22 (PsDBP1) and monkey-infecting *P. simium* (CDC PsDBP1) sequences. Models were established based on the crystal structure of the *P. vivax* DBP1, strain Salvador 1, in complex with human DARC (PDB 4nuv). Individual chains of the DBP1 dimer are shown in light and dark blue. DARC (residues 19-30) is coloured in grey. Residue substitutions between the models and the crystal structure are highlighted in magenta. Right: close-up view of the DARC-binding site, as delimited by a dashed square on the left. Colours as in the left panel. The rearrangement of hydrogen-bonds (black dotted lines) is shown for the Lys-Asn substitution.

**Figure S32 – Mitochondrial haplotype network**

Haplotype network (minimum spanning network) produced using PopART [4,5]. Numbers of mutations are shown on edges. The two identical *Plasmodium simium* samples (AF22 and GenBank accession AY722798) are shown in black. *Plasmodium vivax* haplogroups coloured according to country of origin. Circle sizes indicate the number of sequences in haplogroups.

**Figure S33 – Protein orthology**

Venn diagram showing the number of shared gene orthogroups between *Plasmodium simium* and three other *Plasmodium* genomes.

**Figure S34 - SNP allele frequencies & coverage**

Top: Allele frequencies for SNPs with available data from at least 37 *Plasmodium simium* or *Plasmodium vivax* samples. Bottom: Mean read coverage at SNP sites shown for each samples.

1 Huson, D. H. & Bryant, D. Application of phylogenetic networks in evolutionary studies. *Molecular biology and evolution* **23**, 254-267, doi:10.1093/molbev/msj030 (2006).
2 Bryant, D. & Moulton, V. Neighbor-net: an agglomerative method for the construction of phylogenetic networks. *Molecular biology and evolution* **21**, 255-265, doi:10.1093/molbev/msh018 (2004).
3 Shannon, P. *et al.* Cytoscape: a software environment for integrated models of biomolecular interaction networks. *Genome research* **13**, 2498-2504, doi:10.1101/gr.1239303 (2003).
4 Bandelt, H. J., Forster, P. & Rohl, A. Median-joining networks for inferring intraspecific phylogenies. *Molecular biology and evolution* **16**, 37-48, doi:10.1093/oxfordjournals.molbev.a026036 (1999).
5 Leigh, J. W. & Bryant, D. POPART: full-feature software for haplotype network construction. *Methods Ecol Evol* **6**, 1110-1116, doi:10.1111/2041-210x.12410 (2015).

Figure S1

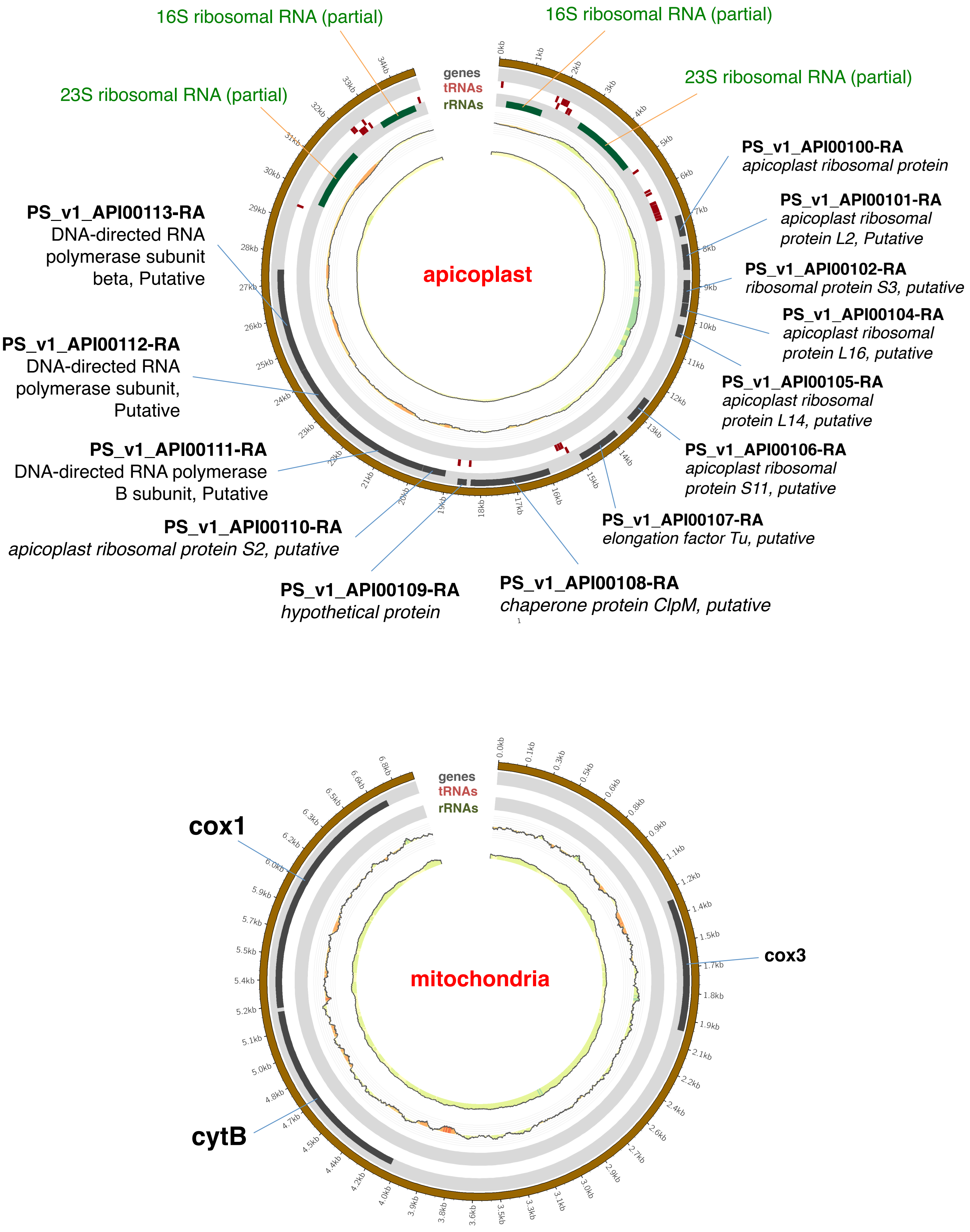
16S ribosomal RNA (partial)
16S ribosomal RNA (partial)
23S ribosomal RNA (partial)
23S ribosomal RNA (partial)
genes
tRNAs
rRNAs
apicoplast
PS_v1_API00100-RA
apicoplast ribosomal protein
PS_v1_API00101-RA
apicoplast ribosomal protein L2, Putative
PS_v1_API00102-RA
ribosomal protein S3, putative
PS_v1_API00104-RA
apicoplast ribosomal protein L16, putative
PS_v1_API00105-RA
apicoplast ribosomal protein L14, putative
PS_v1_API00106-RA
apicoplast ribosomal protein S11, putative
PS_v1_API00107-RA
elongation factor Tu, putative
PS_v1_API00108-RA
chaperone protein ClpM, putative
PS_v1_API00109-RA
hypothetical protein
PS_v1_API00110-RA
apicoplast ribosomal protein S2, putative
PS_v1_API00111-RA
DNA-directed RNA polymerase B subunit, Putative
PS_v1_API00112-RA
DNA-directed RNA polymerase subunit, Putative
PS_v1_API00113-RA
DNA-directed RNA polymerase subunit beta, Putative
genes
tRNAs
rRNAs
mitochondria
cox1
cox3
cytB

Figure S2

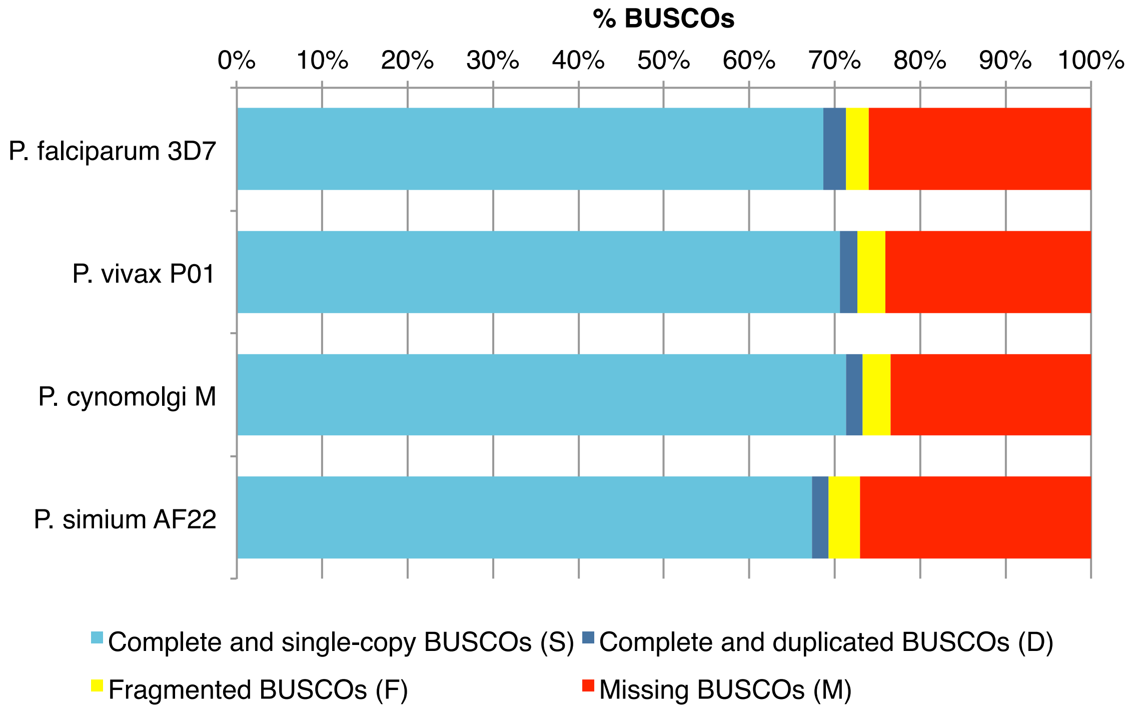
% BUSCOs
0%
10%
20%
30%
40%
50%
60%
70%
80%
90%
100%
P. falciparum 3D7
P. vivax P01
P. cynomolgi M
P. simium AF22
Complete and single-copy BUSCOs (S)
Complete and duplicated BUSCOs (D)
Fragmented BUSCOs (F)
Missing BUSCOs (M)

Figure S3

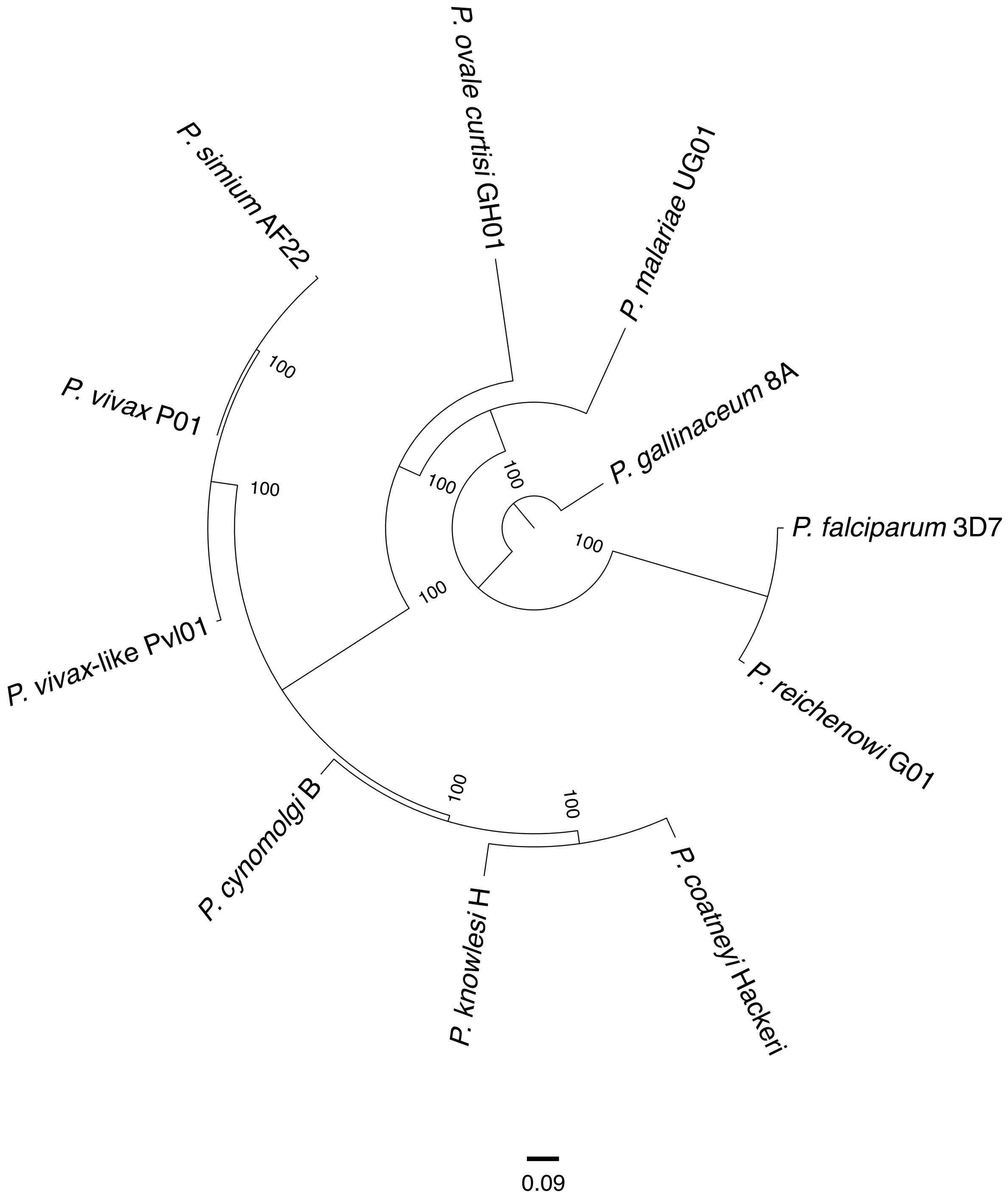
P. ovale curtisi GH01
P. malariae UG01
P. simium AF22
P. vivax P01
P. gallinaceum 8A
P. falciparum 3D7
P. vivax-like Pvl01
P. reichenowi G01
P. cynomolgi B
P. knowlesi H
P. coatneyi Hackeri
100
100
100
100
100
100
100
100
0.09

Figure S4

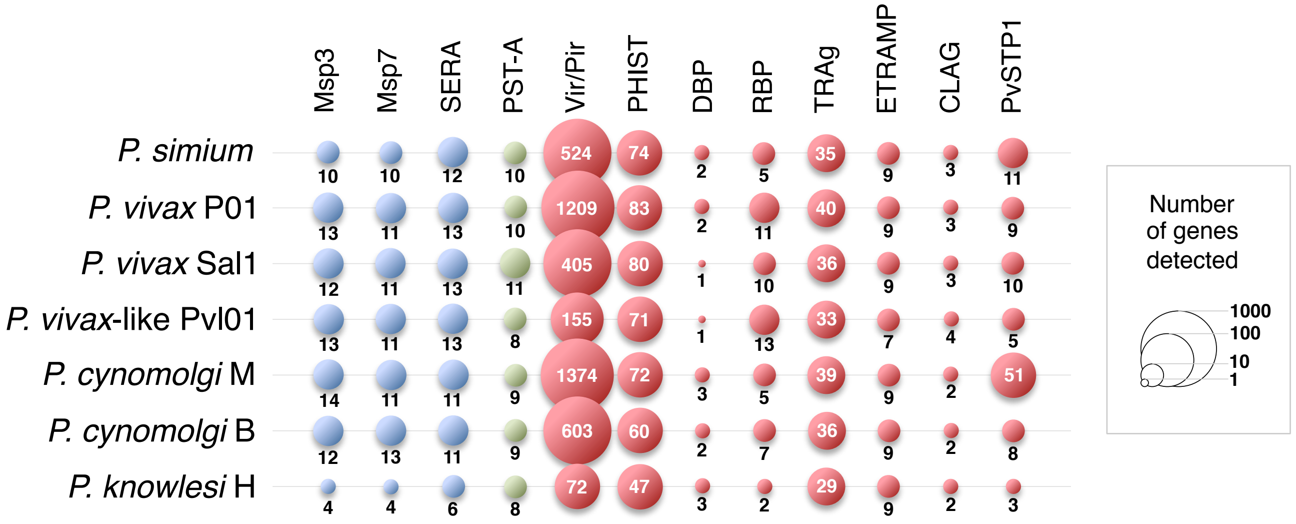

Msp3
Msp7
SERA
PST-A
Vir/Pir
PHIST
DBP
RBP
TRAg
ETRAMP
CLAG
PvSTP1
P. simium 10 10 12 10 524 74 2 5 35 9 3 11
P. vivax P01 13 11 13 10 1209 83 2 11 40 9 3 9
P. vivax Sal1 12 11 13 11 405 80 1 10 36 9 3 10
P. vivax-like PvI01 13 11 13 8 155 71 1 13 33 7 4 5
P. cynomolgi M 14 11 11 9 1374 72 3 5 39 9 2 51
P. cynomolgi B 12 13 11 9 603 60 2 7 36 9 2 8
P. knowlesi H 4 4 6 8 72 47 3 2 29 9 2 3
Number of genes detected
1000
100
10
1

Figure S5

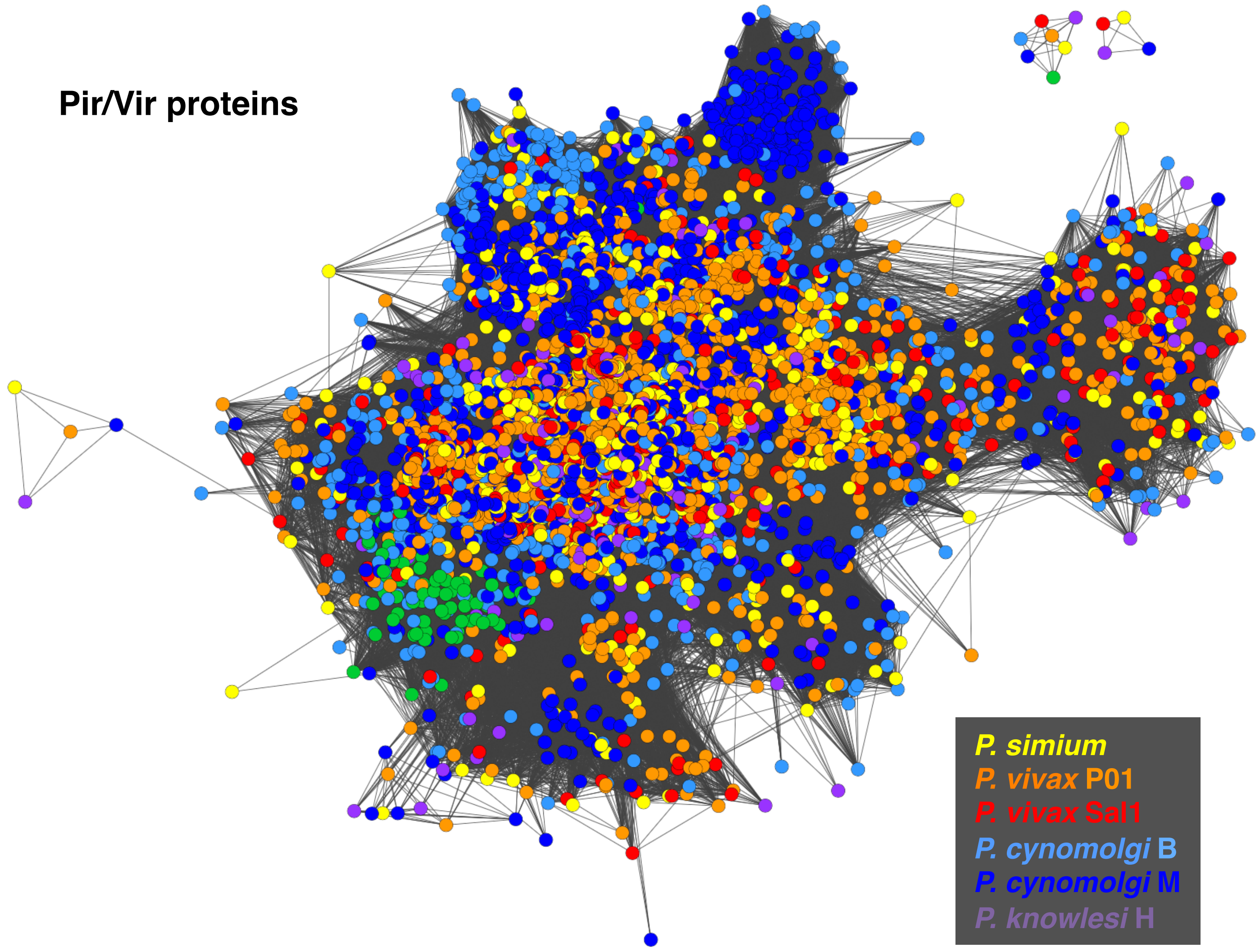
Pir/Vir proteins
P. simium
P. vivax P01
P. vivax Sal1
P. cynomolgi B
P. cynomolgi M
P. knowlesi H

Figure S6

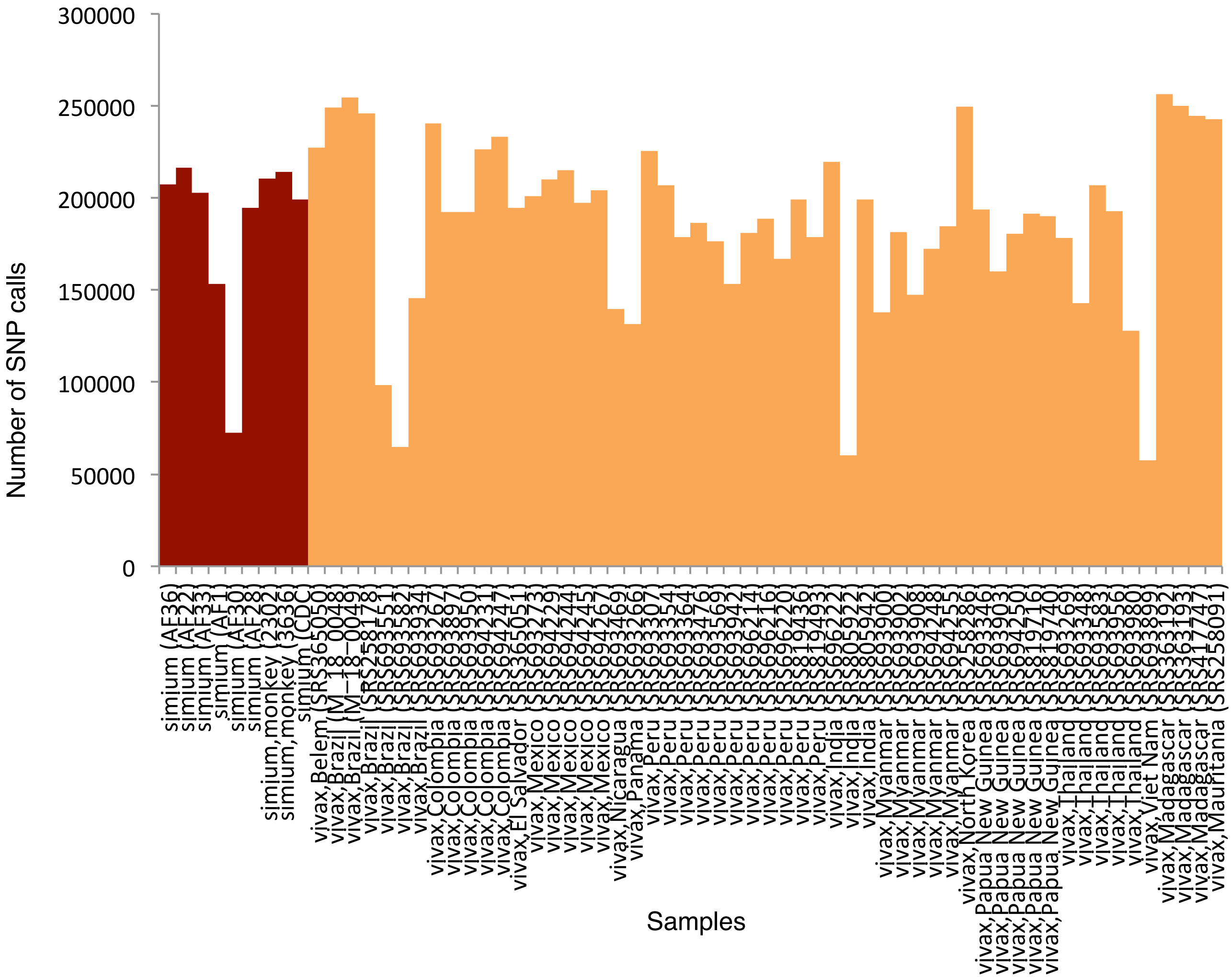

Number of SNP calls
300000
250000
200000
150000
100000
50000
0
Samples


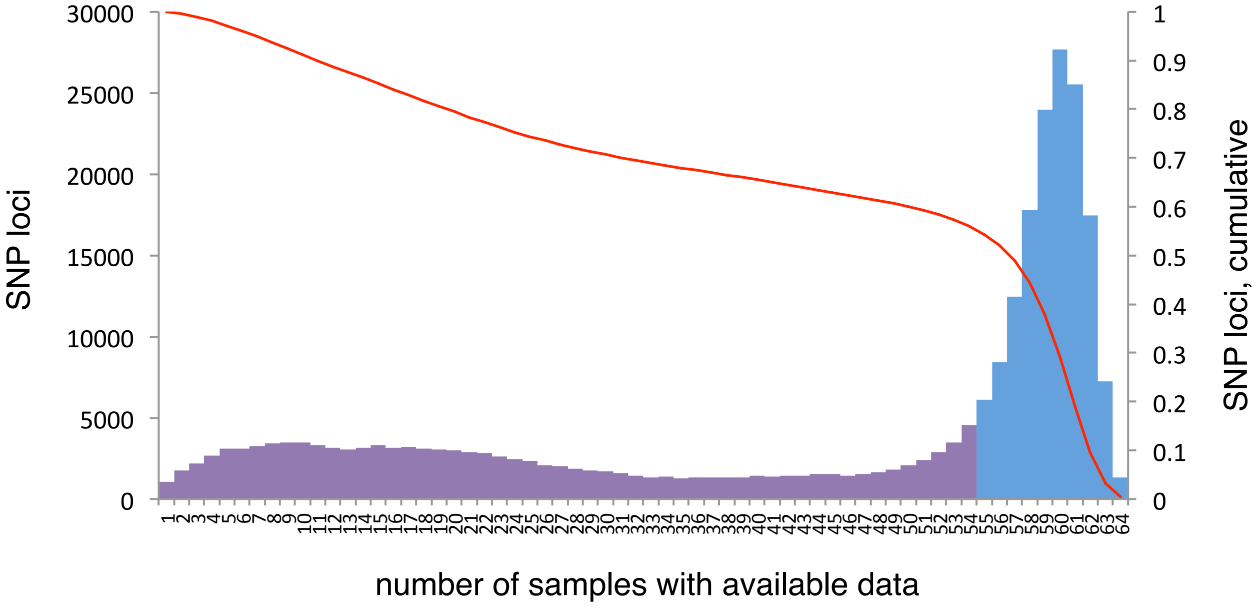

SNP loci
30000
25000
20000
15000
10000
5000
0
SNP loci, cumulative
1
0.9
0.8
0.7
0.6
0.5
0.4
0.3
0.2
0.1
0
number of samples with available data

Figure S7

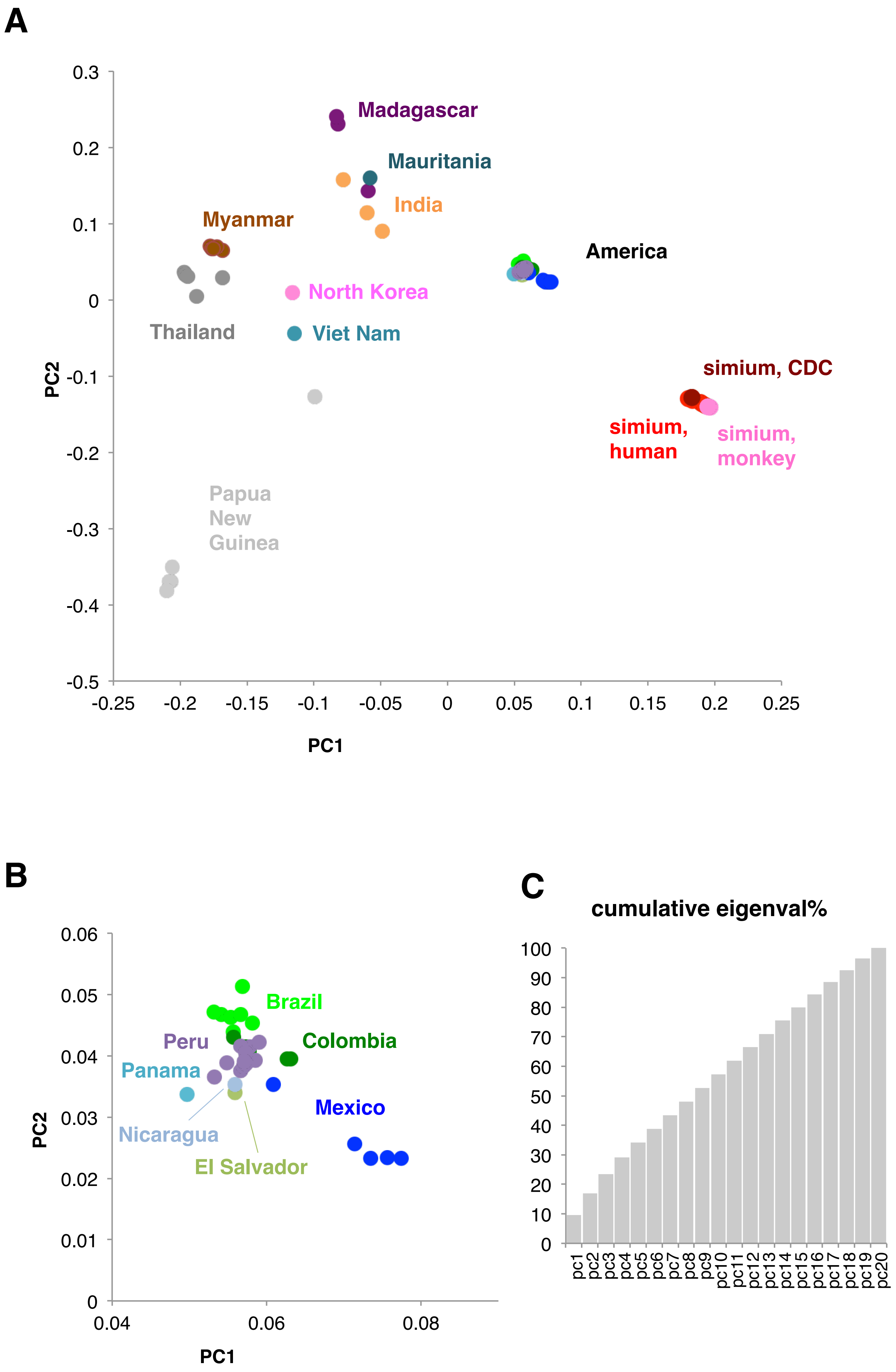
A
Madagascar
Mauritania
India
Myanmar
America
North Korea
Thailand
Viet Nam
simium, CDC
simium, human
simium, monkey
Papua New Guinea
PC2
PC1
0.3
0.2
0.1
0
-0.1
-0.2
-0.3
-0.4
-0.5
-0.25
-0.2
-0.15
-0.1
-0.05
0
0.05
0.1
0.15
0.2
0.25
B
Brazil
Peru
Colombia
Panama
Mexico
Nicaragua
El Salvador
PC2
PC1
0.06
0.05
0.04
0.03
0.02
0.01
0
0.04
0.06
0.08
C
cumulative eigenval%
100
90
80
70
60
50
40
30
20
10
0
pc1
pc2
pc3
pc4
pc5
pc6
pc7
pc8
pc9
pc10
pc11
pc12
pc13
pc14
pc15
pc16
pc17
pc18
pc19
pc20

Figure S8

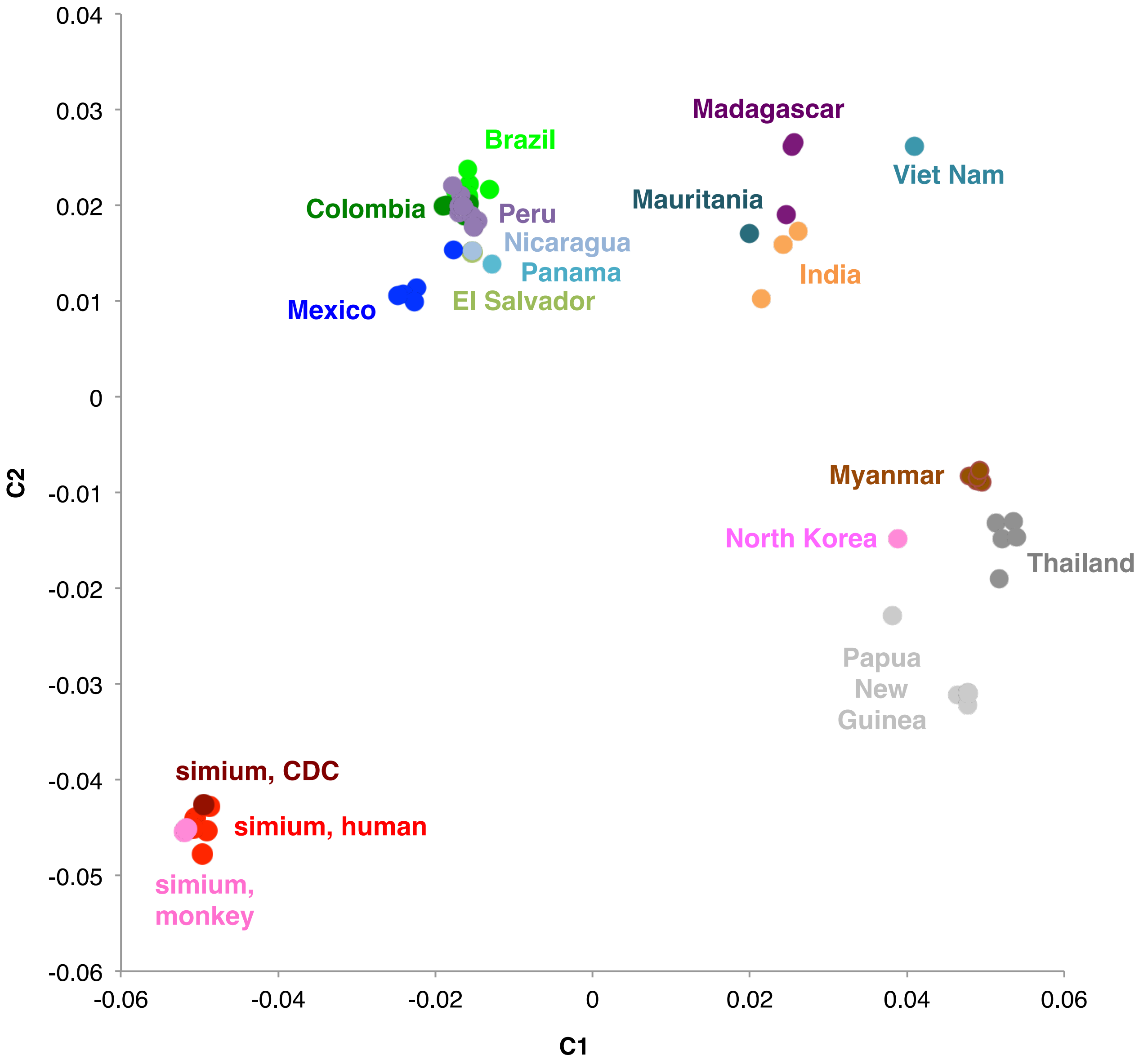

0.04
0.03
0.02
0.01
0
-0.01
-0.02
-0.03
-0.04
-0.05
-0.06
C2
-0.06
-0.04
-0.02
0
0.02
0.04
0.06
C1
Brazil
Colombia
Peru
Nicaragua
Panama
Mexico
El Salvador
Madagascar
Viet Nam
Mauritania
India
Myanmar
North Korea
Thailand
Papua
New
Guinea
simium, CDC
simium, human
simium,
monkey

Figure S9

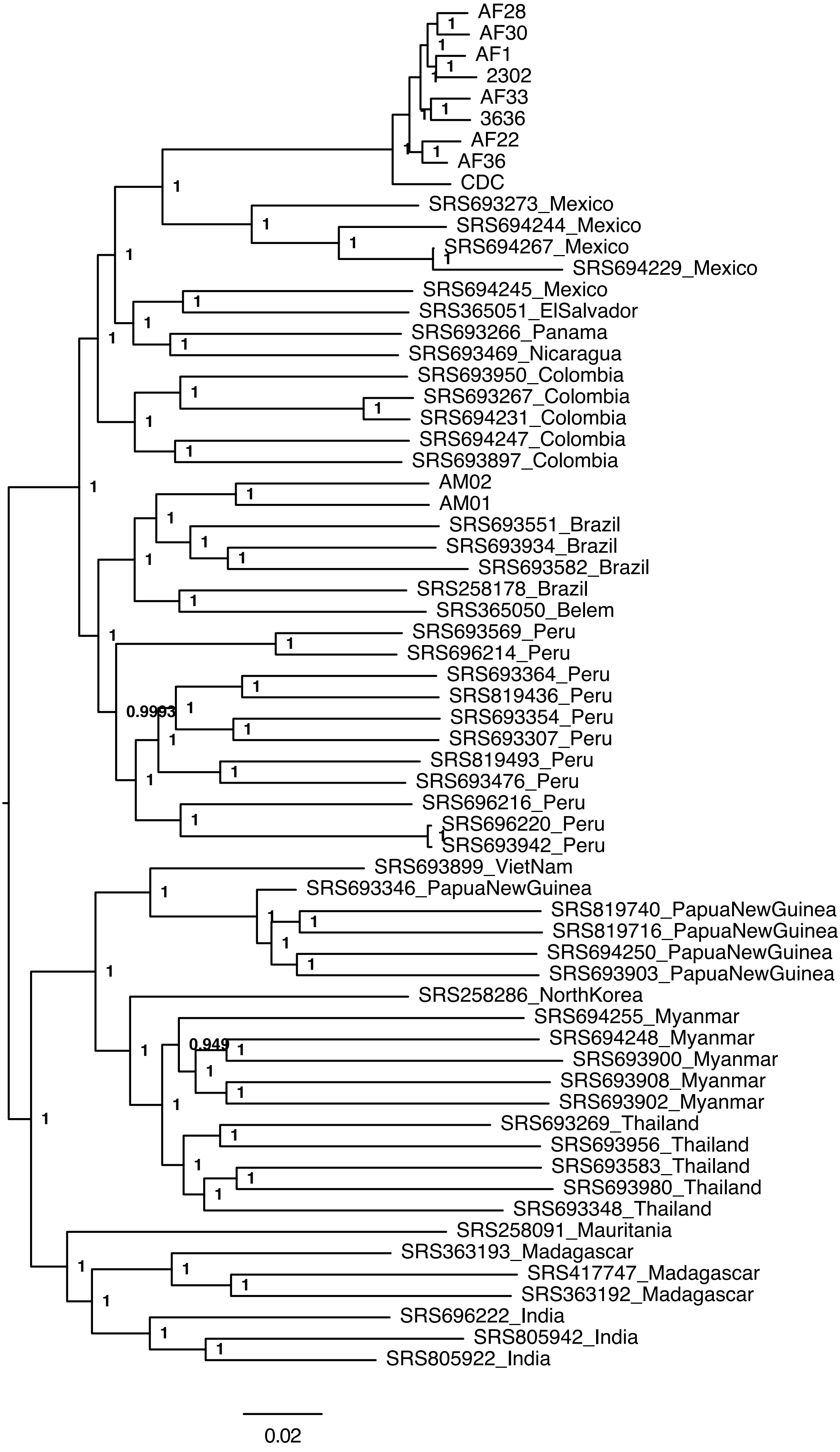
AF28
AF30
AF1
2302
AF33
3636
AF22
AF36
CDC
SRS693273_Mexico
SRS694244_Mexico
SRS694267_Mexico
SRS694229_Mexico
SRS694245_Mexico
SRS365051_ElSalvador
SRS693266_Panama
SRS693469_Nicaragua
SRS693950_Colombia
SRS693267_Colombia
SRS694231_Colombia
SRS694247_Colombia
SRS693897_Colombia
AM02
AM01
SRS693551_Brazil
SRS693934_Brazil
SRS693582_Brazil
SRS258178_Brazil
SRS365050_Belem
SRS693569_Peru
SRS696214_Peru
SRS693364_Peru
SRS819436_Peru
SRS693354_Peru
SRS693307_Peru
SRS819493_Peru
SRS693476_Peru
SRS696216_Peru
SRS696220_Peru
SRS693942_Peru
SRS693899_VietNam
SRS693346_PapuaNewGuinea
SRS819740_PapuaNewGuinea
SRS819716_PapuaNewGuinea
SRS694250_PapuaNewGuinea
SRS693903_PapuaNewGuinea
SRS258286_NorthKorea
SRS694255_Myanmar
SRS694248_Myanmar
SRS693900_Myanmar
SRS693908_Myanmar
SRS693902_Myanmar
SRS693269_Thailand
SRS693956_Thailand
SRS693583_Thailand
SRS693980_Thailand
SRS693348_Thailand
SRS258091_Mauritania
SRS363193_Madagascar
SRS417747_Madagascar
SRS363192_Madagascar
SRS696222_India
SRS805942_India
SRS805922_India
0.9993
0.949
0.02

Figure S10

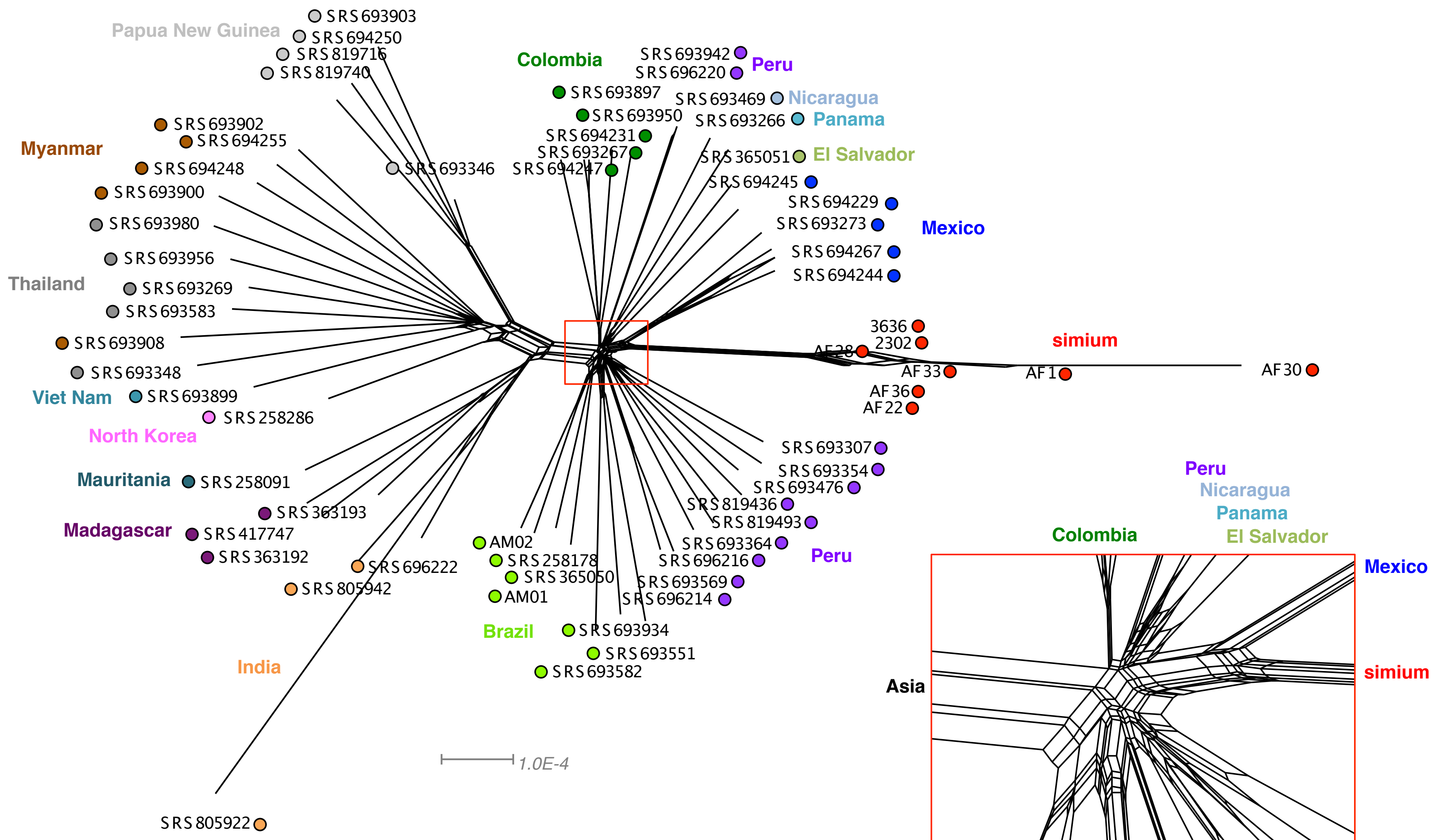

Papua New Guinea
SRS693903
SRS694250
SRS819716
SRS819740
SRS693346
Myanmar
SRS693902
SRS694255
SRS694248
SRS693900
SRS693980
SRS693956
Thailand
SRS693269
SRS693583
SRS693908
SRS693348
Viet Nam
SRS693899
SRS258286
North Korea
Mauritania
SRS258091
SRS363193
Madagascar
SRS417747
SRS363192
SRS696222
SRS805942
India
SRS805922
Colombia
SRS693897
SRS693950
SRS694231
SRS693267
SRS694247
SRS693942
SRS696220
Peru
SRS693469
Nicaragua
SRS693266
Panama
SRS365051
El Salvador
SRS694245
SRS694229
SRS693273
Mexico
SRS694267
SRS694244
3636
2302
AF28
simium
AF33
AF1
AF30
AF36
AF22
SRS693307
SRS693354
SRS693476
SRS819436
SRS819493
SRS693364
SRS696216
Peru
SRS693569
SRS696214
AM02
SRS258178
SRS365050
AM01
Brazil
SRS693934
SRS693551
SRS693582
1.0E-4
Peru
Nicaragua
Panama
Colombia
El Salvador
Mexico
Asia
simium
Brazil
Peru

Figure S11

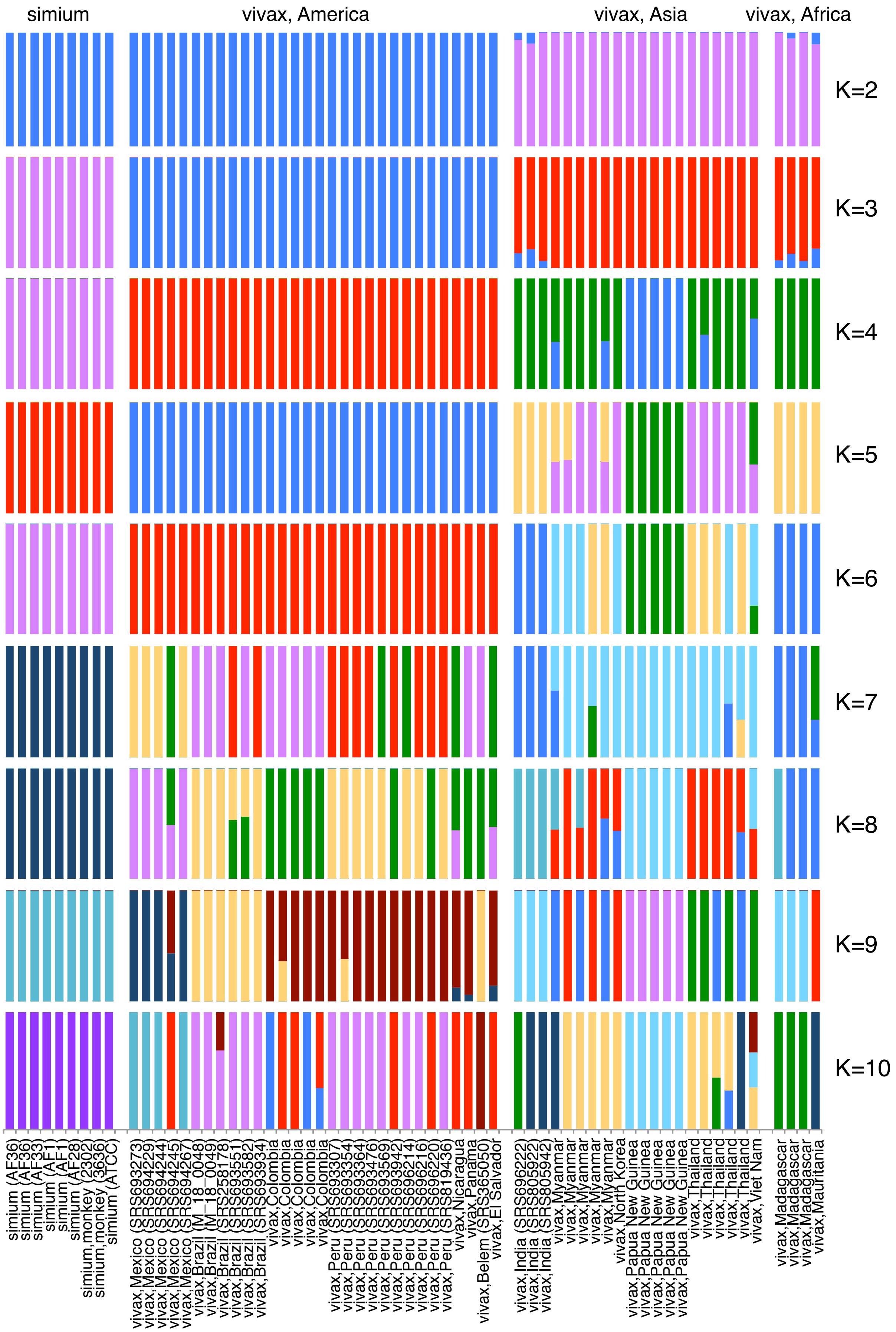

simium
vivax, America
vivax, Asia
vivax, Africa
K=2
K=3
K=4
K=5
K=6
K=7
K=8
K=9
K=10
simium (AF36)
simium (AF36)
simium (AF33)
simium (AF1)
simium (AF1)
simium (AF28)
simium,monkey (2302)
simium,monkey (3636)
simium (ATCC)
vivax,Mexico (SRS693273)
vivax,Mexico (SRS694229)
vivax,Mexico (SRS694244)
vivax,Mexico (SRS694245)
vivax,Mexico (SRS694267)
vivax,Brazil (M_18_0048)
vivax,Brazil (M_18_0049)
vivax,Brazil (SRS258178)
vivax,Brazil (SRS693551)
vivax,Brazil (SRS693582)
vivax,Brazil (SRS693934)
vivax,Colombia
vivax,Colombia
vivax,Colombia
vivax,Colombia
vivax,Peru (SRS693307)
vivax,Peru (SRS693354)
vivax,Peru (SRS693364)
vivax,Peru (SRS693476)
vivax,Peru (SRS693569)
vivax,Peru (SRS693942)
vivax,Peru (SRS696214)
vivax,Peru (SRS696216)
vivax,Peru (SRS696220)
vivax,Peru (SRS819436)
vivax,Nicaragua
vivax,Panama
vivax,Belem (SRS365050)
vivax,El Salvador
vivax,India (SRS696222)
vivax,India (SRS805922)
vivax,India (SRS805942)
vivax,Myanmar
vivax,Myanmar
vivax,Myanmar
vivax,Myanmar
vivax,Myanmar
vivax,North Korea
vivax,Papua New Guinea
vivax,Papua New Guinea
vivax,Papua New Guinea
vivax,Papua New Guinea
vivax,Papua New Guinea
vivax,Thailand
vivax,Thailand
vivax,Thailand
vivax,Thailand
vivax,Thailand
vivax,Viet Nam
vivax,Madagascar
vivax,Madagascar
vivax,Madagascar
vivax,Mauritania

Figure S12

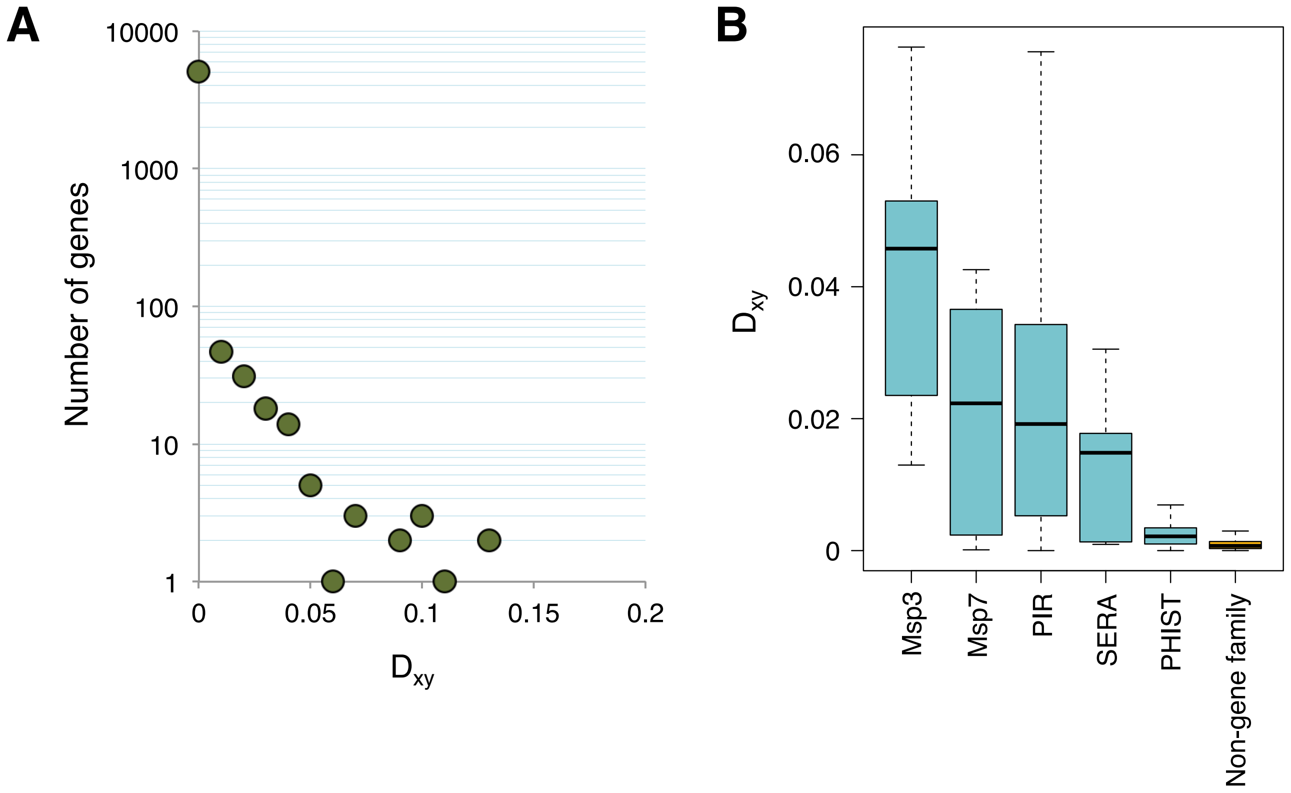
A
10000
1000
100
10
1
Number of genes
0
0.05
0.1
0.15
0.2
Dxy
B
0.06
0.04
0.02
0
Dxy
Msp3
Msp7
PIR
SERA
PHIST
Non-gene family

Figure S13

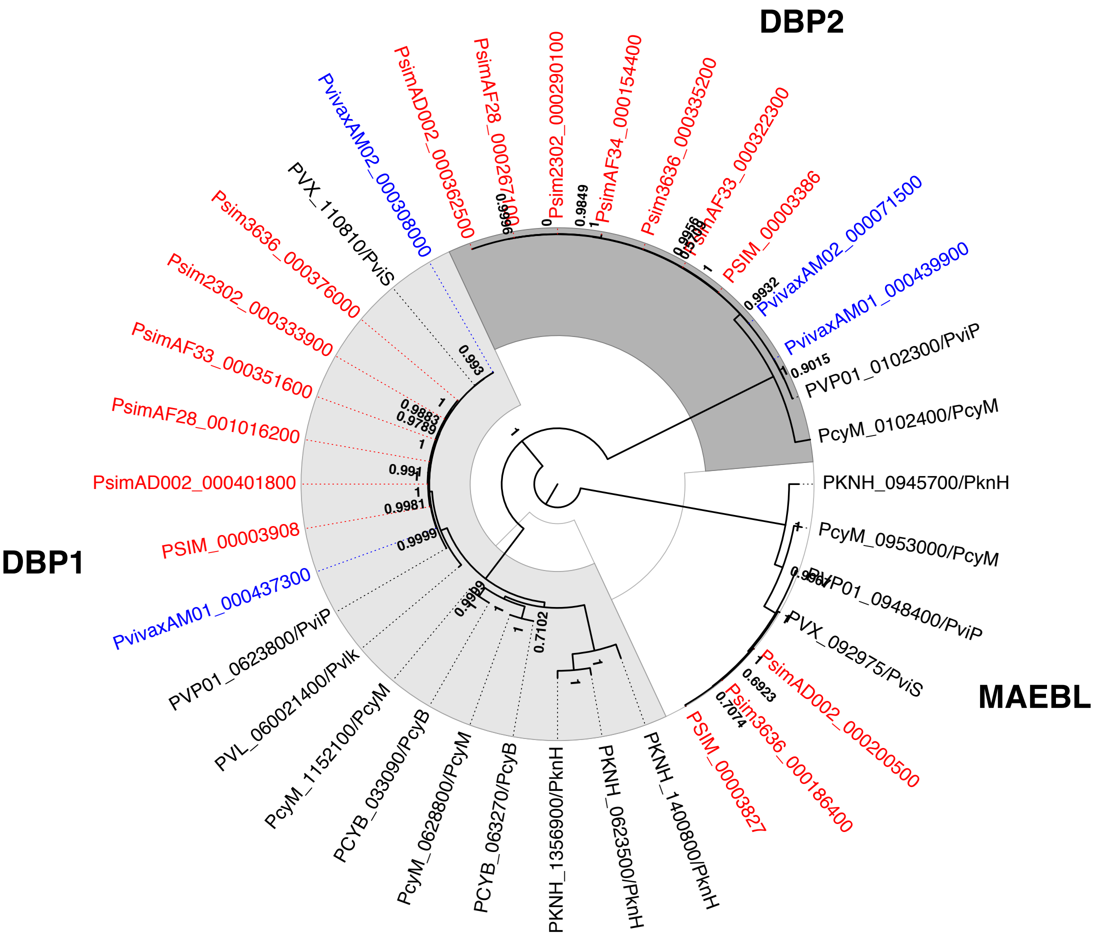

DBP2
DBP1
MAEBL
PvivaxAM02_000308000
PsimAD002_000362500
PsimAF28_000267100
Psim2302_000290100
PsimAF34_000154400
Psim3636_000335200
PsimAF33_000322300
PSIM_00003386
PvivaxAM02_000071500
PvivaxAM01_000439900
PVP01_0102300/PviP
PcyM_0102400/PcyM
PKNH_0945700/PknH
PcyM_0953000/PcyM
PVP01_0948400/PviP
PVX_092975/PviS
PsimAD002_000200500
Psim3636_000186400
PSIM_00003827
PKNH_1400800/PknH
PKNH_0623500/PknH
PKNH_1356900/PknH
PCYB_063270/PcyB
PcyM_0628800/PcyM
PCYB_033090/PcyB
PcyM_1152100/PcyM
PVL_060021400/PvIk
PVP01_0623800/PviP
PvivaxAM01_000437300
PSIM_00003908
PsimAD002_000401800
PsimAF28_001016200
PsimAF33_000351600
Psim2302_000333900
Psim3636_000376000
PVX_110810/PviS

Figure S14

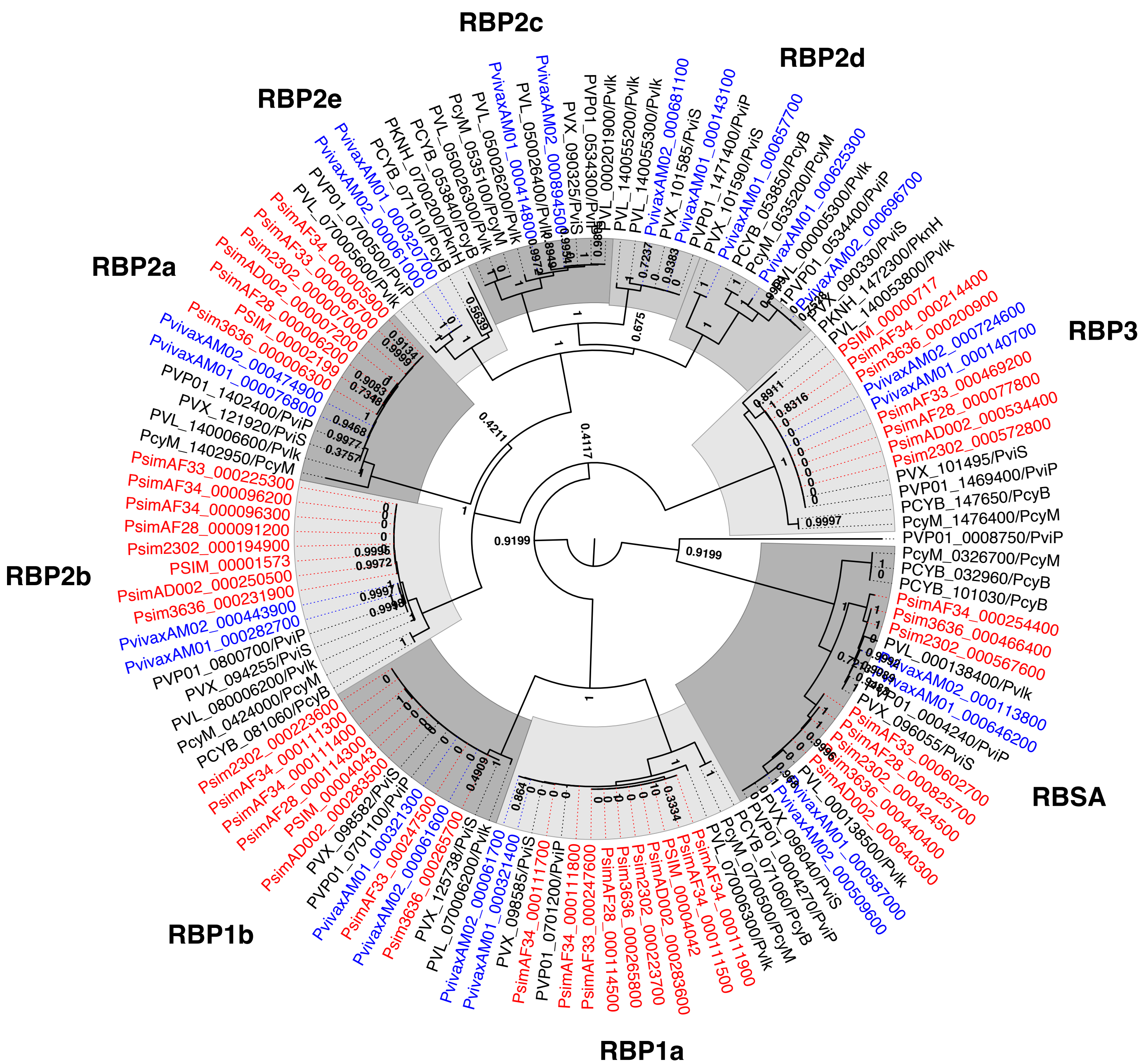
RBP2c
RBP2d
RBP2e
RBP2a
RBP2b
RBP3
RBSA
RBP1a
RBP1b
PSIM_00000717
PsimAF34_000214400
Psim3636_000200900
PvivaxAM02_000724600
PvivaxAM01_000140700
PsimAF33_000469200
PsimAF28_000077800
PsimAD002_000534400
Psim2302_000572800
PVX_101495/PviS
PVP01_1469400/PviP
PCYB_147650/PcyB
PcyM_1476400/PcyM
PVP01_0008750/PviP
PcyM_0326700/PcyM
PCYB_032960/PcyB
PCYB_101030/PcyB
PsimAF34_000254400
Psim3636_000466400
Psim2302_000567600
PVL_000138400/PvIk
PvivaxAM02_000113800
PvivaxAM01_000646200
PVP01_0004240/PviP
PVX_096055/PviS
PsimAF33_000602700
PsimAF28_000825700
Psim3636_000424500
PsimAD002_000440400
PVL_000138500/PvIk
PvivaxAM02_000587000
PvivaxAM01_000509600
PVX_096040/PviS
PVP01_0004270/PviP
PCYB_071060/PcyB
PcyM_0700500/PcyM
PVL_070006300/PvIk
PsimAF34_000111500
PSIM_00004042
PsimAD002_000283600
Psim2302_000223700
Psim3636_000265800
PsimAF28_000114500
PsimAF33_000247600
PsimAF34_000111800
PVP01_0701200/PviP
PsimAF34_000111700
PVX_098585/PviS
PvivaxAM01_000321400
PvivaxAM02_000061700
PVL_070006200/PvIk
PVX_125738/PviS
Psim3636_000265700
PvivaxAM02_000061600
PsimAF33_000247500
PvivaxAM01_000321300
PVP01_0701100/PviP
PVX_098582/PviS
PsimAD002_000283500
PSIM_00004043
PsimAF28_000114400
PsimAF34_000111300
Psim2302_000223600
PCYB_081060/PcyB
PcyM_0424000/PcyM
PVL_080006200/PvIk
PVX_094255/PviS
PVP01_0800700/PviP
PvivaxAM01_000282700
PvivaxAM02_000443900
Psim3636_000231900
PsimAD002_000250500
PSIM_00001573
Psim2302_000194900
PsimAF28_000091200
PsimAF34_000096300
PsimAF34_000096200
PsimAF33_000225300
PcyM_1402950/PcyM
PVL_140006600/PvIk
PVX_121920/PviS
PVP01_1402400/PviP
PvivaxAM01_000076800
PvivaxAM02_000474900
Psim3636_000006300
PSIM_00002199
PsimAF28_000006200
PsimAD002_000007200
Psim2302_000007000
PsimAF33_000006700
PsimAF34_000005900
PVL_070005600/PvIk
PVP01_0700500/PviP
PvivaxAM02_000061000
PvivaxAM01_000320700
PCYB_071010/PcyB
PKNH_0700200/PknH
PCYB_053840/PcyB
PcyM_0535100/PcyM
PVL_050026300/PvIk
PvivaxAM01_000414800
PVL_050026400/PvIk
PvivaxAM02_000894500
PVX_090325/PviS
PVP01_0534300/PviP
PVL_000201900/PvIk
PVL_140055200/PvIk
PVL_140055300/PvIk
PvivaxAM02_000681100
PVX_101585/PviS
PvivaxAM01_000143100
PVP01_1471400/PviP
PVX_101590/PviS
PvivaxAM01_000657700
PCYB_053850/PcyB
PcyM_0535200/PcyM
PvivaxAM01_000625300
PVL_000005300/PvIk
PVP01_0534400/PviP
PvivaxAM02_000696700
PVX_090330/PviS
PKNH_1472300/PknH
PVL_140053800/PvIk

Figure S15

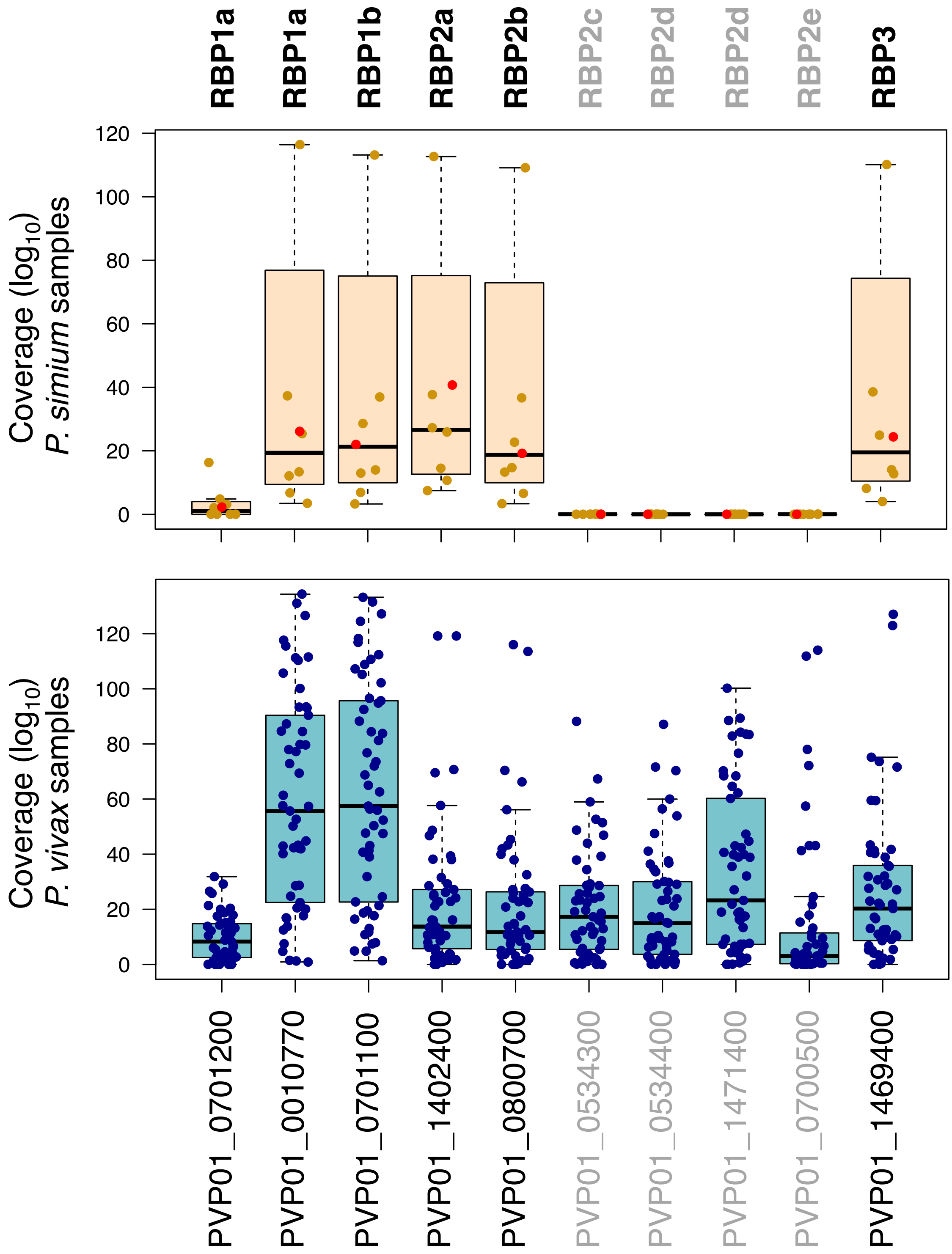

RBP1a
RBP1a
RBP1b
RBP2a
RBP2b
RBP2c
RBP2d
RBP2d
RBP2e
RBP3
Coverage (log10)
P. simium samples
0
20
40
60
80
100
120
Coverage (log10)
P. vivax samples
0
20
40
60
80
100
120
PVP01_0701200
PVP01_0010770
PVP01_0701100
PVP01_1402400
PVP01_0800700
PVP01_0534300
PVP01_0534400
PVP01_1471400
PVP01_0700500
PVP01_1469400

### Figure S16

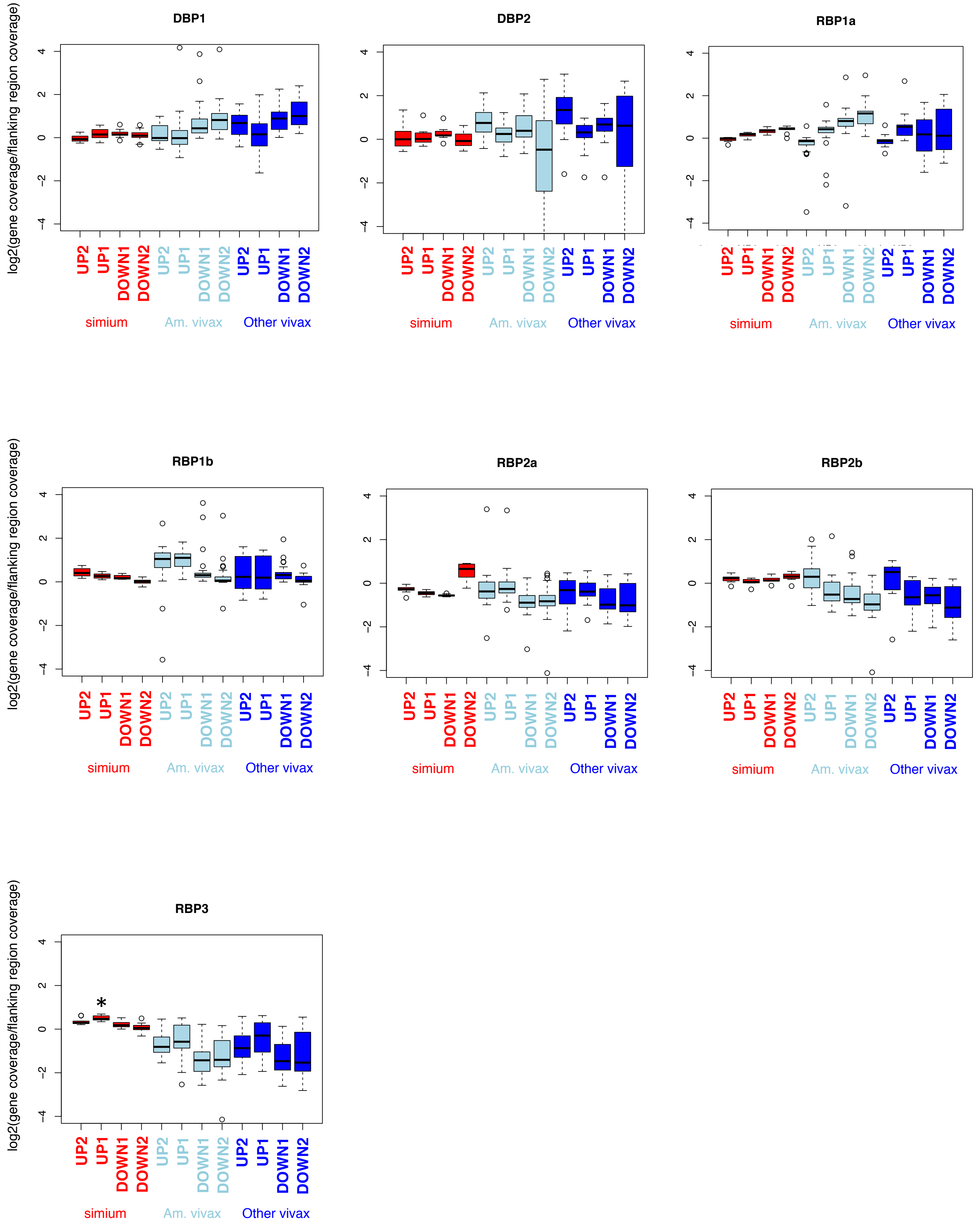
DBP1
DBP2
RBP1a
RBP1b
RBP2a
RBP2b
RBP3
log2(gene coverage/flanking region coverage)
UP2
UP1
DOWN1
DOWN2
simium
Am. vivax
Other vivax
*

### Figure S17

**Complete alignment of DBP1 protein sequences**

```
PCYB_063270-t26_1-p1       MKGKNRPLFFLLVLLLSHKVNNVLLERTIETLQECKNECVKGENCYKLSKGHHYIEEDNIERWLQGTNERRSEENIKYKY
PcyM_0628800-t36_1-p1      MKGKNRPLFFLLVLLLSHKVNNVLLERTIETLQECKNECVKGENCYKLSKGHHYIEEDNIERWLQGTNERRSEENIKYKY
PVP01_0623800.1-p1         MKGKNRSLFVLLVLLLSHKVNNVLLERTIETLLECKNEYVKGENGYKLAKGHHCVEEDNLERWLQGTNERRSEENIKYKY
PVX_110810.1-p1            MKGKNRSLFVLLVLLLLHKVNNVLLERTIETLLECKNEYVKGENGYKLAKGHHCVEEDNLERWLQGTNERRSEENIKYKY
Psim2302_000333900.1       MKGKNRSLFVLLVLLLSHKVNNVLLERTIETLLECKNEYVKGENGYKLAKGHHCVEEDNLERWLQGTNERRSEENIKYKY
Psim3636_000376000.1       MKGKNRSLFVLLVLLLSHKVNNVLLERTIETLLECKNEYVKGENGYKLAKGHHCVEEDNLERWLQGTNERRSEENIKYKY
PsimAD002_000401800.1      MKGKNRSLFVLLVLLLSHKVNNVLLERTIETLLECKNEYVKGENGYKLAKGHHCVEEDNLERWLQGTNERRSEENIKYKY
PsimAD005_000131600.1      MKGKNRSLFVLLVLLLSHKVNNVLLERTIETLLECKNEYVKGENGYKLAKGHHCVEEDNLERWLQGTNERRSEENIKYKY
PsimAF28_001016200.1       MKGKNRSLFVLLVLLLSHKVNNVLLERTIETLLECKNEYVKGENGYKLAKGHHCVEEDNLERWLQGTNERRSEENIKYKY
PsimAF33_000351600.1       MKGKNRSLFVLLVLLLSHKVNNVLLERTIETLLECKNEYVKGENGYKLAKGHHCVEEDNLERWLQGTNERRSEENIKYKY
PvivaxAM01_000437300.1     MKGKNRSLFVLLVLLLLHKVNNVLLERTIETLLECKNEYVKGENGYKLAKGHHCVEEDNLERWLQGTNERRSEENIKYKY
PvivaxAM02_000308000.1     MKGKNRSLFVLLVLLLLHKVNNVLLERTIETLLECKNEYVKGENGYKLAKGHHCVEEDNLERWLQGTNERRSEENIKYKY
AAZ81533.1                 MKGKNRSLFVLLVLLLLHKVNNVLLERTIETLLECKNEYVKGENGYKLAKGHHCVEEDNLERWLQGTNERRSEENIKYKY
AAZ81530.1                 MKGKNRSLFVLLVLLLLHKVNNVLLERTIETLLECKNEYVKGENGYKLAKGHHCVEEDNLERWLQGTNERRSEENIKYKY
ABA39301.1                 MKGKNRSLFVLLVLLLLHKVNNVLLERTIETLLECKNEYVKGENGYKLAKGHHCVEEDNLERWLQGTNERRSEENIKYKY
ACB42428.1                 MKGKNRSLFVLLVLLLLHKVNNVLLERTIETLLECKNEYVKGENGYKLAKGHHCVEEDNLERWLQGTNERRSEENIKYKY
ACB42432                   MKGKNRSLFVLLVLLLLHKVNNVLLERTIETLLECKNEYVKGENGYKLAKGHHCVEEDNLERWLQGTNERRSEENIKYKY
PVL_060021400-t42_1-p1     MKGKNRSLFVLLVLLLSHKVNNVLLERTIETLLECKNEYVKGENGYKLAKGHHCVEEDNLERWLQGTNERRSEENVKYKY
ACB42432.1                 MKGKNRSLFVLLVLLLLHKVNNVLLERTIETLLECKNEYVKGENGYKLAKGHHCVEEDNLERWLQGTNERRSEENIKYKY


PCYB_063270-t26_1-p1       GVTELKIKYEQMNGKRTSRILKESIYEAQNFRDNNYR----------EEKDGEHKTDSKTDNGRSANNLVMLDYDTSSSG
PcyM_0628800-t36_1-p1      GVTELKIKYEQMNGKRTSRILKESIYEAQNFRDNNYR----------EEKDGEHKTDSKTDNGRSANNLVMLDYDTSSSG
PVP01_0623800.1-p1         GVTELKIKYAQMNGKRSSRILKESIYGAHNFGGNSYMEGKDGGDKTGEEKDGEHKTDSKTDNGKGANNLVMLDYETSSNG
PVX_110810.1-p1            GVTELKIKYAQMNGKRSSRILKESIYGAHNFGGNSYMEGKDGGDKTGEEKDGEHKTDSKTDNGKGANNLVMLDYETSSNG
Psim2302_000333900.1       GVTELKIKYAQMNGKRSSRILKESIYGAHNFGGNSYMEGKDGGDKTGEEKDGEHKTDSKTDNGKGANNLVMLDYETSSNG
Psim3636_000376000.1       GVTELKIKYAQMNGKRSSRILKESIYGAHNFGGNSYMEGKDGGDKTGEEKDGEHKTDSKTDNGKGANNLVMLDYETSSNG
PsimAD002_000401800.1      GVTELKIKYAQMNGKRSSRILKESIYGAHNFGGNSYMEGKDGGDKTGEEKDGEHKTDSKTDNGKGANNLVMLDYETSSNG
PsimAD005_000131600.1      GVTELKIKYAQMNGKRSSRILKESIYGAHNFGGNSYMEGKDGGDKTGEEKDGEHKTDSKTDNGKGANNLVMLDYETSSNG
PsimAF28_001016200.1       GVTELKIKYAQMNGKRSSRILKESIYGAHNFGGNSYMEGKDGGDKTGEEKDGEHKTDSKTDNGKGANNLVMLDYETSSNG
PsimAF33_000351600.1       GVTELKIKYAQMNGKRSSRILKESIYGAHNFGGNSYMEGKDGGDKTGEEKDGEHKTDSKTDNGKGANNLVMLDYETSSNG
PvivaxAM01_000437300.1     GVTELKIKYAQMNGKRSSRILKESIYGAHNFGGNSYMEGKDGGDKTGEEKDGEHKTDSKTDNGKGANNLVMLDYETSSNG
PvivaxAM02_000308000.1     GVTELKIKYAQMNGKRSSRILKESIYGAHNFGGNSYMEGKDGGDKTGEEKDGEHKTDSKTDNGKGANNLVMLDYETSSNG
AAZ81533.1                 GVTELKIKYAQMNGKRSSRILKESIYGAHNFGGNSYMEGKDGGDKTGEEKDGEHKTDSKTDNGKGANNLVMLDYETSSNG
AAZ81530.1                 GVTELKIKYAQMNGKRSSRILKESIYGAHNFGGNSYMEGKDGGDKTGEEKDGEHKTDSKTDNGKGENNLVMLDYETSSNG
ABA39301.1                 GVTELKIKYAQMNGKRSSRILKESIYGAHNFGGNSYMEGKDGGDKTGEEKDGEHKTDSKTDNGKGANNLVMLDYETSSNG
ACB42428.1                 GVTELKIKYAQMNGKRSSRILKESIYGAHNFGGNSYMEGKDGGDKTGEEKDGEHKTDSKTDNGKGANNLVMLDYETSSNG
ACB42432                   GVTELKIKYAQMNGKRSSRILKESIYGAHNFGGNSYMEGKDGGDKTGEEKDGEHKTDSKTDNGKGANNLVMLDYETSSNG
PVL_060021400-t42_1-p1     DVTELKIKYAQINGKRSSRILKESIYGAHNFGGNSYMEGKDGGDKIGEEKNGEHKTDSETDNGKGANNLVMLDYGTSSNG
ACB42432.1                 GVTELKIKYAQMNGKRSSRILKESIYGAHNFGGNSYMEGKDGGDKTGEEKDGEHKTDSKTDNGKGANNLVMLDYETSSNG


PCYB_063270-t26_1-p1       HPAVTLDNVLEFVTDHGENSLENSSNGGNLYDIVHKKTMSSGVINHAFLQNSEIVTCNDKRKRQVRDWDCSTKKDVCIPD
PcyM_0628800-t36_1-p1      HPAVTLDNVLEFVTDHGENSLENSSNGGNLYDIVHKKTMSSGVINHAFLQNSEIVTCNDKRKRQVRDWDCSTKKDVCIPD
PVP01_0623800.1-p1         QPAGTLDNVLEFVTGHEGNSRKNSSNGGNPYDIDHKKTISSAIINHAFLQNTVMKNCNYKRKRRERDWDCNTKKDVCIPD
PVX_110810.1-p1            QPAGTLDNVLEFVTGHEGNSRKNSSNGGNPYDIDHKKTISSAIINHAFLQNTVMKNCNYKRKRRERDWDCNTKKDVCIPD
Psim2302_000333900.1       QPAGTLDNVLEFVTGHEGNSRKNSSNGGNPYDIDHKKTISSAIINHAFLQNTVMNNCNYKRKRRERDWDCNTKKDVCIPD
Psim3636_000376000.1       QPAGTLDNVLEFVTGHEGNSRKNSSNGGNPYDIDHKKTISSAIINHAFLQNTVMNNCNYKRKRRERDWDCNTKKDVCIPD
PsimAD002_000401800.1      QPAGTLDNVLEFVTGHEGNSRKNSSNGGNPYDIDHKKTISSAIINHAFLQNTVMNNCNYKRKRRERDWDCNTKKDVCIPD
PsimAD005_000131600.1      QPAGTLDNVLEFVTGHEGNSRKNSSNGGNPYDIDHKKTISSAIINHAFLQNTVMKNCNYKRKRRERDWDCNTKKDVCIPD
PsimAF28_001016200.1       QPAGTLDNVLEFVTGHEGNSRKNSSNGGNPYDIDHKKTISSAIINHAFLQNTVMNNCNYKRKRRERDWDCNTKKDVCIPD
PsimAF33_000351600.1       QPAGTLDNVLEFVTGHEGNSRKNSSNGGNPYDIDHKKTISSAIINHAFLQNTVMNNCNYKRKRRERDWDCNTKKDVCIPD
PvivaxAM01_000437300.1     QPAGTLDNVLEFVTGHEGNSRKNSSNGGNPYDIDHKKTISSAIINHAFLQNTVMKNCNYKRKRRERDWDCNTKKDVCIPD
PvivaxAM02_000308000.1     EPAGTLDNVLEFVTGHEGNSRKNSSNGGNPYDIDHKKTISSAIINHAFLQNTVMKNCNYKRKRRERDWDCNTKKDVCIPD
AAZ81533.1                 QPAGTLDNVLEFVTGHEGNSRKNSSNGGNPYDIDHKKTISSAIINHAFLQNTVMKNCNYKRKRRERDWDCNTKKDVCIPD
AAZ81530.1                 QPAGTLDNVLEFVTEHEGNSRKNSSNGGNPYDIDHKKTISSAIINHAFLQNTVMKNCNYKRKRRERDWDCNTKKDVCIPD
ABA39301.1                 QPAGTLDNVLEFVTGHEGNSRKNSSNGGNPYDIDHKKTISSAIINHAFLQNTVMKNCNYKRKRRERDWDCNTKKDVCIPD
ACB42428.1                 QPAGTLDNVLEFVTGHEGNSRKNSSNGGNPYDIDHKKTISSAIINHAFLQNTVMKNCNYKRKRRERDWDCNTKKDVCIPD
ACB42432                   QPAGTLDNVLEFVTGHEGNSRKNSSNGGNPYDIDHKKTISSAIINHAFLQNTVMKNCNHKRKRRERDWDCNTKKDVCIPD
PVL_060021400-t42_1-p1     QPTGTLDNVLEFVTGHGGNSRENSSNGGNPYDIDHKKTISSAIINHAFLQNSVMKNCNDKRKRRERDWDCNTKKDVCIPD
ACB42432.1                 QPAGTLDNVLEFVTGHEGNSRKNSSNGGNPYDIDHKKTISSAIINHAFLQNTVMKNCNHKRKRRERDWDCNTKKDVCIPD
```

### Figure S17 (cont.)

```
PCYB_063270-t26_1-p1        RRYQLCMKEVTNLVNNTDTHFHGDITFRKLDFKRKLMYDAAIEGELLYKKNNYKYNKDLCRDIRWSLGDFGDIIMGTDME
PcyM_0628800-t36_1-p1       RRYQLCMKEVTNLVNNTDTHFHGDITFRKLDFKRKLMYDAAIEGELLYKKNNYKYNKDLCRDIRWSLGDFGDIIMGTDME
PVP01_0623800.1-p1          RRYQLCMKELTNLVNNTDTNFHSDITFRKLYLKRKLIYDAAVEGDLLLKLNNYRYNKDFCKDIRWSLGDFGDIIMGTDME
PVX_110810.1-p1             RRYQLCMKELTNLVNNTDTNFHRDITFRKLYLKRKLIYDAAVEGDLLLKLNNYRYNKDFCKDIRWSLGDFGDIIMGTDME
Psim2302_000333900.1        RRYQLCMKELTNLVNNTDTYFHSDITFRKLYLKRKLMYDAAVEGDLLFKLNNYRYNKDFCKDIRWSLGDFGDIIMGTDME
Psim3636_000376000.1        RRYQLCMKELTNLVNNTDTYFHSDITFRKLYLKRKLMYDAAVEGDLLFKLNNYRYNKDFCKDIRWSLGDFGDIIMGTDME
PsimAD002_000401800.1       RRYQLCMKELTNLVNNTDTYFHSDITFRKLYLKRKLIYDAAVEGDLLFKLNNYRYNKDFCKDIRWSLGDFGDIIMGTDME
PsimAD005_000131600.1       RRYQLCMKELTNLVNNTDTYFHSDITFRKLYLKRKLIYDAAVEGDLLFKLNNYRYNKDFCKDIRWSLGDFGDIIMGTDME
PsimAF28_001016200.1        RRYQLCMKELTNLVNNTDTYFHSDITFRKLYLKRKLIYDAAVEGDLLFKLNNYRYNKDFCKDIRWSLGDFGDIIMGTDME
PsimAF33_000351600.1        RRYQLCMKELTNLVNNTDTYFHSDITFRKLYLKRKLIYDAAVEGDLLFKLNNYRYNKDFCKDIRWSLGDFGDIIMGTDME
PvivaxAM01_000437300.1      RRYQLCMKELTNLVNNTDTNFHRDITFRKLYLKRKLIYDAAVEGDLLFKLNNYRYNKDFCKDIRWSLGDFGDIIMGTDME
PvivaxAM02_000308000.1      RRYQLCMKELTNLVNNTDTNFHRDITFRKLYLKRKLIYDAAVEGDLLLKLNNYRYNKDFCKDIRWSLGDFGDIIMGTDME
AAZ81533.1                  RRYQLCMKELTNLVNNTDTNFHSDITFRKLYLKRKLIYDAAVEGDLLLKLNNYRYNKDFCKDIRWSLGDFGDIIMGTDME
AAZ81530.1                  RRYQLCMKELTNLVNNTDTNFHSDITFRKLYLKRKLIYDAAVEGDLLLKLNNYRYNKDFCKDIRWSLGDFGDIIMGTDME
ABA39301.1                  RRYQLCMKELTNLVNNTDTNFHRDITFRKLYLKRKLIYDAAVEGDLLLKLNNYRYNKDFCKDIRWSLGDFGDIIMGTDME
ACB42428.1                  RRYQLCMKELTNLVNNTDTNLHSDITFRKLYLKRKLIYDAAVEGDLLLKLNNYRYNKDFCKDIRWSLGDFGDIIMGTDME
ACB42432                    RRYQLCMKELTNLVNNTDTYFHSDITFRKLYLKRKLIYDAAVEGDLLFKLNNYRYNKDFCKDIRWSLGDFGDIIMGTDME
PVL_060021400-t42_1-p1      RRYQLCIKELTNLVNNTDTNFHSDITFRKLYLKRKLIYDAAVEGDLLLKLNNYRYNKDFCKDIRWSLGDFGDIIMGTDME
ACB42432.1                  RRYQLCMKELTNLVNNTDTYFHSDITFRKLYLKRKLIYDAAVEGDLLFKLNNYRYNKDFCKDIRWSLGDFGDIIMGTDME

PCYB_063270-t26_1-p1        GIGHSQVVENNLKSIFGTGKNALQNRVQWWHESKVHIWRAMMYSVRKRLKDRFAWICKINVAVNVEPQIYRWIREWGGDY
PcyM_0628800-t36_1-p1       GIGHSQVVENNLKSIFGTGKNALQNRVQWWHESKVHIWRAMMYSVRKRLKDRFAWICKINVAVNVEPQIYRWIREWGGDY
PVP01_0623800.1-p1          GIGYSKVVENNLRSIFGTGEKAQQHRKQWWNESKAQIWTAMMYSVKKRLKGNFIWICKINVAVNIEPQIYRWIREWGRDY
PVX_110810.1-p1             GIGYSKVVENNLRSIFGTDEKAQQRRKQWWNESKAQIWTAMMYSVKKRLKGNFIWICKLNVAVNIEPQIYRWIREWGRDY
Psim2302_000333900.1        GIGYSNVVENNLRSIFGTGQNAQQHRKQWWNESKAQIWRAMMYSVNKRLKGNFRWICKINVAVNIEPQIYRWIREWGRDY
Psim3636_000376000.1        GIGYSNVVENNLRSIFGTGQNAQQHRKQWWNESKAQIWRAMMYSVNKRLKGNFRWICKINVAVNIEPQIYRWIREWGRDY
PsimAD002_000401800.1       GIGYSKVVENNLRSIFGTGKTAQQHRKQWWNESKAQIWRAMMYSVNKRLKGNFIWICKINVAVNIEPQIYRWIREWGRDY
PsimAD005_000131600.1       GIGYSKVVENNLRSIFGTGKNAQQHRKQWWNESKAQIWRAMMYSVNKRLKGNFLWICKINVAVNIEPQIYRWIREWGRDY
PsimAF28_001016200.1        GIGYSKVVENNLRSIFGTGKTAQQHRKQWWNESKAQIWRAMMYSVNKRLKGNFIWICKINVAVNIEPQIYRWIREWGRDY
PsimAF33_000351600.1        GIGYSKVVENNLRSIFGTGQNAQQHRKQWWNESKAQIWRAMMYSVNKRLKGNFIWICKINVAVNIEPQIYRWIREWGRDY
PvivaxAM01_000437300.1      GIGYSKVVEDNLRSIFGTGKNAQQHRKQWWNESKAQIWTAMMYSVKKRLKGKFIWICKINVAVNIEPQIYRRIREWGRDY
PvivaxAM02_000308000.1      GIGYSEVVENNLRSIFGTGEQAQQRRKQWWNESKAQIWTAMMYSVKKRLKGKFIWICKINVAVNIEPQIYRRIREWGRDY
AAZ81533.1                  GIGYSKVVENNLRSIFGTGKNAQQHRKQWWNETKAQIWRAMMYSVKKRLKGNFIWICKINVAVNIEPQIYRWIREWGRDY
AAZ81530.1                  GIGYSEVVENNLRSIFGTGKNAQQRRKQWWNESKAQIWTAMMYSVKKRLKGKFIWICKINVAVNIEPQIYRRIREWGRDY
ABA39301.1                  GIGYSKVVENNLRSIFGTGEKAQQRRKQWWNESKAQIWTAMMYSVKKRLKGKFMWICKINVAVNIEPQIYRRIREWGRDY
ACB42428.1                  GIGYSKVVENNLRSIFGTGKNAQQHRKQWWNETKAQIWRAMMYSVKKRLKGNFIWICKINVAVNIEPQIYRWIREWGRDY
ACB42432                    GIGYSKVVENNLRSIFGTGKNAQQHRKQWWNESKAQIWRAMMYSVNKRLKGNFIWICKINVAVNIEPQIYRWIREWGRDY
PVL_060021400-t42_1-p1      GIGYSKVVENNLRSIFGTGEKAQQHRKQWWNESKAHIWRAMMYSVKKRLKGDFKWICKINVAVNIEPQIYRWIREWGRDY
ACB42432.1                  GIGYSKVVENNLRSIFGTGKNAQQHRKQWWNESKAQIWRAMMYSVNKRLKGNFIWICKINVAVNIEPQIYRWIREWGRDY

PCYB_063270-t26_1-p1        ITELPTERQKVKEKCDGNIIFTKKKVCTV-SQCKNACNLYDQWITRKKEQWDVLSNKFKSVKNEQSIKTADIATAYDILK
PcyM_0628800-t36_1-p1       ITELPTERQKVKEKCDGNIIFTKKKVCTV-SQCKNACNLYDQWITRKKEQWDVLSNKFKSVKNEQSIKTADIATAYDILK
PVP01_0623800.1-p1          VKELPTEVQKLKEKCDGKINYTDKKVCKV-PPCQNACKSYDQWITRKKNQWDVLSNKFISVKNAEKVQTAGIVTPYDILK
PVX_110810.1-p1             VSELPTEVQKLKEKCDGKINYTDKKVCKV-PPCQNACKSYDQWITRKKNQWDVLSNKFISVKNAEKVQTAGIVTPYDILK
Psim2302_000333900.1        VSELPTEVQKLKEKCDGKINYTDKKVCKV-PPCQNACKSYDQWITRKKNQWDVLSNKFKSVKNAEKVQTAGIVTPYDILK
Psim3636_000376000.1        VSELPTEVQKLKEKCDGKINYTDKKVCKV-PPCQNACKSYDQWITRKKNQWDVLSNKFKSVKNAEKVQTAGIVTPYDILK
PsimAD002_000401800.1       VSELPTEVQKLKEKCDGKINYTDKKVCKV-PPCQNACKSYDQWITRKKNQWDVLSNKFISVKNAEKVQTAGIVTPYDILK
PsimAD005_000131600.1       VSELPTEVQKLKEKCDGKINYTDKKVCKV-PPCQNACKSYDQWITRKKNQWDVLSNKFISVKNAEKVQTAGIVTPYDILK
PsimAF28_001016200.1        VSELPTEVQKLKEKCDGKINYTDKKVCKV-PPCQNACKSYDQWITRKKNQWDVLSNKFISVKNAEKVQTAGIVTPYDILK
PsimAF33_000351600.1        VSELPTEVQKLKEKCDGKINYTDKKVCKV-PPCQNACKSYDQWITRKKNQWDVLSNKFKSVKNAEKVQTAGIVTPYDILK
PvivaxAM01_000437300.1      VSELPTEVQKLKEKCDGKINYTDKKVCKV-PPCQNACKSYDQWITRKKNQWDVLSNKFKSVKNAEKVQTAGIVTPYDILK
PvivaxAM02_000308000.1      VSELPTEVQKLKEKCDGKINYTDKKVCKV-PPCQNACKSYDQWITRKKNQWDVLSNKFKSVKNAEKVQTAGIVTPYDILK
AAZ81533.1                  VSELPTEVQKLKEKCDGKINYTDKKVCKVLPPCQNACKSYDQWITRKKNQWDVLSNKFISVKNAEKVQTAGIVTPYDILK
AAZ81530.1                  VSELPTEVQKLKEKCDGKINYTDKKVCKV-PPCQNACKSYDQWITRKKNQWDVLSNKFKSVKNAEKVQTAGIVTPYDILK
ABA39301.1                  VSELPTEVQKLKEKCDGKINYTDKKVCKV-PPRQDACKSYDQWITRKKNQWDVLSNKFISVKNAEKVQTAGIVTPYDILK
ACB42428.1                  VSELPTEVQKLKEKCDGKINYTDKKVCKVLPPCQNACKSYDQWITRKKNQWDVLSNKFISVKNAEKVQTAGIVTPYDILK
ACB42432                    VSELPTEVQKLKEKCDGKINYTDKKVCKV-PPCQNACKSYDQWITRKKNQWDVLSNKFKSVKNAEKVQTAGIVTPYDILK
PVL_060021400-t42_1-p1      VSELPTEVQKLKEKCDGKINYTDKKVCTV-PPCQDACKSYDQWITRKKNQWDVLSNKFISVKSAEKVQTAGIVTPYDILK
ACB42432.1                  VSELPTEVQKLKEKCDGKINYTDKKVCKV-PPCQNACKSYDQWITRKKNQWDVLSNKFKSVKNAEKVQTAGIVTPYDILK
DBL domain                  |<-----------------------------------------------------------------------------
```

Figure S17 (cont.)

```
PCYB_063270-t26_1-p1       QEINKFNEETFENEINKLDNAYIDLCLCSLEEVKKHTQNVVRNIEKAAKSVAPNPNPINKAVDSSKAEKVQGNPENGNVN
PcyM_0628800-t36_1-p1      QEINKFNEETFENEINKLDNAYIDLCLCSLEEVKKHTQNVVRNIEKAAKSVAPNPNPINKAVDSSKAEKVQGNPENGNVN
PVP01_0623800.1-p1         QELDEFNEVAFENEINKRDGAYIELCVCSVEEAKKNTQEVVTNVDNAAKSQATNSNPISQPVDSSKAEKVPGDSTHGNVN
PVX_110810.1-p1            QELDEFNEVAFENEINKRDGAYIELCVCSVEEAKKNTQEVVTNVDNAAKSQATNSNPISQPVDSSKAEKVPGDSTHGNVN
Psim2302_000333900.1       QELDEFNEVAFENEINKRDGAYIELCDCSVEEAKKNTQEVVTNVDNAAKSQATNSNPISQPVDSSKAEKVPGDSTHGNVN
Psim3636_000376000.1       QELDEFNEVAFENEINKRDGAYIELCDCSVEEAKKNTQEVVTNVDNAAKSQATNSNPISQPVDSSKAEKVPGDSTHGNVN
PsimAD002_000401800.1      QELDEFNEVAFENEINKRDGAYIELCDCSVEEAKKNTQEVVTNVDNAAKSQATNSNPISQPVDSSKAEKVPGDSTHGNVN
PsimAD005_000131600.1      QELDEFNEVAFENEINKRDGAYIELCDCSVEEAKKNTQEVVTNVDNAAKSQATNSNPISQPVDSSKAEKVPGDSTHGNVN
PsimAF28_001016200.1       QELDEFNEVAFENEINKRDGAYIELCDCSVEEAKKNTQEVVTNVDNAAKSQATNSNPISQPVDSSKAEKVPGDSTHGNVN
PsimAF33_000351600.1       QELDEFNEVAFENEINKRDGAYIELCDCSVEEAKKNTQEVVTNVDNAAKSQATNSNPISQPVDSSKAEKVPGDSTHGNVN
PvivaxAM01_000437300.1     QELDEFNEVAFENEINKRDGAYIELCVCSVEEAKKNTQEVVTNVDNAAKSQATNSNPISQPVDSSKAEKVPGDSTHGNVN
PvivaxAM02_000308000.1     QELDEFNEVAFENEINKRDGAYIELCVCSVEEAKKNTQEVVTNVDNAAKSQATNSNPISQPVDSSKAKKVPGDSTHGNVN
AAZ81533.1                 QELDEFNEMAFENEINKRDGAYIELCVCSVEEAKKNTQEVVTNVDNAAKSQATNSNPISQPVDSSKAEKVPGDSTHGNVN
AAZ81530.1                 QELDEFNEVAFENEINKRDGAYIELCVCSVEEAKKNTQEVVTNVDNAAKSQATNSNPISQPVDSSKAEKVPGDSTHGNVN
ABA39301.1                 QELDEFNEVAFENEINKRDGAHIELCVCSVEEAKKNTQEVVTNVDNAAKSQATNSNPISQPVDSSKAEKVPGDSTHGNVN
ACB42428.1                 QELDEFNEVAFENEINKRDGAYIELCVCSVEEAKKNTQEVVTNVDNAAKSQATNSNPISQPVDSSKAEKVPGDSTHGNVN
ACB42432                   QELDEFNEVAFENEINKRDGAYIELCVCSVEEAKKNTQEVVTNVDNAAKSQATNSNPISQPVDSSKAEKVPGDSTHGNVN
PVL_060021400-t42_1-p1     QELDEFNEVAFENEINKHDDAYIELCVCSVEEAKKNTQEVVTNVENAAKSQAPNSNPISQPVDSSKAEKVPGDSTHGNVN
ACB42432.1                 QELDEFNEVAFENEINKRDGAYIELCVCSVEEAKKNTQEVVTNVDNAAKSQATNSNPISQPVDSSKAEKVPGDSTHGNVN
DBL domain                 ----------------------------------------->|

PCYB_063270-t26_1-p1       SGANNSTTGKAATGDGQNGNQTPTKSNVQQSDVAESAGAKNVDPHKSVSEKSADTTSVTSIAEAGKENLGTSNSQPSKST
PcyM_0628800-t36_1-p1      SGANNSTTGKAATGDGQNGNQTPTKSNVQQSDVAESAGAKNVDPHKSVSEKSADTTSVTSIAEAGKENLGTSNSQPSKST
PVP01_0623800.1-p1         SGQDSSTTGKAVTGDGQNGNQTPAESDVQRSDIAESVSAKNVDPQKSVSERSDDTASVTGIAEAGKENLGASNSRPSEST
PVX_110810.1-p1            SGQDSSTTGKAVTGDGQNGNQTPAESDVQRSDIAESVSAKNVDPQKSVSKRSDDTASVTGIAEAGKENLGASNSRPSEST
Psim2302_000333900.1       SGQDSSTTGKAVTGDGQNGNQTPAESDVQRSDIAESVSAKNVDPQKSVSERSDDTASVTGIAEAGKENLGASNSRPSEST
Psim3636_000376000.1       SGQDSSTTGKAVTGDGQNGNQTPAESDVQRSDIAESVSAKNVDPQKSVSERSDDTASVTGIAEAGKENLGASNSRPSEST
PsimAD002_000401800.1      SGQDSSTTGKAVTGDGQNGNQTPAESDVQRSDIAESVSAKNVDPQKSVSERSDDTASVTGIAEAGKENLGASNSRPSEST
PsimAD005_000131600.1      SGQDSSTTGKAVTGDGQNGNQTPAESDVQRSDIAESVSAKNVDPQKSVSERSDDTASVTGIAEAGKENLGASNSRPSEST
PsimAF28_001016200.1       SGQDSSTTGKAVTGDGQNGNQTPAESDVQRSDIAESVSAKNVDPQKSVSERSDDTASVTGIAEAGKENLGASNSRPSEST
PsimAF33_000351600.1       SGQDSSTTGKAVTGDGQNGNQTPAESDVQRSDIAESVSAKNVDPQKSVSERSDDTASVTGIAEAGKENLGASNSRPSEST
PvivaxAM01_000437300.1     SGQDSSTTGKAVTGDGKNGNQTPAESDVQRSDIAESVSAKNVDPQKSVSERSDDTASVTGIAEAGKENLGASNSRPSEST
PvivaxAM02_000308000.1     SGQDSSTTGKAVTGDGQNGNQTPAESDVQRSDIAESVSAKNVDPQKSVSERSDDTASVTGIAEAGKENLGASNSRPSEST
AAZ81533.1                 SGQDSSTTGKAVTGDGQNGNQTPAESDVQRSDIAESVSAKNVDPQKSVSERSDDTASVTGIAEAGKENLSASNSRPSEST
AAZ81530.1                 SGQDSSTTGKAVTGDGQNGNQTPAESDVQRSDIAESVSAKNVDPQKSVSERSDDTASVTGIAEAGKENLGASNSRPSEST
ABA39301.1                 SGQDSSTTGKAVTGDGKNGNQTPAESDVQRSDIAESVSAKNVDPQKSVSERSDDTASVTGIAEAGKENLGASNSRPSEST
ACB42428.1                 SGQDSSTTGKAVTGDGQNGNQTPAESDVQRSDIAESVSAKNVDPQKSVSERSDDTASVTGIAEAGKENLGASNSRPSEST
ACB42432                   SGQDSSTTGKAVTGDGQNGNQTPAESDVQRSDIAESVSAKNVDPQKSVSKRSDDTASVTGIAEAGKENLGASNSRPSEST
PVL_060021400-t42_1-p1     SGQDSSTTGKAVTGDGQNGNQTPAESDVQRSDIAESVSAKNVDPQKSVSERSDDTASVTGVTESGKENLGASNSPPSEPT
ACB42432.1                 SGQDSSTTGKAVTGDGQNGNQTPAESDVQRSDIAESVSAKNVDPQKSVSKRSDDTASVTGIAEAGKENLGASNSRPSEST

PCYB_063270-t26_1-p1       VEANSPGDGTVNSASISVKNSEKPLVTTDKGLEPSKDNSDNNGSTE------SKKSEANPDSNSKGETGMGQDNDKEKAT
PcyM_0628800-t36_1-p1      VEANSPGDGTVNSASISVKNSEKPLVTTDKGLEPSKDNSDNNGSTE------SKKSEANPDSNSKGETGMGQDNDKEKAT
PVP01_0623800.1-p1         VEANSPGDDTVNSASIPVVSGENPLVTPYNGLGHSKDNSDSDGPAEFAESTKSAESMANPDSNSKGETGKGQDNDMAKAT
PVX_110810.1-p1            VEANSPGDDTVNSASIPVVSGENPLVTPYNGLRHSKDNSDSDGP---------AESMANPDSNSKGETGKGQDNDMAKAT
Psim2302_000333900.1       VEANSPGDDTVNSASIPVVSGKNPLVTPYNGL--------------------------------------GQDNDMAKAT
Psim3636_000376000.1       VEANSPGDDTVNSASIPVVSGKNPLVTPYNGL--------------------------------------GQDNDMAKAT
PsimAD002_000401800.1      VEANSPGDDTVNSASIPVVSGKNPLVTPYNGL--------------------------------------GQDNDMAKAT
PsimAD005_000131600.1      VEANSPGDDTVNSASIPVVSGKNPLVTPYNGL--------------------------------------GQDNDMAKAT
PsimAF28_001016200.1       VEANSPGDDTVNSASIPVVSGKNPLVTPYNGL--------------------------------------GQDNDMAKAT
PsimAF33_000351600.1       VEANSPGDDTVNSASIPVVSGKNPLVTPYNGL--------------------------------------GQDNDMAKAT
PvivaxAM01_000437300.1     VEANSPGDDTVNSASIPVVSGENPLVTPYNGLRHSKDNSDSDGP---------AESMANPDSNSKGETGKGQDNDMAKAT
PvivaxAM02_000308000.1     VEANSPGDDTVNSASIPVVSGENPLVTPYNGLRHSKDNSDSDGP---------AESMANPDSNSKGETGKGQDNDMAKAT
AAZ81533.1                 VEANSPGDDTVNSASIPVVSGENPLVTPYNGLRHSKDNSDSDGP---------AESMANPDSNSKGETGKGQDNDMAKAT
AAZ81530.1                 VEANSPGDDTVNSASIPVVSGENPLVTPYNGLRHSKDNSDSDGP---------AESMANPDSNSKGETGKGQDNDMAKAT
ABA39301.1                 VEANSPGDDTVNSASIPVVSGENPLVTPYNGLRHSKDNSDSDGP---------AESMANPDSNSKGETGKGQDNDMAKAT
ACB42428.1                 VEANSPGDDTVNSASIPVVSGENPLVTPYNGLRHSKDNSDSDGP---------AESMANPDSNSKGETGKGQDNDMAKAT
ACB42432                   VEANSPGDDTVNSASIPVVSGENPLVTPYNGLRHSKDNSDSDGP---------AESMANPDSNSKGETGKGQDNDMAKAT
PVL_060021400-t42_1-p1     VEANSPGDGTVNSASIPVVSGENPLVTPDNGLGHSKDNSDSDGPAEFAESMKSVESMANPDSNSKGETGKGQDNDMAKAT
ACB42432.1                 VEANSPGDDTVNSASIPVVSGENPLVTPYNGLRHSKDNSDSDGP---------AESMANPDSNSKGETGKGQDNDMAKAT
```

Figure S17 (cont.)

```
PCYB_063270-t26_1-p1        KDSSNSSDNTSSAKGDTTSAVDRNFNGGVPEDRDKIVGSKKEEKEDNSANKDAATVV----------GGNTNDRTENGME
PcyM_0628800-t36_1-p1       KDSSNSSDNTSSAKGDTTSAVDRNFNGGVPEDRDKIVGSKKEEKEDNSANKDAATVV----------GGNTNDRTENGME
PVP01_0623800.1-p1          KDSSNSSDGTSSATGDTTDAVDREINKGVPEDRDKTVGSKDGGGEDNSANKDAATVVGEDRIRENSAGGSTNDRSKNDTE
PVX_110810.1-p1             KDSSNSSDGTSSATGDTTDAVDREINKGVPEDRDKTVGSKDGGGEDNSANKDAATVVGEDRIRENSAGGSTNDRSKNDTE
Psim2302_000333900.1        KDSSNSSDGTSSATGDTTDAVDREINKGVPEDRDKTVGSKDGGGEDNSANKDAATVVGEDRIRENSAGGSTNDRSKNDTE
Psim3636_000376000.1        KDSSNSSDGTSSATGDTTDAVDREINKGVPEDRDKTVGSKDGGGEDNSANKDAATVVGEDRIRENSAGGSTNDRSKNDTE
PsimAD002_000401800.1       KDSSNSSDGTSSATGDTTDAVDREINKGVPEDRDKTVGSKDGGGEDNSANKDAATVVGEDRIRENSAGGSTNDRSKNDTE
PsimAD005_000131600.1       KDSSNSSDGTSSATGDTTDAVDREINKGVPEDRDKTVGSKDGGGEDNSANKDAATVVGEDRIRENSAGGSTNDRSKNDTE
PsimAF28_001016200.1        KDSSNSSDGTSSATGDTTDAVDREINKGVPEDRDKTVGSKDGGGEDNSANKDAATVVGEDRIRENSAGGSTNDRSKNDTE
PsimAF33_000351600.1        KDSSNSSDGTSSATGDTTDAVDREINKGVPEDRDKTVGSKDGGGEDNSANKDAATVVGEDRIRENSAGGSTNDRSKNDTE
PvivaxAM01_000437300.1      KDSSNSSDGTSSATGDTTDAVDREINKGVPEDRDKTVGSKDGGGEDNSANKDAATVVGEDRIRENSAGGSTNDRSKNDTE
PvivaxAM02_000308000.1      KDSSNSSDGTSSARGDTTDAVDREINKGVPEDRDKTVGSKDGGGEDNSANKDAATVVGEDRIRENSAGGSTNDRSKNDTE
AAZ81533.1                  KDSSNSSDGTSSATGDTTDAVDREINKGVPEDRDKTVGSKDGGGEDNSANKDAATVVGEDRIRENSAGGSTNDRSKNDTE
AAZ81530.1                  KDSSNSSDGTSSATGDTTDAVDREINKGVPEDRDKTVGSKDGGGEDNSANKDAATVVGEDRIRENSAGGSTNDRSKNDTE
ABA39301.1                  KDSSNSSDGTSSATGDTTDAVDREINKGVPEDRDKTVGSKDGGGEDNSANKDAATVVGEDRIRENSAGGSTNDRSKNDTE
ACB42428.1                  KDSSNSSDGTSSATGDTTDAVDREINKGVPEDRDKTVGSKDGGGEDNSANKDAATVVGEDRIRENSAGGSTNDRSKNDTE
ACB42432                    KDSSNSSDGTSSATGDTTDAVDREINKGVPEDRDKTVGSKDGGGEDNSANKDAATVVGEDRIRENSAGGSTNDRSKNDTE
PVL_060021400-t42_1-p1      KDSSNSSDSTNSATGDATGAVDREINKGVPEDSDKTVGSKEGGGEDNSANKDAAVVVGEDRIRENSAGGSINDRSKNDTE
ACB42432.1                  KDSSNSSDGTSSATGDTTDAVDREINKGVPEDRDKTVGSKDGGGEDNSANKDAATVVGEDRIRENSAGGSTNDRSKNDTE


PCYB_063270-t26_1-p1        KNNVPAPDSKQSVDATPLSKTESLELNESAHRITNDTTHSLKNENEGSEKDLQKHDFTNNDMPNEEPNSAQTTDAEGHHR
PcyM_0628800-t36_1-p1       KNNVPAPDSKQSVDATPLSKTESLELNESAHRITNDTTHSLKNENEGSEKDLQKHDFTNNDMPNEEPNSAQTTDAEGHHR
PVP01_0623800.1-p1          KNGASTPDSKQSEDATALSKTESLESTESGDRTTNDTTNSLENKNGGKEKDLQKHDFKSNDTPNEEPNSDQTTDVEGHDR
PVX_110810.1-p1             KNGASTPDSKQSEDATALSKTESLESTESGDRTTNDTTNSLENKNGGKEKDLQKHDFKSNDTPNEEPNSDQTTDAEGHDR
Psim2302_000333900.1        KNGASTPDSKQSEDATALSKTESLESTESGDRTTNDTTNSLENKNGGKEKDLQKHDFKSNDTPNEEPNSDQTTDAEGHDR
Psim3636_000376000.1        KNGASTPDSKQSEDATALSKTESLESTESGDRTTNDTTNSLENKNGGKEKDLQKHDFKSNDTPNEEPNSDQTTDAEGHDR
PsimAD002_000401800.1       KNGASTPDSKQSEDATALSKTESLESTESGDRTTNDTTNSLENKNGGKEKDLQKHDFKSNDTPNEEPNSDQTTDAEGHDR
PsimAD005_000131600.1       KNGASTPDSKQSEDATALSKTESLESTESGDRTTNDTTNSLENKNGGKEKDLQKHDFKSNDTPNEEPNSDQTTDAEGHDR
PsimAF28_001016200.1        KNGASTPDSKQSEDATALSKTESLESTESGDRTTNDTTNSLENKNGGKEKDLQKHDFKSNDTPNEEPNSDQTTDAEGHDR
PsimAF33_000351600.1        KNGASTPDSKQSEDATALSKTESLESTESGDRTTNDTTNSLENKNGGKEKDLQKHDFKSNDTPNEEPNSDQTTDAEGHDR
PvivaxAM01_000437300.1      KNGASTPDSKQSEDATALSKTESLESTESGDRTTNDTTNSLENKNGGKEKDLQKHDFKSNDTPNEEPNSDQTTDAEGHDR
PvivaxAM02_000308000.1      KNGASTPDSKQSEDATALSKTESLESTESGDRTTNDTTNSLENKNGGKEKDLQKHDFKSNDTPNEEPNSDQTTDAEGHDR
AAZ81533.1                  KNGASTPDSKQSEDATALSKTESLESTESGDRTTNDTTISLENKNGGKEKDLQKHDFKSNDTPNEEPNSDQTTDAEGHDR
AAZ81530.1                  KNGASTPDSKQSEDATALSKTESLESTESGDRTTNDTTNSLENKNGGKEKDLQKHDFKSNDTPNEEPNSDQTTDAEGHDR
ABA39301.1                  KNGASTPDSKQSEDATALSKTESLESTESGDRTTNDTTNSLENKNGGKEKDLQKHDFKSNDTPNEEPNSDQTTDAEGHDR
ACB42428.1                  KNGASTPDSKQSEDATALSKTESLESTESGDRTTNDTTNSLENKNGGKEKDLQKHDFKSNDTPNEEPNSDQTTDVEGHDR
ACB42432                    KNGASTPDSKQSEDATALSKTESLESTESGDRTTNDTTNSLENKNGGKEKDLQKHDFKSNDTPNEEPNSDQTTDAEGHDR
PVL_060021400-t42_1-p1      KNGAPTPDSKQSEDATALSKTESLESTESVDRTTTDTTNSLENKNGGKEKDLQKHDFKSNDTPNEEPNSDQTTDAEGPDR
ACB42432.1                  KNGASTPDSKQSEDATALSKTESLESTESGDRTTNDTTNSLENKNGGKEKDLQKHDFKSNDTPNEEPNSDQTTDAEGHDR


PCYB_063270-t26_1-p1        YSMKNDNGEMRKHMNKGTFTKNPNSNQLNSHNDLSNGKLDIKEYKYRDVNATREKIIYMSEVRRCNNNISLNYCNSVKDK
PcyM_0628800-t36_1-p1       YSMKNDNGEMRKHMNKGTFTKNPNSNQLNSHNDLSNGKLDIKEYKYRDVNATREKIIYMSEVRRCNNNISLNYCNSVKDK
PVP01_0623800.1-p1          DSIKNDKAERRKHMNKDTFTKNTNSHHLNSNNNLSNGKLDIKEYKYRDVKATRENIILMSSVRKCNNNISLEYCNSVEDK
PVX_110810.1-p1             DSIKNDKAERRKHMNKDTFTKNTNSHHLNSNNNLSNGKLDIKEYKYRDVKATREDIILMSSVRKCNNNISLEYCNSVEDK
Psim2302_000333900.1        DSIKNDKAERRKHMNKDTFTKNTNSHHLNSNNNLSNGKLDIKEYKYRDVKATRENIILMSSVRKCNNNISLKYCNSVEDK
Psim3636_000376000.1        DSIKNDKAERRKHMNKDTFTKNTNSHHLNSNNNLSNGKLDIKEYKYRDVKATRENIILMSSVRKCNNNISLKYCNSVEDK
PsimAD002_000401800.1       DSIKNDKAERRKHMNKDTFTKNTNSHHLNSNNNLSNGKLDIKEYKYRDVKATRENIILMSSVRKCNNNISLKYCNSVEDK
PsimAD005_000131600.1       DSIKNDKAERRKHMNKDTFTKNTNSHHLNSNNNLSNGKLDIKEYKYRDVKATRENIILMSSVRKCNNNISLEYCNSVEDK
PsimAF28_001016200.1        DSIKNDKAERRKHMNKDTFTKNTNSHHLNSNNNLSNGKLDIKEYKYRDVKATRENIILMSSVRKCNNNISLKYCNSVEDK
PsimAF33_000351600.1        DSIKNDKAERRKHMNKDTFTKNTNSHHLNSNNNLSNGKLDIKEYKYRDVKATRENIILMSSVRKCNNNISLKYCNSVEDK
PvivaxAM01_000437300.1      DSIKNDKAERRKHMNKDTFTKNTNSHHLNSNNNLSNGKLDIKEYKYRDVKATRENIILMSSVRKCNNNISLEYCNSVEDK
PvivaxAM02_000308000.1      DSIKNDKAERRKHMNKDTFTKNTNSHHLNSNNNLSNGKLDIKEYKYRDVKATRENIILMSSVRKCNNNISLEYCNSVEDK
AAZ81533.1                  DSIKNDKAERRKHMNKDTFTKNTNSHHLNSNNNLSNGKLDIKEYKYRDVKATRENIILMSSVRKCSNNISLEYCNSVEDK
AAZ81530.1                  DSIKNDKAERRKHMNKDTFTKNTNSHHLNSNNNLSNGKLDIKEYKYRDVKATREDIILMSSVRKCNNNISLEYCNSVEDK
ABA39301.1                  DSIKNDKAERRKHMNKDTFTKNTNSHHLNSNNNLSNGKLDIKEYKYRDVKATREDIILMSSVRKCNNNISLEYCNSVEDK
ACB42428.1                  DSIKNDKAERRKHMNKDTFTKNTNSHHLNSNNNLSNGKLDIKEYKYRDVKATRENIILMSSVRKCNNNISLEYCNSVEDK
ACB42432                    DSIKNDKAERRKHMNKDTFTKNTNSHHLNSNNNLSNGKLDIKEYKYRDVKATREDIILMSSVRKCNNNISLEYCNSVEDK
PVL_060021400-t42_1-p1      DGIKNDKAERRKHMNKDTFTKNQNSHHLNSNNNLGNGKLDIKEYEYRDVNATREKIILMSAVRKCNNNISLKYCNSVEDK
ACB42432.1                  DSIKNDKAERRKHMNKDTFTKNTNSHHLNSNNNLSNGKLDIKEYKYRDVKATREDIILMSSVRKCNNNISLEYCNSVEDK
```

### Figure S17 (cont.)

```
PCYB_063270-t26_1-p1      MSSNTCSREKSKNLCCSISDYCLNYFEVYTNEYHNCMKKEFEDPSYKCFTKGGFTDKAYFAAGGALLILLLLITSRHMIK
PcyM_0628800-t36_1-p1     MSSNTCSREKSKNLCCSISDYCLNYFEVYTNEYHNCMKKEFEDPSYKCFTKGGFTDKAYFAAGGALLILLLLITSRHMIK
PVP01_0623800.1-p1        ISSNTCSREKSKNLCCSISDFCLNYFDVYSYEYHSCMKKEFEDPSYKCFTKGGFKDKTYFAAAGALLILLLLIASRKMIK
PVX_110810.1-p1           ISSNTCSREKSKNLCCSISDFCLNYFDVYSYEYLSCMKKEFEDPSYKCFTKGGFKDKTYFAAAGALLILLLLIASRKMIK
Psim2302_000333900.1      ISSNTCSREKSKNLCCSISDFCLNYFDVHSYEYLSCMKKEFEDPSYKCFTKGGFKDKAYFAAAGALLILLLLIASRKMIK
Psim3636_000376000.1      ISSNTCSREKSKNLCCSISDFCLNYFDVHSYEYLSCMKKEFEDPSYKCFTKGGFKDKAYFAAAGALLILLLLIASRKMIK
PsimAD002_000401800.1     ISSNTCSREKSKNLCCSISDFCLNYFDVHSYEYLSCMKKEFEDPSYKCFTKGGFKDKAYFAAAGALLILLLLIASRKMIK
PsimAD005_000131600.1     ISSNTCSREKSKNLCCSISDFCLNYFDVHSYEYLSCMKKEFEDPSYKCFTKGGFKDKTYFAAAGALLILLLLIASRKMIK
PsimAF28_001016200.1      ISSNTCSREKSKNLCCSISDFCLNYFDVHSYEYLSCMKKEFEDPSYKCFTKGGFKDKAYFAAAGALLILLLLIASRKMIK
PsimAF33_000351600.1      ISSNTCSREKSKNLCCSISDFCLNYFDVHSYEYLSCMKKEFEDPSYKCFTKGGFKDKAYFAAAGALLILLLLIASRKMIK
PvivaxAM01_000437300.1    ISSNTCSREKSKNLCCSISDFCLNYFDVNSYEYHSCMKKEFEDPSYKCFTKGGFKDKTYFAAAGALLILLLLIASRKMIK
PvivaxAM02_000308000.1    ISSNTCSREKSKNLCCSISDFCLNYFDVYSYEYLSCMKKEFEDPSYKCFTKGGFKDKTYFAAAGALLILLLLIASRKMIK
AAZ81533.1                ISSNTCSREKSKNLCCSISDFCLNYFDVYSYEYLSCMKKEFEDPSYKCFTKGGFKDKTYFAAAGALLILLLLIASRKMIK
AAZ81530.1                ISSNTCSREKSKNLCCSISDFCLNYFDVNSYEYHSCMKKEFEDPSYKCFTKGGFKDKTYFAAAGALLILLLLIASRKMIK
ABA39301.1                ISSNTCSREKSKNLCCSISDFCLNYFDVYSYEYLSCMKKEFEDPSYKCFTKGGFKDKTYFAAAGALLILLLLIASRKMIK
ACB42428.1                ISSNTCSREKSKNLCCSISDFCLNYFDVNSYEYHSCVKKEFEDPSYKCFTKGGFKDKTYFAAAGALLILLLLIASRKMIK
ACB42432                  ISSNTCSREKSKNLCCSISDFCLNYFDVHSYEYLSCMKKEFEDPSYKCFTIGGFKDKTYFAAAGALLILLLLIASRKMIK
PVL_060021400-t42_1-p1    ISSNTCSREKSKNLCCSISDFCLNYFDVYSFEYHNCMKKEFEDPSYKCFTKGGFTDNTYFAAAGALLILLLLIASRKMIK
ACB42432.1                ISSNTCSREKSKNLCCSISDFCLNYFDVHSYEYLSCMKKEFEDPSYKCFTIGGFKDKTYFAAAGALLILLLLIASRKMIK
Trans-membrane domain                                                                |<-------------->|

PCYB_063270-t26_1-p1      NDSEEATFNEFEEHCDNIQRIPLMPNNIEHMQPLTPLDYS
PcyM_0628800-t36_1-p1     NDSEEATFNEFEEHCDNIQRIPLMPNNIEHMQPLTPLDYS
PVP01_0623800.1-p1        NDSEEATFNEFEEYCDNIHRIPLMPNNIEHMQPSTPLDYS
PVX_110810.1-p1           NDSEEATFNEFEEYCDNIHRIPLMPNNIEHMQPSTPLDYS
Psim2302_000333900.1      NDSEEATFNEFEEYCDNIHRIPLMPNNIEHMQPSTPLDYS
Psim3636_000376000.1      NDSEEATFNEFEEYCDNIHRIPLMPNNIEHMQPSTPLDYS
PsimAD002_000401800.1     NDSEEATFNEFEEYCDNIHRIPLMPNNIEHMQPSTPLDYS
PsimAD005_000131600.1     NDSEEATFNEFEEYCDNIHRIPLMPNNIEHMQPSTPLDYS
PsimAF28_001016200.1      NDSEEATFNEFEEYCDNIHRIPLMPNNIEHMQPSTPLDYS
PsimAF33_000351600.1      NDSEEATFNEFEEYCDNIHRIPLMPNNIEHMQPSTPLDYS
PvivaxAM01_000437300.1    NDSEEATFNEFEEYCDNIHRIPLMPNNIEHMQPSTPLDYS
PvivaxAM02_000308000.1    NDSEEATFNEFEEYCDNIHRIPLMPNNIEHMQPSTPLDYS
AAZ81533.1                NDSEEATFNEFEEYCDNIHRIPLM----------------
AAZ81530.1                NDSEEATFNEFEEYCDNIHRIPLM----------------
ABA39301.1                NDSEEATFNAFEEYCDNIHRIPLM----------------
ACB42428.1                NDSEEATFKEFEEYCDNIHRIPLMPNNIEHMQPSTPLDYS
ACB42432                  NDSEEATFNEFEEYCDNIHRIPLMPNNIEHMQPSTPLDYS
PVL_060021400-t42_1-p1    NDSEEATFNEFEEYCDNIHRIPLMPNNIEHMQPSTPLDYS
ACB42432.1                NDSEEATFNEFEEYCDNIHRIPLMPNNIEHMQPSTPLDYS
```

Figure S18

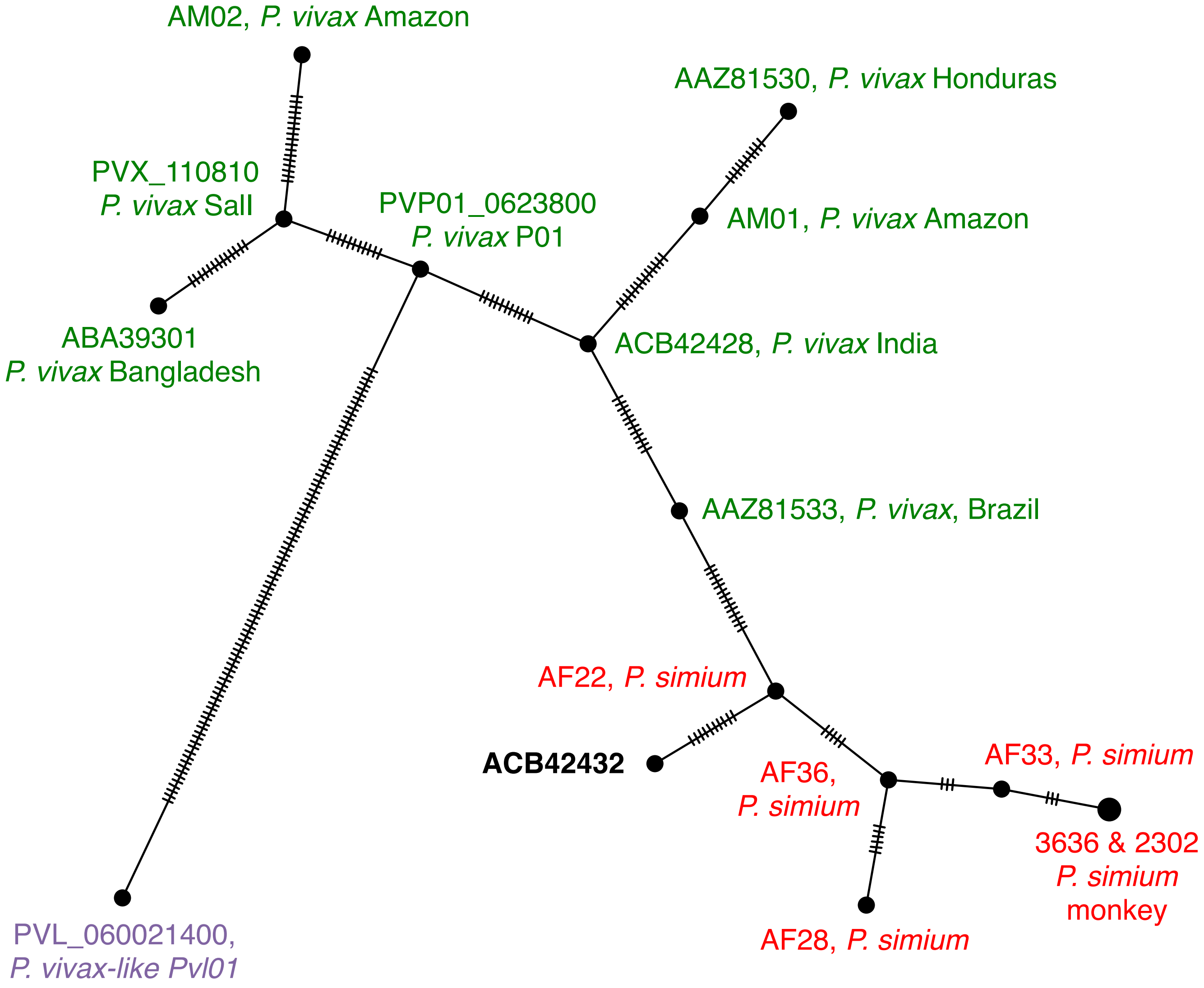
AM02, *P. vivax* Amazon
AAZ81530, *P. vivax* Honduras
PVX_110810
*P. vivax* SalI
PVP01_0623800
*P. vivax* P01
AM01, *P. vivax* Amazon
ABA39301
*P. vivax* Bangladesh
ACB42428, *P. vivax* India
AAZ81533, *P. vivax*, Brazil
AF22, *P. simium*
ACB42432
AF36,
*P. simium*
AF33, *P. simium*
3636 & 2302
*P. simium*
monkey
AF28, *P. simium*
PVL_060021400,
*P. vivax-like Pvl01*

Figure S19

```
simium          GGTAACCCCCTATAATGGTTT------------------------------------------------------------------------------------------------------------------GGGGCAAGATAATGAT
vivax 1, P01    GGTAACCCCCTATAATGGTTTGGGGCATTCGAAAGACAATAGTGATAGCGATGGACCTGCGGAATTTGCAGAATCTACGAAATCTGCGGAATCAATGGCGAATCCTGATTCAAATAGTAAAGGTGAGACGGGAAAGGGGCAAGATAATGAT
vivax 2, SalI   GGTAACCCCCTATAATGGTTTGGGGCATTCGAAAGACAATAGTGATAGCGATGGAC----------------------------CTGCGGAATCAATGGCGAATCCTGATTCAAATAGTAAAGGTGAGACGGGAAAGGGGCAAGATAATGAT
vivax 3,        GGTAACCCCCTATAATGGTTTGGGGCATTCGAAAGACAATAGT------GATGGACCTGCGGAATTTGCAGAATCTACGAAATCTGCGGAATCAATGGCGAATCCTGATTCAAATAGTAAAGGTGAGACGGGAAAGGGGCAAGATAATGAT
vivax 4         GGTAACCCCCTATAATGGTTTGGGGCATTCGAAAGACAATAGT------GATGGAC----------------------------CTGCGGAATCAATGGCGAATCCTGATTCAAATAGTAAAGGTGAGACGGGAAAGGGGCAAGATAATGAT
```

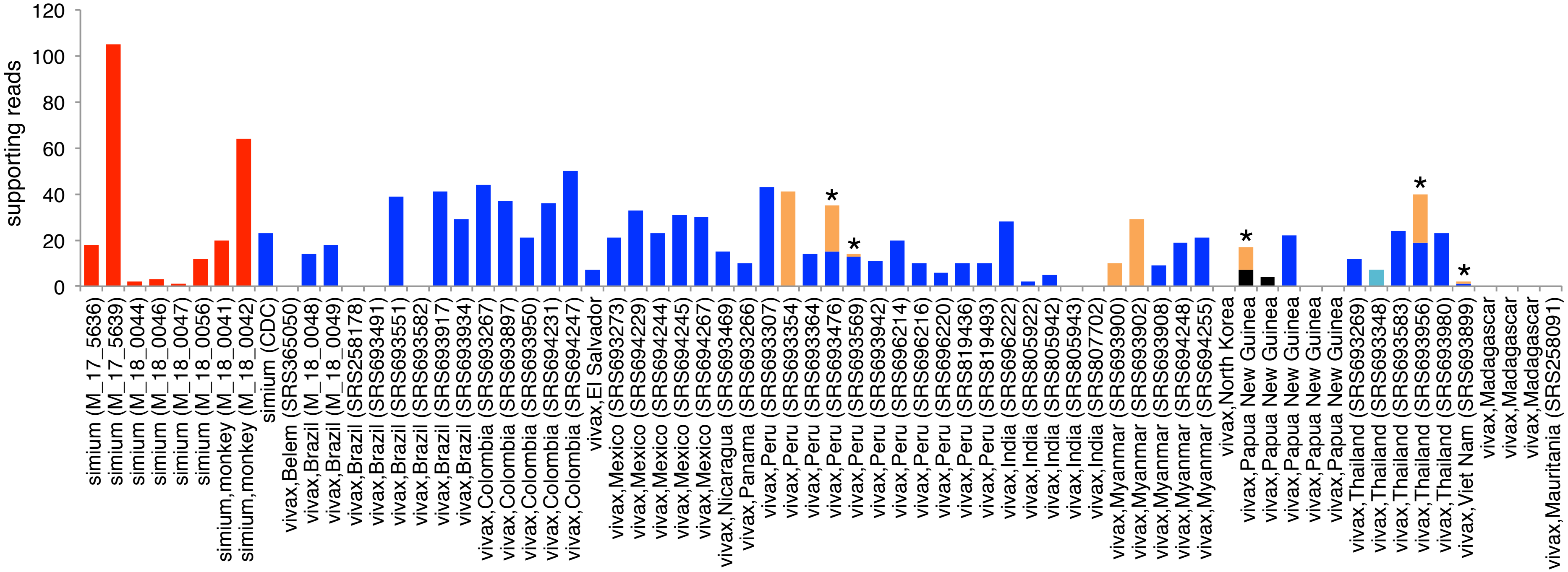

Figure S20

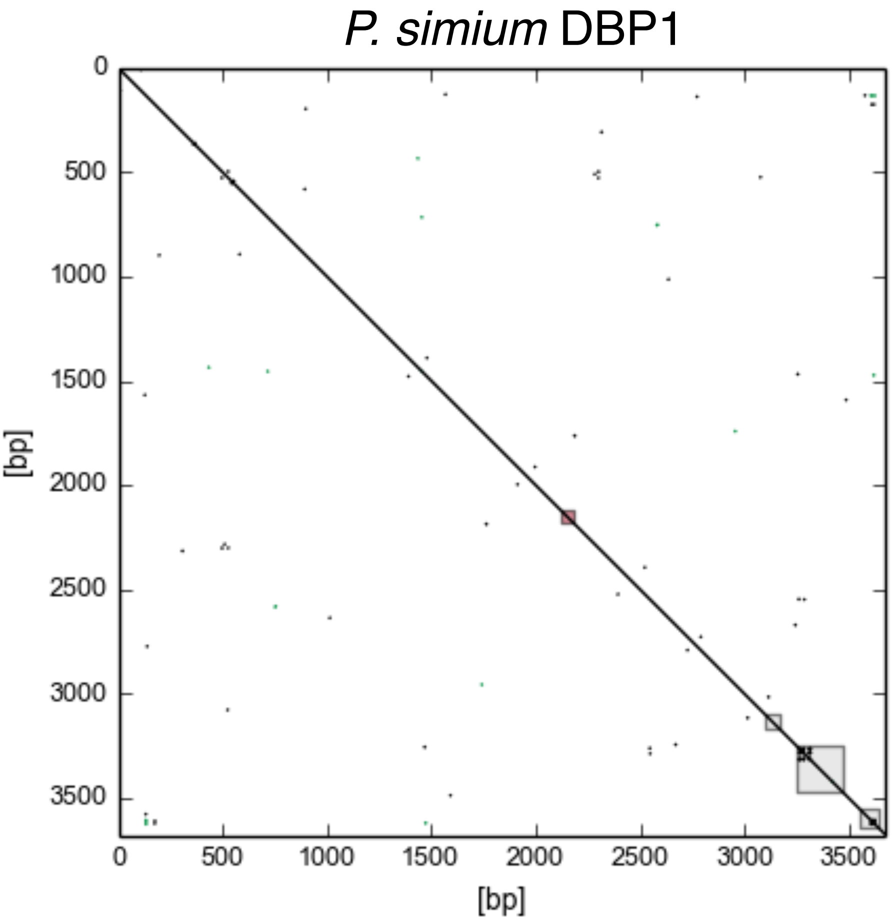
P. simium DBP1
0
500
1000
1500
2000
2500
3000
3500
[bp]
0
500
1000
1500
2000
2500
3000
3500
[bp]

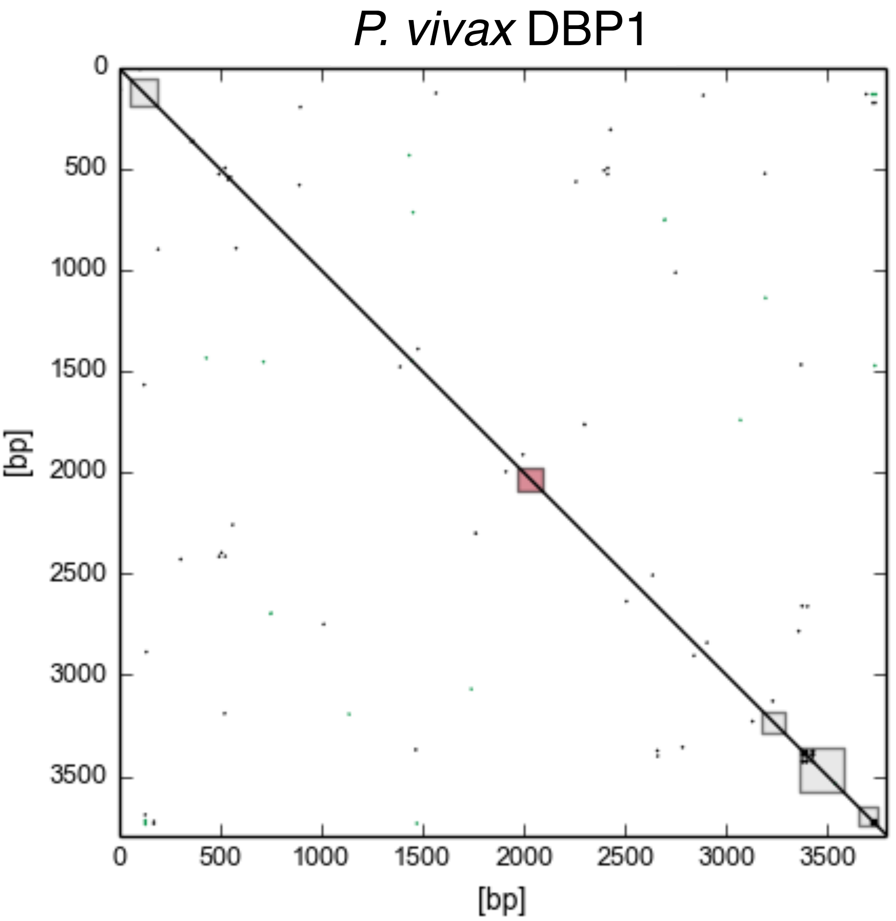
P. vivax DBP1
0
500
1000
1500
2000
2500
3000
3500
[bp]
0
500
1000
1500
2000
2500
3000
3500
[bp]

Figure S21

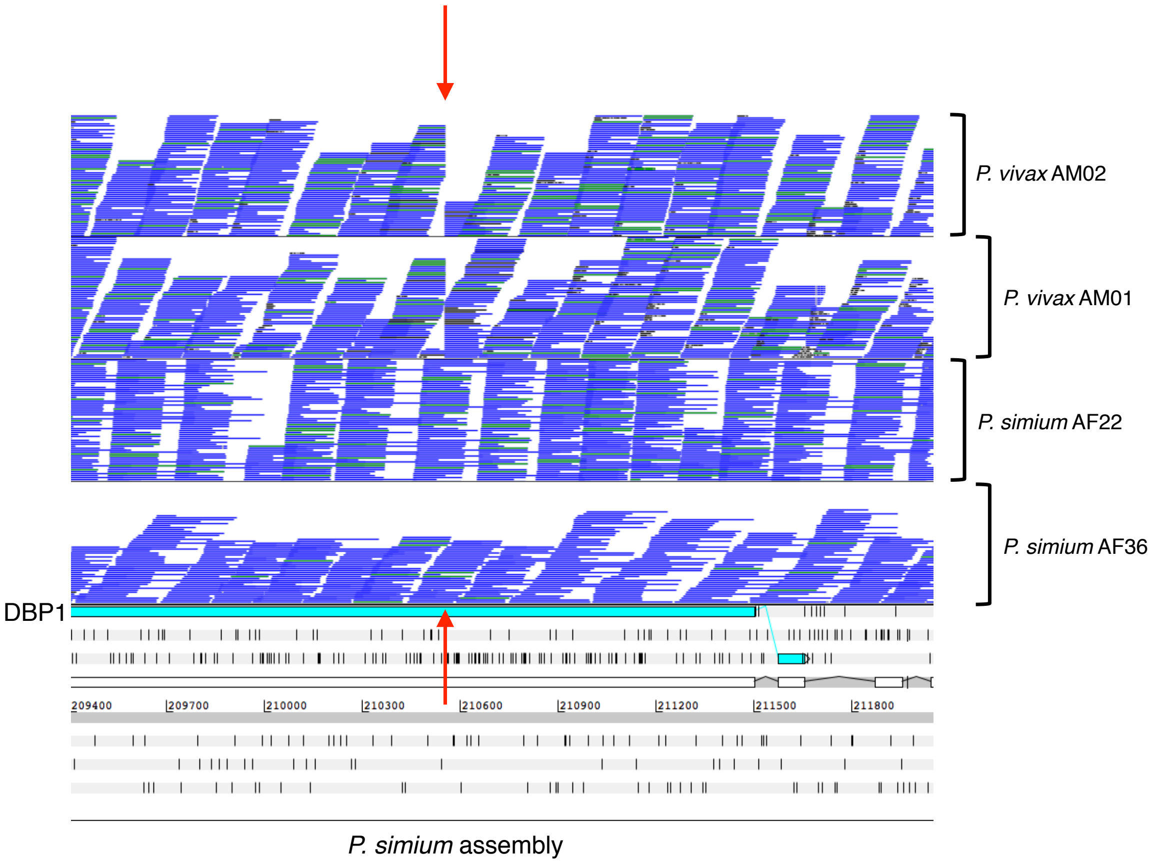

P. vivax AM02
P. vivax AM01
P. simium AF22
P. simium AF36
DBP1
209400
209700
210000
210300
210600
210900
211200
211500
211800
P. simium assembly

Figure S22

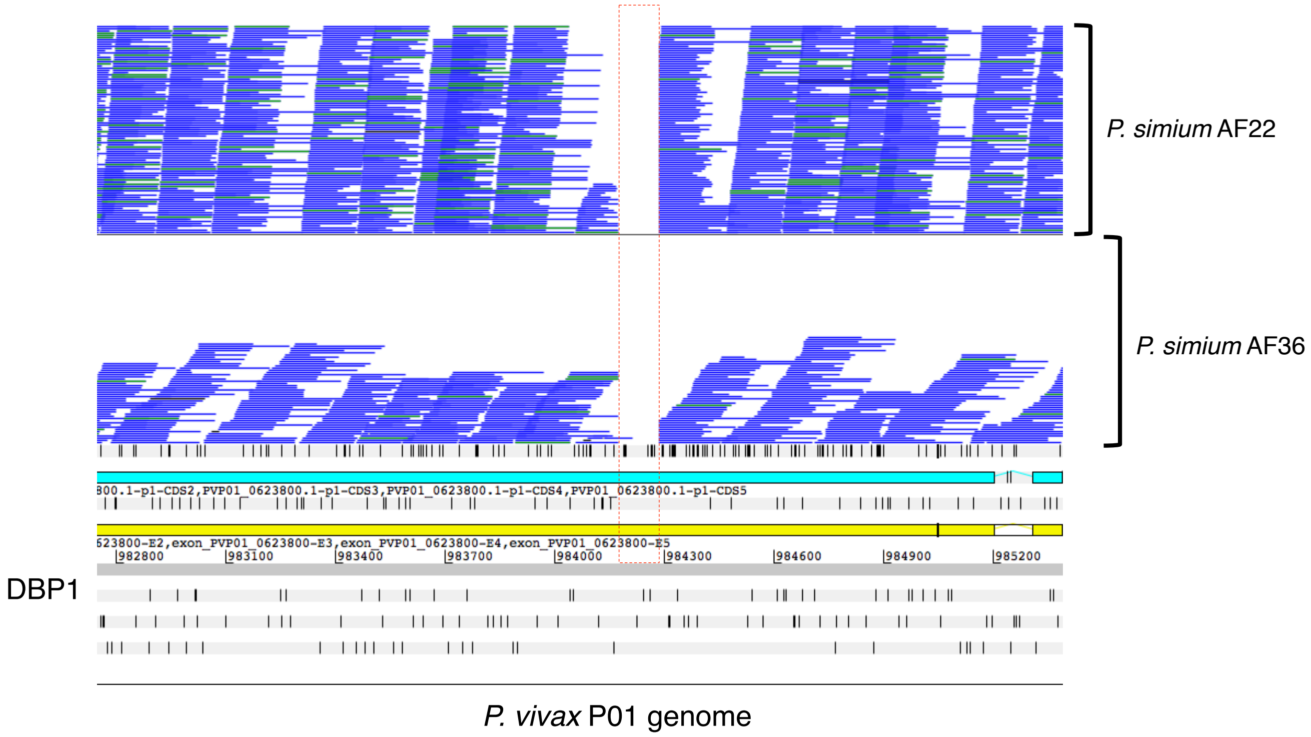

Figure S23

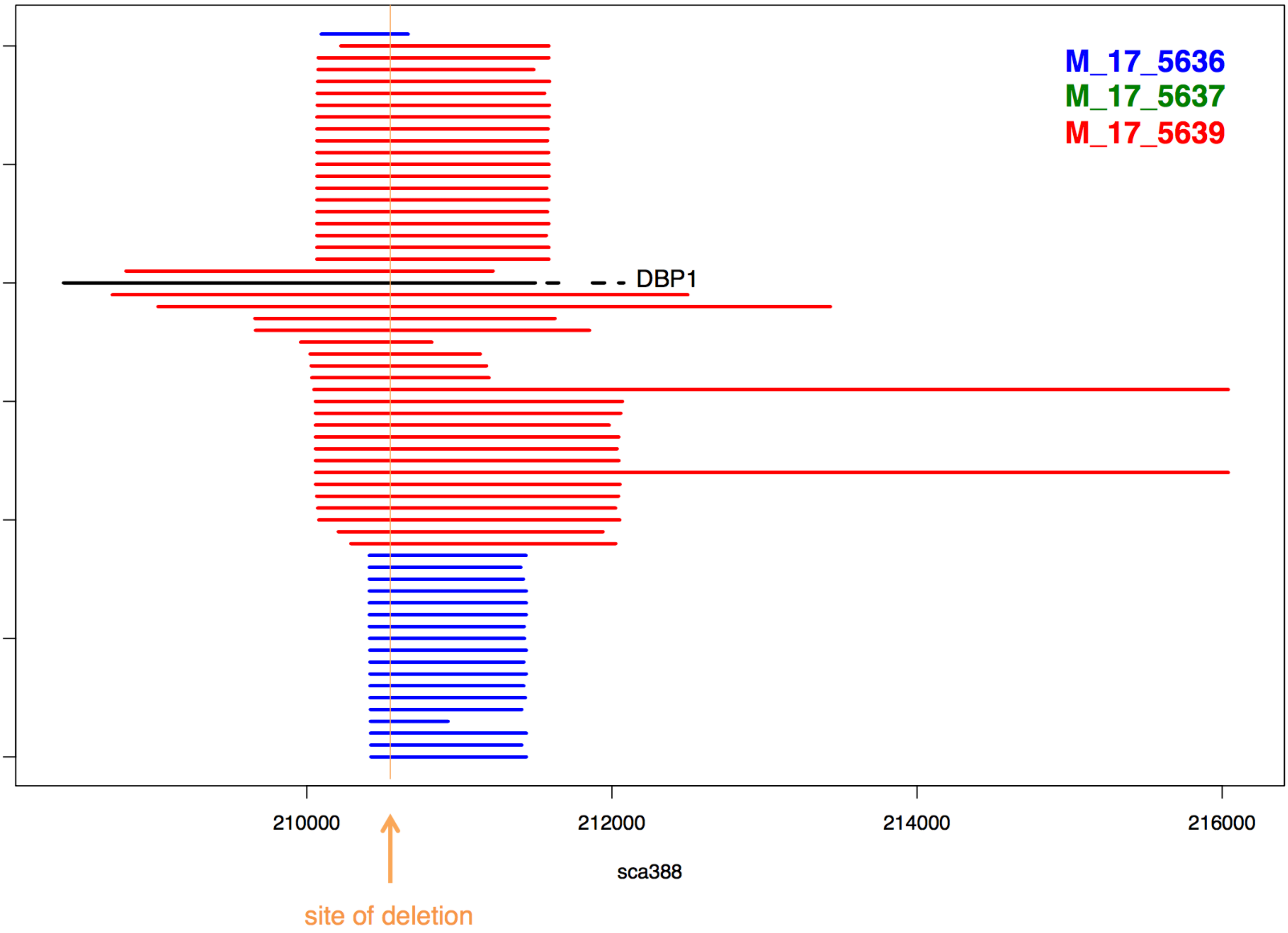

M_17_5636
M_17_5637
M_17_5639
DBP1
210000
212000
214000
216000
sca388
site of deletion

Figure S24

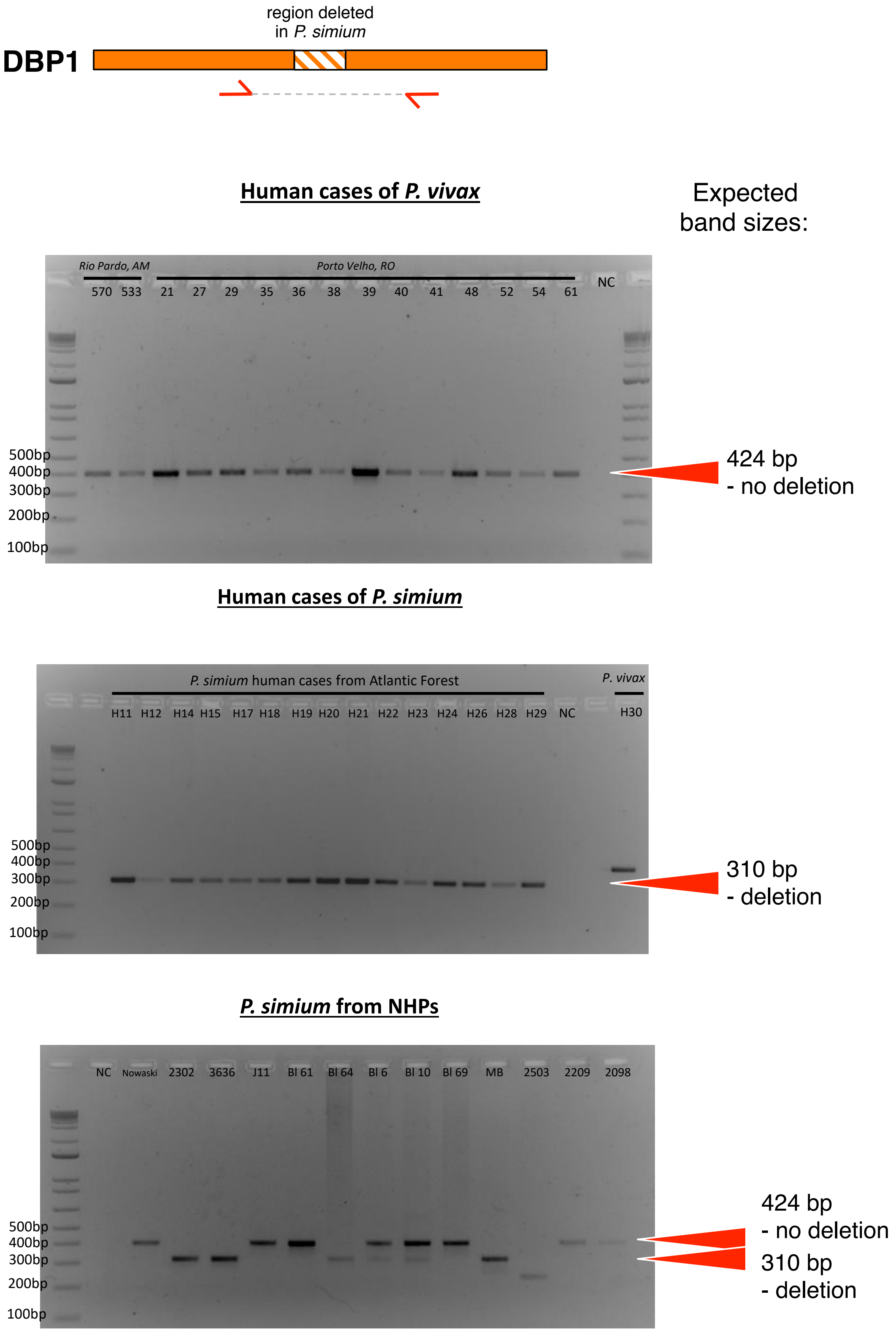


forward primer: ATCGCGACAAAGCTACTGCC
reverse primer: TCCCACAACAATCCCTGCTG

### Figure S25

**Complete alignment of RBP2a protein sequences**

```
PVX_121920.1-p1            MENKVLWAVFYNLVLFLLASSKESNRIKAYKLKKEPKLWPLQDSLNESDKFEYTNGGKENPPNFFSSNVKAHNKKEGKKY
PVP01_1402400.1-p1         MENKVLWAVFYNLVLFLLASSKESNRIKAYKLKKEPKLWPLQDSLNESDKFEYTNGGKENPPNFFSSNVKAHNKKEGKKY
Psim2302_000007000.1       MENKVLWAVFYNLVLFLLASSKESNRIKAYKLKKEPKLWPLQDSLNESDKFEYTNGGKENPPNFFSSNVKAHNKKEGKKY
Psim3636_000006300.1       MENKVLWAVFYNLVLFLLASSKESNRIKAYKLKKEPKLWPLQDSLNESDKFEYTNGGKENPPNFFSSNVKAHNKKEGKKY
PsimAD002_000007200.1      MENKVLWAVFYNLVLFLLASSKESNRIKAYKLKKEPKLWPLQDSLNESDKFEYTNGGKENPPNFFSSNVKAHNKKEGKKY
PsimAD005_000448800.1      MENKVLWAVFYNLVLFLLASSKESNRIKAYKLKKEPKLWPLQDSLNESDKFEYTNGGKENPPNFFSSNVKAHNKKEGKKY
PsimAF28_000006200.1       MENKVLWAVFYNLVLFLLASSKESNRIKAYKLKKEPKLWPLQDSLNESDKFEYTNGGKENPPNFFSSNVKAHNKKEGKKY
PsimAF33_000006700.1       MENKVLWAVFYNLVLFLLASSKESNRIKAYKLKKEPKLWPLQDSLNESDKFEYTNGGKENPPNFFSSNVKAHNKKEGKKY
PsimAF34_000005900.1       MENKVLWAVFYNLVLFLLASSKESNRIKAYKLKKEPKLWPLQDSLNESDKFEYTNGGKENPPNFFSSNVKAHNKKEGKKY
PvivaxAM01_000076800.1     MENKVLWAVFYNLVLFLLASSKESNRIKAYKLKKEPKLWPLQDSLNESDKFEYTNGGKENPPNFFSSNVKAHNKKEGKKY
PvivaxAM02_000474900.1     MENKVLWAVFYNLVLFLLASSKESNRIKAYKLKKEPKLWPLQDSLNESDKFEYTNGGKENPPNFFSSNVKAHNKKEGKKY
PVL_140006600-t42_1-p1     MENKVLWAVFYNLVLFLLASSKESNRIKAYKLKKESKLWPLQDSLNESDKFEYTNGGKENPPNFFNSNVKAHNKKEGKKY

PVX_121920.1-p1            EQNLSLPDNTSFVTVKNYNYTRTPSHHAYIRRDNTHTNSTNNNQIRNVPKELNPREFLFTPKNQISASLIQTNGPVAPMD
PVP01_1402400.1-p1         EQNLSLPDNTSFVTVKNYNYTRTPSHHAYIRRDNTHTNSTNNNQIRNVPKELNPREFLFTPKNQISASLIQTNGPVAPMD
Psim2302_000007000.1       EQNLSLPDNTSFVTVKNYNYTRTPSHHAYIRRDNTHTNSTNNNQIRNVPKELNPREFLFTPKNQISASLIQTNKPVAPMD
Psim3636_000006300.1       EQNLSLPDNTSFVTVKNYNYTRTPSHHAYIRRDNTHTNSTNNNQIRNVPKELNPREFLFTPKNQISASLIQTNKPVAPMD
PsimAD002_000007200.1      EQNLSLPDNTSFVTVKNYNYTRTPSHHAYIRRDNTHTNSTNNNQIRNVPKELNPREFLFTPKNQISASLIQTNKPVAPMD
PsimAD005_000448800.1      EQNLSLPDNTSFVTVKNYNYTRTPSHHAYIRRDNTHTNSTNNNQIRNVPKELNPREFLFTPKNQISASLIQTNKPVAPMD
PsimAF28_000006200.1       EQNLSLPDNTSFVTVKNYNYTRTPSHHAYIRRDNTHTNSTNNNQIRNVPKELNPREFLFTPKNQISASLIQTNKPVAPMD
PsimAF33_000006700.1       EQNLSLPDNTSFVTVKNYNYTRTPSHHAYIRRDNTHTNSTNNNQIRNVPKELNPREFLFTPKNQISASLIQTNKPVAPMD
PsimAF34_000005900.1       EQNLSLPDNTSFVTVKNYNYTRTPSHHAYIRRDNTHTNSTNNNQIRNVPKELNPREFLFTPKNQISASLIQTNGPVAPMD
PvivaxAM01_000076800.1     EQNLSLPDNTSFVTVKNYNYTRTPSHHAYIRRDNTHTNSTNNNQIRNVPKELNPREFLFTPKNQISASLIQTNGPVAPMD
PvivaxAM02_000474900.1     EQNLSLPDNTSFVTVKNYNYTRTPSHHAYIRRDNTHTNSTNNNQIRNVPKELNPREFLFTPKNQISASLIQTNGPVAPMD
PVL_140006600-t42_1-p1     EQNLSLPNNTSFVTVKNYNYTRTPSHHAYIRRDNTHTNSTNNNQIRNVPKELNPREFLFTPNNQISASLIQTNGPVAPMD

PVX_121920.1-p1            ILRYLDFSNSSGQIISTVYPFYVQMNYFAEIKYYITYHYEAKKNYDEAYNQSVNPLMSSIQNQINSCVPKKAALEKTIFV
PVP01_1402400.1-p1         ILRYLDFSNSSGQIISTVYPFYVQMSYFAEIKYYITYHYEAKKNYDEAYNQSVNPLMSRIQNQINSCVPKKAALEKTIFV
Psim2302_000007000.1       ILRYLDFSNSSGQIISTVYPFYVQMNYFAEIKYYITYHYEAKKKYDEAYNQSVNPLMSSIQNQINSCVPKKAALEKTIFV
Psim3636_000006300.1       ILRYLDFSNSSGQIISTVYPFYVQMNYFAEIKYYITYHYEAKKNYDEAYNQSVNPLMSSIQNQINSCVPKKAALEKTIFV
PsimAD002_000007200.1      ILRYLDFSNSSGQIISTVYPFYVQMNYFAEIKYYITYHYEAKKNYDEAYNQSVNPLMSSIQNQINSCVPKKAALEKTIFV
PsimAD005_000448800.1      ILRYLDFSNSSGQIISTVYPFYVQMNYFAEIKYYITYHYEAKKNYDEAYNQSVNPLMSSIQNQINSCVPKKAALEKTIFV
PsimAF28_000006200.1       ILRYLDFSNSSGQIISTVYPFYVQMNYFAEIKYYITYHYEAKKKYDEAYNQSVNPLMSSIQNQINSCVPKKAALEKTIFV
PsimAF33_000006700.1       ILRYLDFSNSSGQIISTVYPFYVQMNYFAEIKYYITYHYEAKKNYDEAYNQSVNPLMSSIQNQINSCVPKKAALEKTIFV
PsimAF34_000005900.1       ILRYLDFSNSSGQIISTVYPFYVQMNYFAEIKYYITYHYEAKKNYDEEYNQSVNPLMSSIQNQINSCVPKKAALEKTIFV
PvivaxAM01_000076800.1     ILRYLDFSNSSGQIISTVYPFYVQMSYFAEIKYYITYHYEAKKNYDEAYNQSVNPLMSSIQNQINSCVPKKAALEKTIFV
PvivaxAM02_000474900.1     ILRYLDFSNSSGQIISTVYPFYVQMSYFAEIKYYITYHYEAKKNYDEAYNQSVNPLMSSIQNQINSCVPKKAALEKTIFV
PVL_140006600-t42_1-p1     ILRYLDFSNSSGQIISTVYPFYVQMNYFAEIKYYITFHFEAKKNYDEAYNQSVNPLMSSIQNQINSCVPKKAALEKTIFV

PVX_121920.1-p1            LEYPENHNINLSNYEAKHNEYKQQLDAYKNCVQANMESYTDRMSKFNEKIYSILNSVKCTDACETDTYEIMLEIYVERVK
PVP01_1402400.1-p1         LEYPENHNINLSNYEAKHNEYKQQLDAYKNCVQANMKSYTDRMSNFNEKIYSILNSVKCTDACETDTYEIMLEIYVERVK
Psim2302_000007000.1       LEYPENHNINLSNYEAKHNEYKQQLDAYKNCVQANMESYTDRMSEFNEKIYSILNSVKCTDACETVTYKIMVEIYVERVN
Psim3636_000006300.1       LEYPENHNINLSNYEAKHNEYKQQLDAYKNCVQANMESYTDRMSEFNEKIYSILNSVKCTDACETVTYKIMVEIYVERVN
PsimAD002_000007200.1      LEYPENHNINLSNYEAKHNEYKQQLDAYKNCVQANMESYTDRMSKFNEKIYSILNSVKCTDACKTVMYKIMLEIYVERVK
PsimAD005_000448800.1      LEYPENHNINLSNYEAKHNEYKQQLDAYKNCVQANMESYTDRMSKFNEKIYSILNSVKCTDACKTVMYKIMLEIYVERVK
PsimAF28_000006200.1       LEYPENHNINLSNYEAKHNEYKQQLDAYKNCVQANMESYTDRMSKFNEKIYSILNSVKCTDACKTVMYKIMLEIYVERVK
PsimAF33_000006700.1       LEYPENHNINLSNYEAKHNEYKQQLDAYKNCVQANMESYTDRMSKFNEKIYSILNSVKCTDACKTVMYKIMLEIYVERVK
PsimAF34_000005900.1       LEYPENHNINLSNYEAKHNEYKQQLDAYKNCVQANMESYTDRMSKFNEKIYSILNSVKCTDACKTVMYKIMLEIYVERVK
PvivaxAM01_000076800.1     LEYPENHNINLSNYEAKHNEYKQQLDAYKNCVQANMKSYTDRMSKFNEEIYSILNSVKCTDACKTVTYEIMLEIYVERVK
PvivaxAM02_000474900.1     LEYPENHNINLSNYEAKHNEYKQQLDAYKNCVQANMKSYTDRMSKFNEEIYSILNSVKCTDACKTVTYEIMLEIYVERVK
PVL_140006600-t42_1-p1     LENPESHNINLSNYEEKHNEYKQQLDAYKNCVQANMESYTDRMSKFNEKIYSILNSVKCTDACETDTYEIMLEIYVERVK

PVX_121920.1-p1            EVNHNNYVNYLSTLKASLQLGVTLMLKVKQEIDNNVTISAINFLQEEMLDIITIGEAHTGKIIHGKENVLKLQNNNIPPQ
PVP01_1402400.1-p1         EVNHNNYVNYLSTLKASLQLGVTLMLKVKQEIDNNVTISAINFLQEEMLDIITIGEAHTGKIIHGKENVLKLQNNNIPSQ
Psim2302_000007000.1       EVNHNNYVNYLSTLKASLQLGVTLMLKVKQEIDNNVTISAINFLQEEMLDIITIGEAHTGKIIHGKENVLKLQNNNIPPK
Psim3636_000006300.1       EVNHNNYVNYLSTLKASLQLGVTLMLKVKQEIDNNVTISAINFLQEEMLDIITIGEAHTGKIIHGKENVLKLQNNNIPPK
PsimAD002_000007200.1      EVNHNNYVNYLSTLKASLQLGVTLMLKVKQEIDNNVTISAINFLQEEMLDIITIGEAHTGKIIHGKENVLKLQNNNIPPK
PsimAD005_000448800.1      EVNHNNYVNYLSTLKASLQLGVTLMLKVKQEIDNNVTISAINFLQEEMLDIITIGEAHTGKIIHGKENVLKLQNNNIPPK
PsimAF28_000006200.1       EVNHNNYVNYLSTLKASLQLGVTLMLKVKQEIDNNVTISAINFLQEEMLDIITIGEAHTGKIIHGKENVLKLQNNNIPPK
PsimAF33_000006700.1       EVNHNNYVNYLSTLKASLQLGVTLMLKVKQEIDNNVTISAINFLQEEMLDIITIGEAHTGKIIHGKENVLKLQNNNIPPK
PsimAF34_000005900.1       EVNHNNYVNYLSTLKASLQLGVTLMLKVKQEIDNNVTISAINFLQEEMLDIITIGEAHTGKIIHGKENVLKLQNNNIPPK
PvivaxAM01_000076800.1     EVNHNNYVNYLSTLKASLQLGVTLMLKVKQEIDNNVTISAINFLQEEMLDIITIGEAHTGKIIHGKENVLKLQNNNIPSQ
PvivaxAM02_000474900.1     EVNHNNYVNYLSTLKASLQLGVTLMLKVKQEIDNNVTISAINFLQEEMLDIITIGEAHTGKIIHGKENVLKLQNNNIPSQ
PVL_140006600-t42_1-p1     EVNLNNYVNYLSTLKASLQFGVTLMLKVKQEIDNNVTISTINFLQEEMLDIITICEAHTGKIIHGKENVLKLQGKEIPAE

PVX_121920.1-p1            VPLSTLKKLYFDSANFYATYKFSLKRADTTTAALKEKGKLLANLYNKLITYVSEKIDKNLDSLYFISKSESMISEFEDTF
PVP01_1402400.1-p1         VPLSTLKKLYFDSANFYATYKFSLKRADTTTAALKEKEKLLANLYNKLITYVSEKIDKNLDSLYFISKSESMISEFEDTF
Psim2302_000007000.1       VPLSTLKKLYFDSANFYATYKFSLKRADTTTAALKEKGILLANLYNKLITYVSEKIDKNLDSLYFISKSESMISEFEDTF
Psim3636_000006300.1       VPLSTLKKLYFDSANFYATYKFSLKRADTTTAALKKKRKLLRNLYKKLITYVSEKIDKNLDSLYFISKSESMISEFEDTF
PsimAD002_000007200.1      VPLSTLKKLYFDSANFYATYKFSLKRADTTTAALKKKRKLLRNLYKKLITYVSEKIDKNLDSLYFISKSESMISEFEDTF
PsimAD005_000448800.1      VPLSTLKKLYFDSANFYATYKFSLKRADTTTAALKKKRKLLRNLYKKLITYVSEKIDKNLDSLYFISKSESMISEFEDTF
PsimAF28_000006200.1       VPLSTLKKLYFDSANFYATYKFSLKRADTTTAALKKKRKLLRNLYKKLITYVSEKIDKNLDSLYFISKSESMISEFEDTF
PsimAF33_000006700.1       VPLSTLKKLYFDSANFYATYKFSLKRADTTTAALKEKGILLANLYNKLITYVSEKIDKNLDSLYFISKSESMISEFEDTF
PsimAF34_000005900.1       VPLSTLKKLYFDSANFYATYKFSLKRADTTTAALKEKGILLANLYNKLITYVSEKIDKNLDSLYFISKSESMISEFEDTF
PvivaxAM01_000076800.1     VPLSTLKKLYFDSANFYATYKFSLKRADTTTAALKEKGKLLANLYNKLITYVSEKIDKNLDSLYFISKSESMISEFEDTF
PvivaxAM02_000474900.1     VPLSTLKKLYFDSANFYATYKFSLKRADTTTAALKEKGKLLANLYNKLITYVSEKIDKNLDSLYFISKSESMISEFEDTF
PVL_140006600-t42_1-p1     VPLSTLKKLYFDSANFYATYKFSLKRTDTTTAALREKGKLLADLYNKLITYVSEKIDKNLDSSYFISKSESMISEFEDTF
```

### Figure S25 (cont.)

```
PVX_121920.1-p1            KAGTEASKENSSITADYARYSNLQIIDIKSKYDYKITELKKYLNQLKVLISPMKDLYNLNSLQKDQRDSVRSIRSSDSSP
PVP01_1402400.1-p1         KAGTEASKENSSITADYARYSNLQIIDIKSMYDYKITELKKYLNQLKVLISPMKDLYNLNSLQKDQRDSVRSIRSSNSSP
Psim2302_000007000.1       KAGTEASKENSSITADYARYSNLQIIDIKSKYDYKITELKKYLNQLKVLISPMKDLYNLNSLQKDQRDSVRSIRSSNSSP
Psim3636_000006300.1       KAGTEASKENSSITADYARYSNLQIIDIKSKYDYKITELKKYLNQLKVLISPMKDLYNLNSLQKDQRDSVRSIRSSNSSP
PsimAD002_000007200.1      KAGTEASKENSSITADYARYSNLQIIDIKSKYDYKITELKKYLNQLKVLISPMKDLYNLNSLQKDQRDSVRSIRSSNSSP
PsimAD005_000448800.1      KAGTEASKENSSITADYARYSNLQIIDIKSKYDYKITELKKYLNQLKVLISPMKDLYNLNSLQKDQRDSVRSIRSSNSSP
PsimAF28_000006200.1       KAGTEASKENSSITADYARYSNLQIIDIKSKYDYKITELKKYLNQLKVLISPMKDLYNLNSLQKDQRDSVRSIRSSNSSP
PsimAF33_000006700.1       KAGTEASKENSSITADYARYSNLQIIDIKSKYDYKITELKKYLNQLKVLISPMKDLYNLNSLQKDQRDSVRSIRSSNSSP
PsimAF34_000005900.1       KAGTEASKENSSITADYARYSNLQIIDIKSKYDYKITELKKYLNQLKVLISPMKDLYNLNSLQKDQRDSVRSIRSSNSSP
PvivaxAM01_000076800.1     KAGTEASKENSSITADYARYSNLQIIDIKSKYDYKITELKKYLNQLKVLISPMKDLYNLNSLQKDQRDSVRSIRSSDSSP
PvivaxAM02_000474900.1     KAGTEASKENSSITADYARYSNLQIIDIKSKYDYKITELKKYLNQLKVLISPMKDLYNLNSLQKDQRDSVRSIRSSDSSP
PVL_140006600-t42_1-p1     KAGTEASKENPSITADYARYSNLQIIDIKSKYDYKITELKKYLNQLKVLISPMKDLYNLNSLQKDQRDSGRSIRSSNSSP

PVX_121920.1-p1            QEKVSALLNFLSTIKEENGTVLENFKNIERYYEQGKPLKGTLEGLIKMIKTYEQDIVNLIDNEKHNRSYKDDIKNKIIYI
PVP01_1402400.1-p1         QEKVSALLNFLSKIKEENGTVLENFKNIERYYEQGKPLKGTLEGLIKMIKTYEQDIVNLIDNEKHNRSYKDDIKNKIIYI
Psim2302_000007000.1       QEKVSALLNFLSKIKEEYGTVLENFKNIERYYEQGKPLKGTLEGLIKMIKAYEQDIVNLIDNEKHNRSYKDDIKNKIIYI
Psim3636_000006300.1       QEKVSALLNFLSTIKKEYGTVLENFKNIERYYEQGKPLKGTLEGLIKMIKAYEQDIVNLIDNEKHNRSYKDDIKNKIIYI
PsimAD002_000007200.1      QEKVSALLNFLSTIKKEYGTVLENFKNIERYYEQGKPLKGTLEGLIKMIKAYEQDIVNLIDNEKHNRSYKDDIKNKIIYI
PsimAD005_000448800.1      QEKVSALLNFLSTIKKEYGTVLENFKNIERYYEQGKPLKGTLEGLIKMIKAYEQDIVNLIDNEKHNRSYKDDIKNKIIYI
PsimAF28_000006200.1       QEKVSALLNFLSTIKKEYGTVLENFKNIERYYEQGKPLKGTLEGLIKMIKAYEQDIVNLIDNEKHNRSYKDDIKNKIIYI
PsimAF33_000006700.1       QEKVSALLNFLSKIKEEYGTVLENFKNIERYYEQGKPLKGTLEGLIKMIKAYEQDIVNLIDNEKHNRSYKDDIKNKIIYI
PsimAF34_000005900.1       QEKVSALLNFLSKIKEEYGTVLENFKNIERYYEQGKPLKGTLEGLIKMIKAYEQDIVNLIDNEKHNRSYKDDIKNKIIYI
PvivaxAM01_000076800.1     QEKVSALLNFLSTIKEENGTVLENFKNIERYYEQGKPLKGTLEGLIKMIKTYEQDIVNLIDNEKHNRSYKDDIENKIIYI
PvivaxAM02_000474900.1     QEKVSALLNFLSTIKEENGTVLENFKNIERYYEQGKPLKGTLEGLIKMIKAYEQDIVNLIDNEKHNRSYKDDIENKIIYI
PVL_140006600-t42_1-p1     QKKVSALINLLSTIKDEDGTVLENFKNIERYYEQGKPLKGTLEGLIKMIKAYEQDIVNLIDNEKHNRSYKDDIKNKIKYI

PVX_121920.1-p1            TEKTDAIKDIIPLNEEVDNIIKQIEKLFNEDLFHEGLPNMEKLKKEKEKIKATIKSLNTNFYDGDLKELVDDMLNFVYKH
PVP01_1402400.1-p1         TEKTDAIKDIIPLNEEVDNIIKQIEKLFNEDLFHEGLPNMEKLKKEKEKIKATIKSLNTNFYDGDLKELVDDMLNFVYKH
Psim2302_000007000.1       TEKTDAIKDIIPLNEEVDNIIKQIEKLFNEDLFHEGLPNMEKLKKEKEKIKATIKSLNTNFYDGDLKELVDDMLNFVYKH
Psim3636_000006300.1       TEKTDAIKDIIPLNEEVDNIIKQIEKLFNEDLFHEGLPNMEKLKKEKEKIKATIKSLNTNFYDGDLKELVDDMLNFVYKH
PsimAD002_000007200.1      TEKTDAIKDIIPLNEEVDNIIKQIEKLFNEDLFHEGLPNMEKLKKEKEKIKATIKSLNTNFYDGDLKELVDDMLNFVYKH
PsimAD005_000448800.1      TEKTDAIKDIIPLNEEVDNIIKQIEKLFNEDLFHEGLPNMEKLKKEKEKIKATIKSLNTNFYDGDLKELVDDMLNFVYKH
PsimAF28_000006200.1       TEKTDAIKDIIPLNEEVDNIIKQIEKLFNEDLFHEGLPNMEKLKKEKEKIKATIKSLNTNFYDGDLKELVDDMLNFVYKH
PsimAF33_000006700.1       TEKTDAIKDIIPLNEEVDNIIKQIEKLFNEDLFHEGLPNMEKLKKEKEKIKATIKSLNTNFYDGDLKELVDDMLNFVYKH
PsimAF34_000005900.1       TEKTDAIKDIIPLNEEVDNIIKQIEKLFNEDLFHEGLPNMEKLKKEKEKIKATIKSLNTNFYDGDLKELVDDMLNFVYKH
PvivaxAM01_000076800.1     TEKTDAIKDIIPLNEEVDNIIKQIEKLFNEDLFHEGLPNMEKLKKEKEKIKATIKSLNTNFYDGDLKELVDDMLNFVYKH
PvivaxAM02_000474900.1     TEKTDAIKDIIPLNEEVDNIIKQIEKLFNEDLFHEGLPNMEKLKKEKEKIKATIKSLNTNFYDGDLKELVDDMLNFVYKH
PVL_140006600-t42_1-p1     TEKTDAIKDIIPLNEEVDNIIKQIEKLFNEDLFHEGLPNMEKLKKEKEKIKATIKSLNTNFYDGDLKELVDDMLNFVYKH

PVX_121920.1-p1            SDFYNQAYAKGDVVNLQKEAEQEYEKITKLKSNNIPQMLKDLKSESKNLSELKDTMMDEMLNNVHQDISNVFEQIRNKYK
PVP01_1402400.1-p1         SDFYDQAYAKGDVVNLQKEAEQEYEKITKLKSNNIPQMLKDLKSESKNLSELKDTMMDEMLNNVHQDISNVFEQIRNKYK
Psim2302_000007000.1       SDFYDQAYDKGDVVNLQKEAEQEYEKITKLKSNNIPQMLKDLKSESKNLSELKDTMMDEMLNNVHQDISNVFEQIRNKYK
Psim3636_000006300.1       SDFYDQAYDKGDVVNLQKEAEQEYEKITKLKSNNIPQMLKDLKSESKNLSELKDTMMDEMLNNVHQDISNVFEQIRNKYK
PsimAD002_000007200.1      SDFYDQAYDKGDVVNLQKEAEQEYEKITKLKSNNIPQMLKDLKSESKNLSELKDTMMDEMLNNVHQDISNVFEQIRNKYK
PsimAD005_000448800.1      SDFYDQAYDKGDVVNLQKEAEQEYEKITKLKSNNIPQMLKDLKSESKNLSELKDTMMDEMLNNVHQDISNVFEQIRNKYK
PsimAF28_000006200.1       SDFYDQAYDKGDVVNLQKEAEQEYEKITKLKSNNIPQMLKDLKSESKNLSELKDTMMDEMLNNVHQDISNVFEQIRNKYK
PsimAF33_000006700.1       SDFYDQAYDKGDVVNLQKEAEQEYEKITKLKSNNIPQMLKDLKSESKNLSELKDTMMDEMLNNVHQDISNVFEQIRNKYK
PsimAF34_000005900.1       SDFYDQAYDKGDVVNLQKEAEQEYEKITKLKSNNIPQMLKDLKSESKNLSELKDTMMDEMLNNVHQDISNVFEQIRNKYK
PvivaxAM01_000076800.1     SDFYDQAYAKGDVVNLQKEAEQEYEKITKLKSNNIPQMLKDLKSESKNLSELKDTMMDEMLNNVHQDISNVFEQIRNKYK
PvivaxAM02_000474900.1     SDFYDQAYAKGDVVNLQKEAEQEYEKITKLKSNNIPQMLKDLKSESKNLSELKDTMMDEMLNNVHQDISNVFEQIRNKYK
PVL_140006600-t42_1-p1     SDFYDQAYAKGDVVNLQKEAEQEYEKITKLKSNNIPQILKDLKSESKNLSELKDTMMDEMLNNVHQDISNAFEQIRNKYK

PVX_121920.1-p1            DITTSLSAYKEDKQKFEKFKVDMIQTKNELLGSLQQNDKKLSEANNMYEEFLAHIDTTTNREKQMYNQANALKENLANVQ
PVP01_1402400.1-p1         DITTSLSAYKEDKQKFEKFKVDMIQTKNELLGSLQQNDKKLSEANNMYEEFLAHIDTTTNREKQMYNQANALKENLANVQ
Psim2302_000007000.1       DITTSLSAYKEDKQKFEKFKVDMIQTKNELVGSLQQNDKKLSEANNMYEEFLAHIDTTTNREKQMYNQANALKENLANVQ
Psim3636_000006300.1       DITTSLSAYKEDKQKFEKFKVDMIQTKNELVGSLQQNDKKLSEANNMYEEFLAHIDTTTNREKQMYNQANALKENLANVQ
PsimAD002_000007200.1      DITTSLSAYKEDKQKFEKFKVDMIQTKNELVGSLQQNDKKLSEANNMYEEFLAHIDTTTNREKQMYNQANALKENLANVQ
PsimAD005_000448800.1      DITTSLSAYKEDKQKFEKFKVDMIQTKNELVGSLQQNDKKLSEANNMYEEFLAHIDTTTNREKQMYNQANALKENLANVQ
PsimAF28_000006200.1       DITTSLSAYKEDKQKFEKFKVDMIQTKNELVGSLQQNDKKLSEANNMYEEFLAHIDTTTNREKQMYNQANALKENLANVQ
PsimAF33_000006700.1       DITTSLSAYKEDKQKFEKFKVDMIQTKNELVGSLQQNDKKLSEANNMYEEFLAHIDTTTNREKQMYNQANALKENLANVQ
PsimAF34_000005900.1       DITTSLSAYKEDKQKFEKFKVDMIQTKNELVGSLQQNDKKLSEANNMYEEFLAHIDTTTNREKQMYNQANALKENLANVQ
PvivaxAM01_000076800.1     DITTSLSAYKEDKQKFEKFKVDMIQTKNELLGSLQQNDKKLSEANNMYEEFLAHIDTTTNREKQMYNQANALKENLANVQ
PvivaxAM02_000474900.1     DITTSLSAYKEDKQKFEKFKVDMIQTKNELLGSLQQNDKKLSEANNMYEEFLAHIDTTTNREKQMYNQANALKENLANVQ
PVL_140006600-t42_1-p1     DITTSLSAYKEDKQKFEKFKVDMIQTKNELLGSLQQNDKKLSEANNMYEEFLAHIDTTTNRENQMYNQANALKENLANVQ

PVX_121920.1-p1            SKISSYQGKIEKLKAHTRKNYEKIKTLLQTFGNENINAKLAEYEKEFNENKRVVNTIINQIQESNKNIENIKILNNSIDR
PVP01_1402400.1-p1         SKISSYQGKIEKLKAHTRKNYEKIKTLLQTFGNENINAKLAEYEKEFNENKRVVNTIINQIQESNKNIENIKILNNSIDR
Psim2302_000007000.1       SKISSYQGKIEKLKAHTRKNYEKIKTLLQTFGNENINAKLAEYEKEFNENKRVVNTIINQIQESNKNIENIKILNNSIDR
Psim3636_000006300.1       SKISSYQGKIEKLKAHTRKNYEKIKTLLQTFGNENINAKLAEYEKEFNENKRVVNTIINQIQESNKNIENIKILNNSIDR
PsimAD002_000007200.1      SKISSYQGKIEKLKAHTRKNYEKIKTLLQTFGNENINAKLAEYEKEFNENKRVVNTIINQIQESNKNIENIKILNNSIDR
PsimAD005_000448800.1      SKISSYQGKIEKLKAHTRKNYEKIKTLLQTFGNENINAKLAEYEKEFNENKRVVNTIINQIQESNKNIENIKILNNSIDR
PsimAF28_000006200.1       SKISSYQGKIEKLKAHTRKNYEKIKTLLQTFGNENINAKLAEYEKEFNENKRVVNTIINQIQESNKNIENIKILNNSIDR
PsimAF33_000006700.1       SKISSYQGKIEKLKAHTRKNYEKIKTLLQTFGNENINAKLAEYEKEFNENKRVVNTIINQIQESNKNIENIKILNNSIDR
PsimAF34_000005900.1       SKISSYQGKIEKLKAHTRKNYEKIKTLLQTFGNENINAKLAEYEKEFNENKRVVNTIINQIQESNKNIENIKILNNSIDR
PvivaxAM01_000076800.1     SKISSYQGKIEKLKAHTRKNYEKIKTLLQTFGNENINAKLAEYEKEFNENKEVVNTIINQIQESNKNIENIKILNNSIDR
PvivaxAM02_000474900.1     SKISSYQGKIEKLKAHTRKNYEKIKTLLQTFGNENINAKLAEYEKEFNENKEVVNTIINQIQESNKNIENIKILNNSIDR
PVL_140006600-t42_1-p1     SKISSYQGKIEKLKAHTRKNYEKIKTLLQTFGNENINAKLAEYEKEFNENKRVVNTIINQIQESNKNIENIKILNNSIDR
```

### Figure S25 (cont.)

```
PVX_121920.1-p1           SEVNKQLIEELATNKQQLKEKIDSHIKQLNSYNIIAKDDKLIFEKRLNEEKANINTQLSDTSIDNLKAQIQETLDYYKTA
PVP01_1402400.1-p1        SEVNKQLIEELATNKQQLKEKIDSHIKQLNSYNIIAKDDKLIFEKTLNEEKANINTQLSDTSIDNLKAQIQETLDYYKTA
Psim2302_000007000.1      SEVNKQLIEELATNKQQLKEKIDSHIKQLNSYNIIAKDDKLIFEKTLNEEKANINTQLSDTSIDNLKAQIQETLDYYKTA
Psim3636_000006300.1      SEVNKQLIEELATNKQQLKEKIDSHIKQLNSYNIIAKDDKLIFEKTLNEEKANINTQLSDTSIDNLKAQIQETLDYYKTA
PsimAD002_000007200.1     SEVNKQLIEELATNKQQLKEKIDSHIKQLNSYNIIAKDDKLIFEKTLNEEKANINTQLSDTSIDNLKAQIQETLDYYKTA
PsimAD005_000448800.1     SEVNKQLIEELATNKQQLKEKIDSHIKQLNSYNIIAKDDKLIFEKTLNEEKANINTQLSDTSIDNLKAQIQETLDYYKTA
PsimAF28_000006200.1      SEVNKQLIEELATNKQQLKEKIDSHIKQLNSYNIIAKDDKLIFEKTLNEEKANINTQLSDTSIDNLKAQIQETLDYYKTA
PsimAF33_000006700.1      SEVNKQLIEELATNKQQLKEKIDSHIKQLNSYNIIAKDDKLIFEKTLNEEKANINTQLSDTSIDNLKAQIQETLDYYKTA
PsimAF34_000005900.1      SEVNKQLIEELATNKQQLKEKIDSHIKQLNSYNIIAKDDKLIFEKTLNEEKANINTQLSDTSIDNLKAQIQETLDYYKTA
PvivaxAM01_000076800.1    SEVNKQLIEELATNKQQLKEKIDSHIKQLNSYNIIAKDDKLIFEKTLNEEKANINTQLSDTSIDNLKAQIQETLDYYKTA
PvivaxAM02_000474900.1    SEVNKQLIEELATNKQQLKEKIDSHIKQLNSYNIIAKDDKLIFEKTLNEEKANINTQLSDTSIDNLKAQIQETLDYYKTA
PVL_140006600-t42_1-p1    SEVNKQLIEELATNKQQLKEKIDNHIKQLNSYNIIAKDDKLIFEKTLDEEKANINTQLSDTSIDNLKAQIQEMLDYCKTA

PVX_121920.1-p1           KANTNDKDGTHLEKLDKKKMDWKDSKVKIDKLSTSYQVLDKKINDLIKKQHDEIVALIDKLITEKGNEIREKTDEKIKHL
PVP01_1402400.1-p1        KANTNDKDGTHLEKLDKKKMDWKDSKVKIDILSTSYQVLDKKINDLIKKQHDEIVALIDKLITEKGNEIREKTDEKIKHL
Psim2302_000007000.1      KANTNDKDGTHLEKLDKKKMDWKDSKVKIDKLSTSYQVLDKKINDLIKKQHDEIVALIDKLITEKGNEIREKTDEKIKHL
Psim3636_000006300.1      KANTNDKDGTHLEKLDKKKMDWKDSKVKIDKLSTSYQVLDKKINDLIKKQHDEIVALIDKLITEKGNEIREKTDEKIKHL
PsimAD002_000007200.1     KANTNDKDGTHLEKLDKKKMDWKDSKVKIDKLSTSYQVLDKKINDLIKKQHDEIVALIDKLITEKGNEIREKTDEKIKHL
PsimAD005_000448800.1     KANTNDKDGTHLEKLDKKKMDWKDSKVKIDKLSTSYQVLDKKINDLIKKQHDEIVALIDKLITEKGNEIREKTDEKIKHL
PsimAF28_000006200.1      KANTNDKDGTHLEKLDKKKMDWKDSKVKIDKLSTSYQVLDKKINDLIKKQHDEIVALIDKLITEKGNEIREKTDEKIKHL
PsimAF33_000006700.1      KANTNDKDGTHLEKLDKKKMDWKDSKVKIDKLSTSYQVLDKKINDLIKKQHDEIVALIDKLITEKGNEIREKTDEKIKHL
PsimAF34_000005900.1      KANTNDKDGTHLEKLDKKKMDWKDSKVKIDKLSTSYQVLDKKINDLIKKQHDEIVALIDKLITEKGNEIREKTDEKIKHL
PvivaxAM01_000076800.1    KANTNDKDGTHLEKLDKKKMDWKDSKVKIDKLSTSYQVLDKKINDLIKKQHDEIVALIDKLITEKGNEIREKTDEKIKHL
PvivaxAM02_000474900.1    KANTNDKDGTHLEKLDKKKMDWKDSKVKIDKLSTSYQVLDKKINDLIKKQHDEIVALIDKLITEKGNEIREKTDEKIKHL
PVL_140006600-t42_1-p1    KAKTNDKDGTHLEKLNKQKMDWKDSKVKIDKLSTSYQVLDKKINDLIKKQHDEIVALIDKLITEKGNEIREKTDEKIKHL

PVX_121920.1-p1           SEIKTKLSSFTVSDNVKNTQNVAIKGEISGFEKKMEALLRRIDNNNAKLTELRGKADQHVTTLRGEKNKTAEFNAKKDSL
PVP01_1402400.1-p1        SEIKTKLSSFTVSDNVKNTQNVAIKGEISGFEKKMEALLRRIDNNNAKLTELRGKADQHVTTLRGEKNKTAEFNAKKDSL
Psim2302_000007000.1      SEIKTKLSSFTVSDNVKNTQNVAIKGEISGFEKKMEALLRRIDNNNAKLTELRGKADQHVTTLRGEKNKTAEFNAKKDSL
Psim3636_000006300.1      SEIKTKLSSFTVSDNVKNTQNVAIKGEISGFEKKMEALLRRIDNNNAKLTELRGKADQHVTTLRGEKNKTAEFNAKKDSL
PsimAD002_000007200.1     SEIKTKLSSFTVSDNVKNTQNVAIKGEISGFEKKMEALLRRIDNNNAKLTELRGKADQHVTTLRGEKNKTAEFNAKKDSL
PsimAD005_000448800.1     SEIKTKLSSFTVSDNVKNTQNVAIKGEISGFEKKMEALLRRIDNNNAKLTELRGKADQHVTTLRGEKNKTAEFNAKKDSL
PsimAF28_000006200.1      SEIKTKLSSFTVSDNVKNTQNVAIKGEISGFEKKMEALLRRIDNNNAKLTELRGKADQHVTTLRGEKNKTAEFNAKKDSL
PsimAF33_000006700.1      SEIKTKLSSFTVSDNVKNTQNVAIKGEISGFEKKMEALLRRIDNNNAKLTELRGKADQHVTTLRGEKNKTAEFNAKKDSL
PsimAF34_000005900.1      SEIKTKLSSFTVSDNVKNTQNVAIKGEISGFEKKMEALLRRIDNNNAKLTELRGKADQHVTTLRGEKNKTAEFNAKKDSL
PvivaxAM01_000076800.1    SEIKTKLSSFTVSDNVKNTQNVAIKGEISGFEKKMEALLRRIDNNNAKLTELRGKADQHVTTLRGEKNKTAEFNAKKDSL
PvivaxAM02_000474900.1    SEIKTKLSSFTVSDNVKNTQNVAIKGEISGFEKKMEALLRRIDNNNAKLTELRGKADQHVTTLRGEKNKTAEFNAKKDSL
PVL_140006600-t42_1-p1    SEIKTKLSSFTVSDNVKNTQNVAIKGEVNGFEKKIEALLKRVDNNNAKLTELRGKADQHVTTLRGEKNKTAEFNAKKDSL

PVX_121920.1-p1           EKIHQQMENTHKELESMENEKIPINDINQMEIEYARILIHHMVQQIGEKKERSKIILEEIKTTIASIDQAKKNIPPEQQD
PVP01_1402400.1-p1        EKIHQQMENTHKELESMENEKIPINDINQMEIEYARILIHHMVQQIGEKKERSKIILEEIKTTIASIDQAKKNIPPEQQD
Psim2302_000007000.1      EKIHQQMENTHKELESMENEKIPINDINQMEIEYARILIHHMVQQIGEKKERSKIILEEIKTTIASIDQAKKNIPPEQQD
Psim3636_000006300.1      EKIHQQMENTHKELESMENEKIPINDINQMEIEYARILIHHMVQQIGEKKERSKIILEEIKTTIASIDQAKKNIPPEQQD
PsimAD002_000007200.1     EKIHQQMENTHKELESMENEKIPINDINQMEIEYARILIHHMVQQIGEKKERSKIILEEIKTTIASIDQAKKNIPPEQQD
PsimAD005_000448800.1     EKIHQQMENTHKELESMENEKIPINDINQMEIEYARILIHHMVQQIGEKKERSKIILEEIKTTIASIDQAKKNIPPEQQD
PsimAF28_000006200.1      EKIHQQMENTHKELESMENEKIPINDINQMEIEYARILIHHMVQQIGEKKERSKIILEEIKTTIASIDQAKKNIPPEQQD
PsimAF33_000006700.1      EKIHQQMENTHKELESMENEKIPINDINQMEIEYARILIHHMVQQIGEKKERSKIILEEIKTTIASIDQAKKNIPPEQQD
PsimAF34_000005900.1      EKIHQQMENTHKELESMENEKIPINDINQMEIEYARILIHHMVQQIGEKKERSKIILEEIKTTIASIDQAKKNIPPEQQD
PvivaxAM01_000076800.1    EKIHQQMENTHKELESMENEKIPINDINQMEIEYARILIHHMVQQIGEKKERSKIILEEIKTTIASIDQAKKNIPPEQQD
PvivaxAM02_000474900.1    EKIHQQMENTHKELESMENEKIPINDINQMEIEYARILIHHMVQQIGEKKERSKIILEEIKTTIASIDQAKKNIPPEQQD
PVL_140006600-t42_1-p1    EKIHQQMENTHKELESMENEKIPINDINQMEIEYGRILIHHMVQQIGEKKERSKSILEEITSTIESIEQAKQNIPPEQQD

PVX_121920.1-p1           ILREFEYSGYRDKATASSTEIEQIDKSAKAEEGKANSSTNLSDIDVIKKQVLSYFRKATTESNTMENALREIKNVNTFLT
PVP01_1402400.1-p1        ILREFEYSGYRDKATASSTEIEQIDKSAKAEEGKANSSTNLSDIDVIKKQVLSYFRKATTESNTMENALREIKNVNTFLT
Psim2302_000007000.1      ILREFEYSGYRDKATASSTEIEQIDKSAKAEEGKANSSTNLSDIDVIKKQVLSYFRKATTESNTMENALREIKNVNTFLT
Psim3636_000006300.1      ILREFEYSGYRDKATASSTEIEQIDKSAKAEEGKANSSTNLSDIDVIKKQVLSYFRKATTESNTMENALREIKNVNTFLT
PsimAD002_000007200.1     ILREFEYSGYRDKATASSTEIEQIDKSAKAEEGKANSSTNLSDIDVIKKQVLSYFRKATTESNTMENALREIKNVNTFLT
PsimAD005_000448800.1     ILREFEYSGYRDKATASSTEIEQIDKSAKAEEGKANSSTNLSDIDVIKKQVLSYFRKATTESNTMENALREIKNVNTFLT
PsimAF28_000006200.1      ILREFEYSGYRDKATASSTEIEQIDKSAKAEEGKANSSTNLSDIDVIKKQVLSYFRKATTESNTMENALREIKNVNTFLT
PsimAF33_000006700.1      ILREFEYSGYRDKATASSTEIEQIDKSAKAEEGKANSSTNLSDIDVIKKQVLSYFRKATTESNTMENALREIKNVNTFLT
PsimAF34_000005900.1      ILREFEYSGYRDKATASSTEIEQIDKSAKAEEGKANSSTNLSDIDVIKKQVLSYFRKATTESNTMENALREIKNVNTFLT
PvivaxAM01_000076800.1    ILREFEYSGYRDKATASSTEIEQIDKSAKAEEGKANSSTNLSDIDVIKKQVLSYFRKATTESNTMENALREIKNVNTFLT
PvivaxAM02_000474900.1    ILREFEYSGYRDKATASSTEIEQIDKSAKAEEGKANSSTNLSDIDVIKKQVLSYFRKATTESNTMENALREIKNVNTFLT
PVL_140006600-t42_1-p1    ILHEFEYSGYRDKATASSTEIEQIDKSAKTEEGKANSSTNLSDIDVIKKEVLSYLRKATTESNTMEDALSEIKNVNTFLT

PVX_121920.1-p1           SNISEEILANILKSAARTQELNQQAVGEFNKTDRLIKEVEAKLSQANEHKSAISGSVEYKQIEQKINLIKQIQKEITAGK
PVP01_1402400.1-p1        SNISEEILANILKSAARTQELNQQAVGEFNKTDRLIKEVEAKLSQANEHKSAISGSVEYKQIEQKINLIKQIQKEITAGK
Psim2302_000007000.1      SKNSEEILA-----------------------------------------------------------------------
Psim3636_000006300.1      SKNSEEILA-----------------------------------------------------------------------
PsimAD002_000007200.1     SKNSEEILA-----------------------------------------------------------------------
PsimAD005_000448800.1     SKNSEEILA-----------------------------------------------------------------------
PsimAF28_000006200.1      SKNSEEILA-----------------------------------------------------------------------
PsimAF33_000006700.1      SKNSEEILA-----------------------------------------------------------------------
PsimAF34_000005900.1      SKNSEEILA-----------------------------------------------------------------------
PvivaxAM01_000076800.1    SNISEEILANILKSAARTQELNQQAVGEFNKTDRLIKEVEAKLSQANEHKSAISGSVEYKQIEQKINLIKQIQKEITAGK
PvivaxAM02_000474900.1    SNISEEILANILKSAARTQELNQQAVGEFNKTDRLIKEVEAKLSQANEHKSAISGSVEYKQIEQKINLIKQIQKEITAGK
PVL_140006600-t42_1-p1    SNISEEILANILKSAARTQELNQQAVGEFNKADRLIKEVEAKLSQATEHKSAIPGSAEYKQIEQKINLIKQIQKEITAGK
```

Figure S25 (cont.)

```
PVX_121920.1-p1           EEINNCLSNTKEYKEKCESEVNSVNRGKAKVDFLQKREALEK-KMSQENLGKITDSIDQCKKDLADITSLEVKVKANYDS
PVP01_1402400.1-p1        EEINNCLSNTKEYKEKCESEVNSVNRGKAKVDFLQKREALEK-KMSQENLGKITDSIDQCKKDLADITSLEVKVKANYDS
Psim2302_000007000.1      --------------------------------------------------------------------------------
Psim3636_000006300.1      --------------------------------------------------------------------------------
PsimAD002_000007200.1     --------------------------------------------------------------------------------
PsimAD005_000448800.1     --------------------------------------------------------------------------------
PsimAF28_000006200.1      --------------------------------------------------------------------------------
PsimAF33_000006700.1      --------------------------------------------------------------------------------
PsimAF34_000005900.1      --------------------------------------------------------------------------------
PvivaxAM01_000076800.1    EEINNCLSNTKEYKEKCESEVNSVNRGKAKVDFLQKREALEK-KMSQENLGKITDSIDQCKKDLADITSLEVKVKANYDS
PvivaxAM02_000474900.1    EEINNCLSNTKEYKEKCESEVNSVNRGKAKVDFLQKREALEK-KMSQENLGKITDSIDQCKKDLADITSLEVKVKANYDS
PVL_140006600-t42_1-p1    EEINTCLSNTKEYKEKCESEVNSVNRGKAKVDFLQKRKELEENRMSQENLSKITDSIDQCKKDLAEIASSELKVKANYDS

PVX_121920.1-p1           IIKYEESINTILNYSSILEYKTKLEIRKKEKTDLMTYINTENSAIQEKLLNLQKKLNQLNENTDYTKVGNDLNNAKSTKA
PVP01_1402400.1-p1        IIKYEESINTILNYSSILEYKTKLEIRKKEKTDLMTYINTENSAIQEKLLNLQKKLNQLNENTDYTKVGNDLNNAKSTKA
Psim2302_000007000.1      --------------------------------------------------------------------------------
Psim3636_000006300.1      --------------------------------------------------------------------------------
PsimAD002_000007200.1     --------------------------------------------------------------------------------
PsimAD005_000448800.1     --------------------------------------------------------------------------------
PsimAF28_000006200.1      --------------------------------------------------------------------------------
PsimAF33_000006700.1      --------------------------------------------------------------------------------
PsimAF34_000005900.1      --------------------------------------------------------------------------------
PvivaxAM01_000076800.1    IIKYEESINTILNYSSILEYKTKLEIRKKEKTDLMTYINTENSAIQEKLLNLQKKLNQLNENTDYTKVGNDLNNAKSTKA
PvivaxAM02_000474900.1    IIKYEESINTILNYSSILEYKTKLEIRKKEKTDLMTYINTENSAIQEKLLNLQKKLNQLNENTDYTKVGNDLNNAKSTKA
PVL_140006600-t42_1-p1    IIKYEESINTILNYSSILEYQTKLEIRKKEKTDLMNYINTENSAIQEKLLNLQKKLNQLNENTDYTKVGNDLNNAKSTKA

PVX_121920.1-p1           NVTIQYNLGRVKHQLENLSVIKQELEKVLSAATDLERDISKIADVTESNNLESLNGKEADYTKRIRSFNKLKQLVQEKAA
PVP01_1402400.1-p1        NVIIQYNLGRVKHQLENVSVIKQELEKVLSAATDLERDISKIADVTESNNLESLNGKEADYTKRIRSFNKLKQLVQEKAA
Psim2302_000007000.1      --------------------------------------------------------------------------------
Psim3636_000006300.1      --------------------------------------------------------------------------------
PsimAD002_000007200.1     --------------------------------------------------------------------------------
PsimAD005_000448800.1     --------------------------------------------------------------------------------
PsimAF28_000006200.1      --------------------------------------------------------------------------------
PsimAF33_000006700.1      --------------------------------------------------------------------------------
PsimAF34_000005900.1      --------------------------------------------------------------------------------
PvivaxAM01_000076800.1    NVTIQYNLGRVKHQLENVSVIKQELEKVLSAATDLERDISKIADVTESNNLESLNGKEADYTKRIRSFNKLKQLVQEKAA
PvivaxAM02_000474900.1    NVTIQYNLGRVKHQLENLSVIKQELEKVLSAATDLERDTSKIADVTESNNLESLNGKEADYTKRIRSFNKLKQLVQEKAA
PVL_140006600-t42_1-p1    NVTIQYNLGRVKHQLENVSVIKQELEKVLSAATDLEREVSKISDVTESNNLESLNGKEADYTKHIKSFNKLKLLVQEKAA

PVX_121920.1-p1           KVEEISSDTDNIEKELTEHKIIFEVGLTERLIEIVKNRKSYVDTTKELLNSSLNNFASLFNGLDLNGYNPKANLEMYTQK
PVP01_1402400.1-p1        KVEEISSDTDNIEKELTEHKIIFEVGLTERLIEIVKNRKSYVDTTKELLNSSLNNFASLFNGLDLKGYNPKANLEMYTQK
Psim2302_000007000.1      --------------------------------------------------------------------------------
Psim3636_000006300.1      --------------------------------------------------------------------------------
PsimAD002_000007200.1     --------------------------------------------------------------------------------
PsimAD005_000448800.1     --------------------------------------------------------------------------------
PsimAF28_000006200.1      --------------------------------------------------------------------------------
PsimAF33_000006700.1      --------------------------------------------------------------------------------
PsimAF34_000005900.1      --------------------------------------------------------------------------------
PvivaxAM01_000076800.1    KVEEISSDTDNIEKELTEHKIIFEVGLTERLIEIVKNRKSYVDTTKELLNSSLNNFASLFNGLDLNGYNPKANLEMYTQK
PvivaxAM02_000474900.1    KVEEISSDTDNIEKELTEHKIIFEVGLTERLIEIVKNRKSYVDTTKELLNSSLNNFASLFNGLDLNGYNPKANLEMYTQK
PVL_140006600-t42_1-p1    KVEEISSDTDNIEKELTEHKIIFEVGLTERLIEIIKNRKSYVDTTKELLNSSLNNFASLFNGLDLKGYNPKANLEMYTQK

PVX_121920.1-p1           LNTIHNEFMASHKIFDEKSKKVLDKDVNFTEAKTLREEAQKEDVTLKNKEEEAKSYLSDIKKKESFEFILHMKEKLNQIS
PVP01_1402400.1-p1        LNTIHNEFMASHKIFDEKSKKVLDKDVNFTEAKTLREEAQKEDVTLKNKEEEAKSYLSDIKKKESFEFILHMKEKLNQIS
Psim2302_000007000.1      --------------------------------------------------------------------------------
Psim3636_000006300.1      --------------------------------------------------------------------------------
PsimAD002_000007200.1     --------------------------------------------------------------------------------
PsimAD005_000448800.1     --------------------------------------------------------------------------------
PsimAF28_000006200.1      --------------------------------------------------------------------------------
PsimAF33_000006700.1      --------------------------------------------------------------------------------
PsimAF34_000005900.1      --------------------------------------------------------------------------------
PvivaxAM01_000076800.1    LNTIHNEFMASHKIFDEKSKKVLDKDVNFTEAKTLREEAQKEDVTLKNKEEEAKSYLSDIKKKESFEFILHMKEKLNQIS
PvivaxAM02_000474900.1    LNTIHNEFMASHKIFDEKSKKVLDKDVNFTEAKTLREEAQKEDVTLKNKEEEAKSYLSDIKKKESFEFILHMKEKLNQIS
PVL_140006600-t42_1-p1    LNTIHNEFMASHKIFDEKSKKVLVKDVNFTEAKTLREEAQKEDVTLKNKEEEAKSYLSDIKKKESFEFILHMKEKLTQIS

PVX_121920.1-p1           KMCEQQYEQADKGYSEVKTSIDRIANLNDENSIADVLKEANDKNEQVQNLTHYTYKNEAQNVLRHMAKSANFIGINLVTG
PVP01_1402400.1-p1        KMCEQQYEQADKGYSEVKTSIDRIANLNDENSIADVLKEANDKNEQVQNLTHYTYKNEAQNVLRHMAKSANFIGINLVTG
Psim2302_000007000.1      --------------------------------------------------------------------------------
Psim3636_000006300.1      --------------------------------------------------------------------------------
PsimAD002_000007200.1     --------------------------------------------------------------------------------
PsimAD005_000448800.1     --------------------------------------------------------------------------------
PsimAF28_000006200.1      --------------------------------------------------------------------------------
PsimAF33_000006700.1      --------------------------------------------------------------------------------
PsimAF34_000005900.1      --------------------------------------------------------------------------------
PvivaxAM01_000076800.1    KMCEQQYEQADKGYSEVKTSIDRIANLNDENSIADVLKEANDKNEQVQNLTHYTYKNEAQNVLRHMAKSANFIGINLVTG
PvivaxAM02_000474900.1    KMCEQQYEQADKGYSEVKTSIDRIANLNDENSIADVLKEANDKNEQVQNLTHYTYKNEAQNVLRHMAKSANFIGINLVTG
PVL_140006600-t42_1-p1    KMCEQQYEQADKGYSEVKTSIDRIANLNDENSVADVLKEANDQNEQVQNLTHYTYKNEAQNVLRHMAKSANFIGINLVTG
```

### Figure S25 (cont.)

```
PVX_121920.1-p1           IQPTELSSQASESTTPELKFESEGEMKLEILTLSGNTTKLDYYKNMKDAYQSVLSIFKYSHGIDEKQRESQKITESANGS
PVP01_1402400.1-p1        IQPTELSSQASESTTPELKFESEGEMKLEILTLSGNTTKLDYYKNMKDAYQSVLSIFKYSHGIDEKQRESQKITESANGS
Psim2302_000007000.1      --------------------------------------------------------------------------------
Psim3636_000006300.1      --------------------------------------------------------------------------------
PsimAD002_000007200.1     --------------------------------------------------------------------------------
PsimAD005_000448800.1     --------------------------------------------------------------------------------
PsimAF28_000006200.1      --------------------------------------------------------------------------------
PsimAF33_000006700.1      --------------------------------------------------------------------------------
PsimAF34_000005900.1      --------------------------------------------------------------------------------
PvivaxAM01_000076800.1    IQPTELSSQASESTTPELKFESEGEMKLEILTLSGNTTKLDYYKNMKDAYQSVLSIFKYSHGIDEKQRESQKITESANGS
PvivaxAM02_000474900.1    IQPTELSSQASESTTPELKFESEGEMKLEILTLSGNTTKLDYYKNMKDAYQSVLSIFKYSHGIDEKQRESQKITESANGS
PVL_140006600-t42_1-p1    IQPTELSSQASESTTPELKFESEREMKLEILTLSGNTTKLDYYKNMKDAYQSVLSILKYSHGIDEKQRESQKITESANGS

PVX_121920.1-p1           YLNNKEINEFKGRLNNVKSKQTAISNKIDNATTLLHNLNKIKTDDKNYDTILEKDASEELKRRRDSFNQEMKNTVDGLKL
PVP01_1402400.1-p1        YLNNKEINEFKGRLNNVKSKQTAISNKIDNATTLLHNLNKIKTDDKNYDTILEKDASEELKRRRDSFNQEIKNTVDGLKL
Psim2302_000007000.1      --------------------------------------------------------------------------------
Psim3636_000006300.1      --------------------------------------------------------------------------------
PsimAD002_000007200.1     --------------------------------------------------------------------------------
PsimAD005_000448800.1     --------------------------------------------------------------------------------
PsimAF28_000006200.1      --------------------------------------------------------------------------------
PsimAF33_000006700.1      --------------------------------------------------------------------------------
PsimAF34_000005900.1      --------------------------------------------------------------------------------
PvivaxAM01_000076800.1    YLNNKEINEFKGRLNNVKSKQTAISNKIDNATTLLHNLNKIKTDDKNYDTILEKDASEELKRRRDSFNQEMKNTVDGLKL
PvivaxAM02_000474900.1    YLNNKEINEFKGRLNNVKSKQTAISNKIDNATTLLHNLNKIKTDDKNYDTILEKDASEELKRRRDSFNQEMKNTVDGLKL
PVL_140006600-t42_1-p1    YLNNKAINEFKGRLNNVKSKQTAISNKIDNATTLLHNLNKIKTDDKNYDTILEKDASEELKRRRDSFNQEMKNTVDGLKL

PVX_121920.1-p1           KEIQEKFNEQVKLLQNLETKVSTLNVHEGNATETVKKENTAVDAIQAAMEGIEKDVEYINYSYDELLKKGQKIENQRYTS
PVP01_1402400.1-p1        KEIQEKFNEQVKLLQNLETKVSTLNVHEGNATETVKKENTAVDAIQAAMEGIEKDVEYINYSYDELLKKGQKIENQRYTS
Psim2302_000007000.1      --------------------------------------------------------------------------------
Psim3636_000006300.1      --------------------------------------------------------------------------------
PsimAD002_000007200.1     --------------------------------------------------------------------------------
PsimAD005_000448800.1     --------------------------------------------------------------------------------
PsimAF28_000006200.1      --------------------------------------------------------------------------------
PsimAF33_000006700.1      --------------------------------------------------------------------------------
PsimAF34_000005900.1      --------------------------------------------------------------------------------
PvivaxAM01_000076800.1    KEIQEKFNEQVKLLQNLETKVSTLNVHEGNATETVKKENTAVDAIQAAMEGIEKDVEYINYSYDELLKKGQKIENQRYTS
PvivaxAM02_000474900.1    KEIQEKFNEQVKLLQNLETKVSTLNVHEGNATETVKKENTAVDAIQAAMEGIEKDVEYINYSYDELLKKGQKIENQRYTS
PVL_140006600-t42_1-p1    KEIQEKFNEQVKLLQNLETKVSTLNVHEGNVTETVKKENTAVDAIQAAMEGIEKDVEYINYSYDELLKKGQKIENQRYTS

PVX_121920.1-p1           IRENLTNKIANDSSAINKIKKKAQQYLAYIKNNYNSIYNDTGTLNEYFDTKRLSNHDLTNVQEATRLHIEMSAAVEASEE
PVP01_1402400.1-p1        IRENLTNKIANDSSAINKIKKKAQEYLAYIKNNYNSIYNDTGTLNEYFDTKRLSNHDLTNVQEATRLHIEMSAAVEASEE
Psim2302_000007000.1      --------------------------------------------------------------------------------
Psim3636_000006300.1      --------------------------------------------------------------------------------
PsimAD002_000007200.1     --------------------------------------------------------------------------------
PsimAD005_000448800.1     --------------------------------------------------------------------------------
PsimAF28_000006200.1      --------------------------------------------------------------------------------
PsimAF33_000006700.1      --------------------------------------------------------------------------------
PsimAF34_000005900.1      --------------------------------------------------------------------------------
PvivaxAM01_000076800.1    IRENLTNKIANDSSAINKIKEKAQQYLAYIKNNYNSIYNDTGTLNEYFDTKRLSNHDLTNVQEATRLHIEMSAAVEASEE
PvivaxAM02_000474900.1    IRENLTNKIANDSSAINKIKEKAQQYLAYIKNNYNSIYNDTGTLNEYFDTKRLSNHDLTNVQEATRLHIEMSAAVEASEE
PVL_140006600-t42_1-p1    IRENLTNKIANDSSAINKIKKKAQQYLAYIKNNYNSIYNDIGTLNEYFDIKRLSNHDLTNVQEATRLHIEMSAAVEASEE

PVX_121920.1-p1           IIADMKNEFITNTEADISALQNSADRLKSLYSNLKRKQISINQIYKKINLIKLQEIKTSANKYMDIAKLFNNVLEAQHKE
PVP01_1402400.1-p1        IIADMKNEFITNTEADISALQNSADRLMSLYSNLKRKQISINQIYKKINLIKLQEIKTSANKYMDIAKLFNNVLEAQHKE
Psim2302_000007000.1      --------------------------------------------------------------------------------
Psim3636_000006300.1      --------------------------------------------------------------------------------
PsimAD002_000007200.1     --------------------------------------------------------------------------------
PsimAD005_000448800.1     --------------------------------------------------------------------------------
PsimAF28_000006200.1      --------------------------------------------------------------------------------
PsimAF33_000006700.1      --------------------------------------------------------------------------------
PsimAF34_000005900.1      --------------------------------------------------------------------------------
PvivaxAM01_000076800.1    IIADMKNEFITNTEADISALQNSADRLKSLYSNLKRKQISINQIYKKINLIKLQEIKTSANKYMDIAKLFNNVLEAQHKE
PvivaxAM02_000474900.1    IIADMKNEFITNTEADISALQNSADRLKSLYSNLKRKQISINQIYKKINLIKLQEIKTSANKYMDIAKLFNNVLEAQHKE
PVL_140006600-t42_1-p1    IIADMKNEFITNTEADISALQNSADRLKSLYSNLKRKQISINQIYKKINLIKLQEIKTSANKYMDIAKLFNNVLEAQHKK

PVX_121920.1-p1           LAQDRSKILQAK--------------------------------------------------------------------
PVP01_1402400.1-p1        LAQDISKILQAK--------------------------------------------------------------------
Psim2302_000007000.1      --------------------------------------------------------------------------------
Psim3636_000006300.1      --------------------------------------------------------------------------------
PsimAD002_000007200.1     --------------------------------------------------------------------------------
PsimAD005_000448800.1     --------------------------------------------------------------------------------
PsimAF28_000006200.1      --------------------------------------------------------------------------------
PsimAF33_000006700.1      --------------------------------------------------------------------------------
PsimAF34_000005900.1      --------------------------------------------------------------------------------
PvivaxAM01_000076800.1    LAQDRSKILQAK--------------------------------------------------------------------
PvivaxAM02_000474900.1    LAQDRSKILQAK--------------------------------------------------------------------
PVL_140006600-t42_1-p1    LAQDRSKILQVKEKLINTEKELANLDETITLQSLKKSNELCNSATKNIQDIRELEKENNKEDKKIKIYGEKISHLINRRK
```

### Figure S25 (cont.)

```
PVX_121920.1-p1            --------------------------------------------------------------------------------
PVP01_1402400.1-p1         --------------------------------------------------------------------------------
Psim2302_000007000.1       --------------------------------------------------------------------------------
Psim3636_000006300.1       --------------------------------------------------------------------------------
PsimAD002_000007200.1      --------------------------------------------------------------------------------
PsimAD005_000448800.1      --------------------------------------------------------------------------------
PsimAF28_000006200.1       --------------------------------------------------------------------------------
PsimAF33_000006700.1       --------------------------------------------------------------------------------
PsimAF34_000005900.1       --------------------------------------------------------------------------------
PvivaxAM01_000076800.1     --------------------------------------------------------------------------------
PvivaxAM02_000474900.1     --------------------------------------------------------------------------------
PVL_140006600-t42_1-p1     VLLNDVSEYDRTENFDHENEQAANDLQNDIATIKKVLVSSEEQYRKLLENVKKNESLYSNNDTKNFTLEISKKIENVKRK

PVX_121920.1-p1            --------------------------------------------------------------------------------
PVP01_1402400.1-p1         --------------------------------------------------------------------------------
Psim2302_000007000.1       --------------------------------------------------------------------------------
Psim3636_000006300.1       --------------------------------------------------------------------------------
PsimAD002_000007200.1      --------------------------------------------------------------------------------
PsimAD005_000448800.1      --------------------------------------------------------------------------------
PsimAF28_000006200.1       --------------------------------------------------------------------------------
PsimAF33_000006700.1       --------------------------------------------------------------------------------
PsimAF34_000005900.1       --------------------------------------------------------------------------------
PvivaxAM01_000076800.1     --------------------------------------------------------------------------------
PvivaxAM02_000474900.1     --------------------------------------------------------------------------------
PVL_140006600-t42_1-p1     ISINIPESEQLLQIENRFGDIKAIINGIKADNDVDEYVEEVYKNIQREKEKLTDMRNGEQVKEAIKNITHYNDETKNKLS

PVX_121920.1-p1            ----------------------------------------ENSINMQQLLESHIDKLRALITNIDKELIELTNGKIKESN
PVP01_1402400.1-p1         ----------------------------------------ENSINMQQLLESHIDKLRALITNIDKELIELTNGKIKASN
Psim2302_000007000.1       --------------------------------------------------------------------------------
Psim3636_000006300.1       --------------------------------------------------------------------------------
PsimAD002_000007200.1      --------------------------------------------------------------------------------
PsimAD005_000448800.1      --------------------------------------------------------------------------------
PsimAF28_000006200.1       --------------------------------------------------------------------------------
PsimAF33_000006700.1       --------------------------------------------------------------------------------
PsimAF34_000005900.1       --------------------------------------------------------------------------------
PvivaxAM01_000076800.1     ----------------------------------------ENSINMQQLLESHIDKLRALITNIDKELIELTNDKIKASN
PvivaxAM02_000474900.1     ----------------------------------------ENSINMQQLLESHIDKLRALITNIDKELIELTNGKIKESN
PVL_140006600-t42_1-p1     RIYNAFEKVKMKKKDMEKIFASISEKSENNAIQNDVKNAIEHSINMQQLLESHIDKLRALITNIDKELIELKNGKIKASN

PVX_121920.1-p1            RRISQISPMGQGKLFSTPEGQAYNNLHNTGYNHYGSGNHSRGRNENGNVRFAAGIVVGFGVCSFFASALFKGKGENETYG
PVP01_1402400.1-p1         RRISQISPMGQGKLFSTPEGQAYNNLHNTGYNHNGSGNHSRGRNENGNVRFAAGIVVGFGVCSFFASALFKGKGENETYG
Psim2302_000007000.1       --ISQISPMEQGKLFSTPEGQAYNNLHNTGYNHYGSGNHSRGRNENGNVRFAAGIVVVFGVCSFFASALFKGKGENETYG
Psim3636_000006300.1       --ISQISPMEQGKLFSTPEGQAYNNLHNTGYNHYGSGNHSRGRNENGNVRFAAGIVVVFGVCSFFASALFKGKGENETYG
PsimAD002_000007200.1      --ISQISPMEQGKLFSTPEGQAYNNLHNTGYNHYGSGNHSRGRNENGNVRFAAGIVVVFGVCSFFASALFKGKGENETYG
PsimAD005_000448800.1      --ISQISPMEQGKLFSTPEGQAYNNLHNTGYNHYGSGNHSRGRNENGNVRFAAGIVVVFGVCSFFASALFKGKGENETYG
PsimAF28_000006200.1       --ISQISPMEQGKLFSTPEGQAYNNLHNTGYNHYGSGNHSRGRNENGNVRFAAGIVVVFGVCSFFASALFKGKGENETYG
PsimAF33_000006700.1       --ISQISPMEQGKLFSTPEGQAYNNLHNTGYNHYGSGNHSRGRNENGNVRFAAGIVVVFGVCSFFASALFKGKGENETYG
PsimAF34_000005900.1       --ISQISPMEQGKLFSTPEGQAYNNLHNTGYNHYGSGNHSRGRNENGNVRFAAGIVVVFGVCSFFASALFKGKGENETYG
PvivaxAM01_000076800.1     RRISQISPMGQGKLFSTPEGQAYNNLHNTGYNHYGSGNHSRGRNENGNVRFAAGIVVGFGVCSFFASALFKGKGENETYG
PvivaxAM02_000474900.1     RRISQISPMGQGKLFSTPEGQAYNNLHNTGYNHYGSGNHSRGRNENGNVRFAAGIVVGFGVCSFFASALFKGKGENETYG
PVL_140006600-t42_1-p1     RRISQISPMGQGKLFSTPEGQAYNNLHNTGYNHYGSGNHSRGRNENGNVRFAAGIVVGLGVCSFFASALFKGKKENETYG

PVX_121920.1-p1            RDLNSRDEEFEGKNNGNLQDKEEIIEVSFHESENVY
PVP01_1402400.1-p1         RDLNSRDEEFEGKNNGNLQDKEEIIEVSFHESENAY
Psim2302_000007000.1       RDLNSRDEEFEGKNNGNLQDKEEIIEVSFHESENVY
Psim3636_000006300.1       RDLNSRDDEFEGKNNGNLQDKEEIIEVSFHESENVY
PsimAD002_000007200.1      RDLNSRDDEFEGKNNGNLQDKEEIIEVSFHESENVY
PsimAD005_000448800.1      RDLNSRDDEFEGKNNGNLQDKEEIIEVSFHESENVY
PsimAF28_000006200.1       RDLNSRDDEFEGKNNGNLQDKEEIIEVSFHESENVY
PsimAF33_000006700.1       RDLNSRDEEFEGKNNGNLQDKEEIIEVSFHESENVY
PsimAF34_000005900.1       RDLNSRDEEFEGKNNGNLQDKEEIIEVSFHESENVY
PvivaxAM01_000076800.1     RDLNSRDEEFEGKNNGNLQDKEEIIEVSFHESENVY
PvivaxAM02_000474900.1     RDLNSRDEEFEGKNNGNLQDKEEIIEVSFHESENVY
PVL_140006600-t42_1-p1     RDLNSRDEEFEGKNNGNLQDKEEIIEVSFHESENVY
```

Figure S26

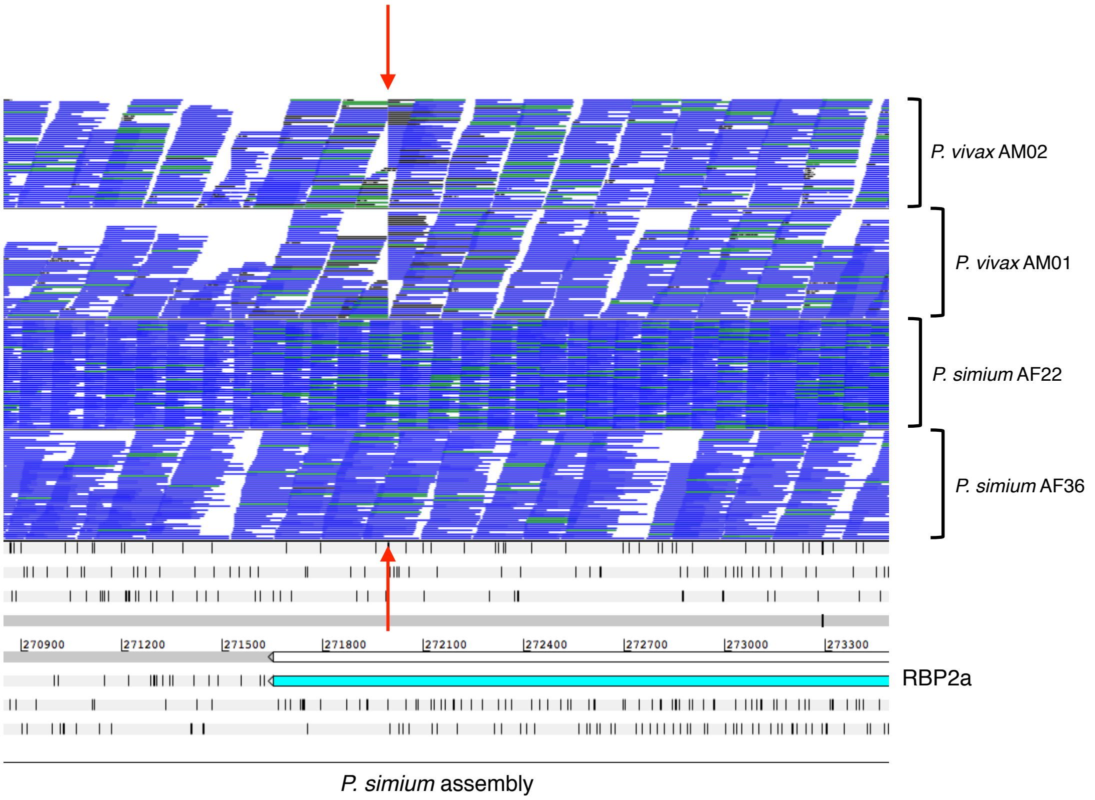

P. vivax AM02
P. vivax AM01
P. simium AF22
P. simium AF36
270900
271200
271500
271800
272100
272400
272700
273000
273300
RBP2a
P. simium assembly

Figure S27

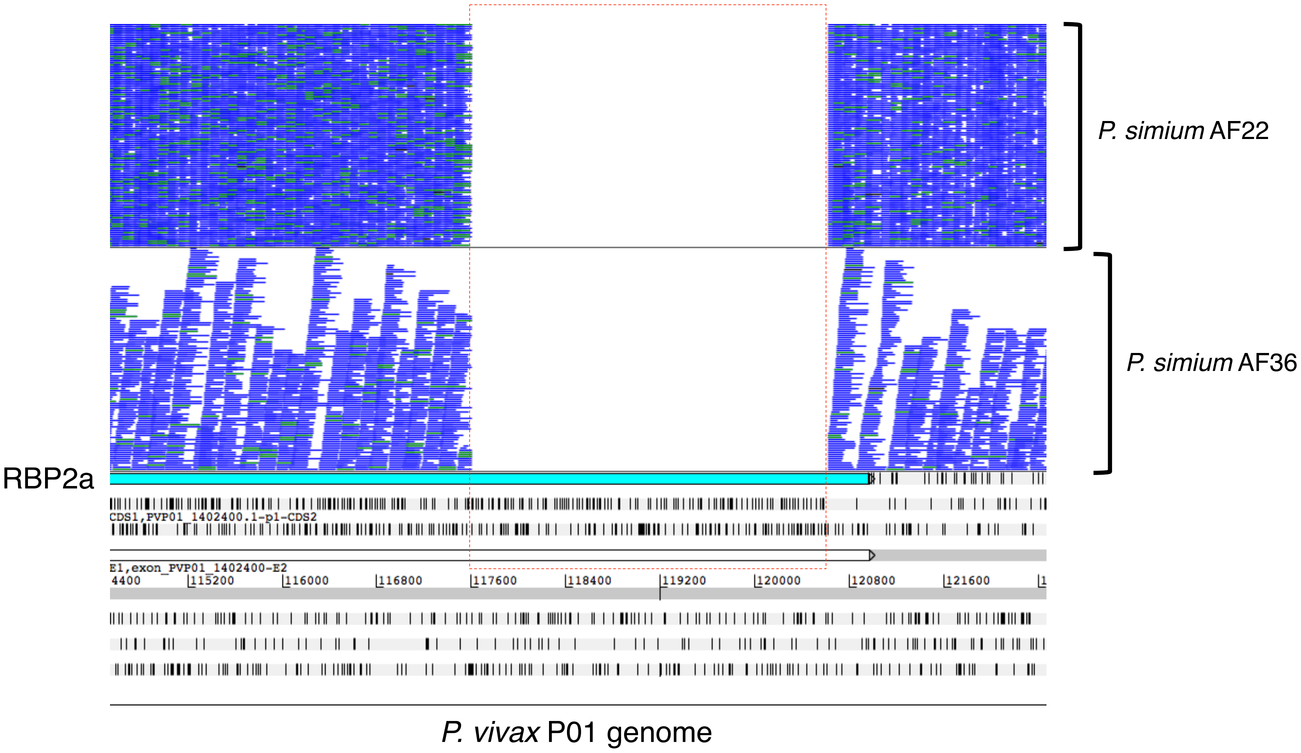
P. simium AF22
P. simium AF36
RBP2a
CDS1,PVP01_1402400.1-p1-CDS2
E1,exon_PVP01_1402400-E2
4400
115200
116000
116800
117600
118400
119200
120000
120800
121600
P. vivax P01 genome

Figure S28

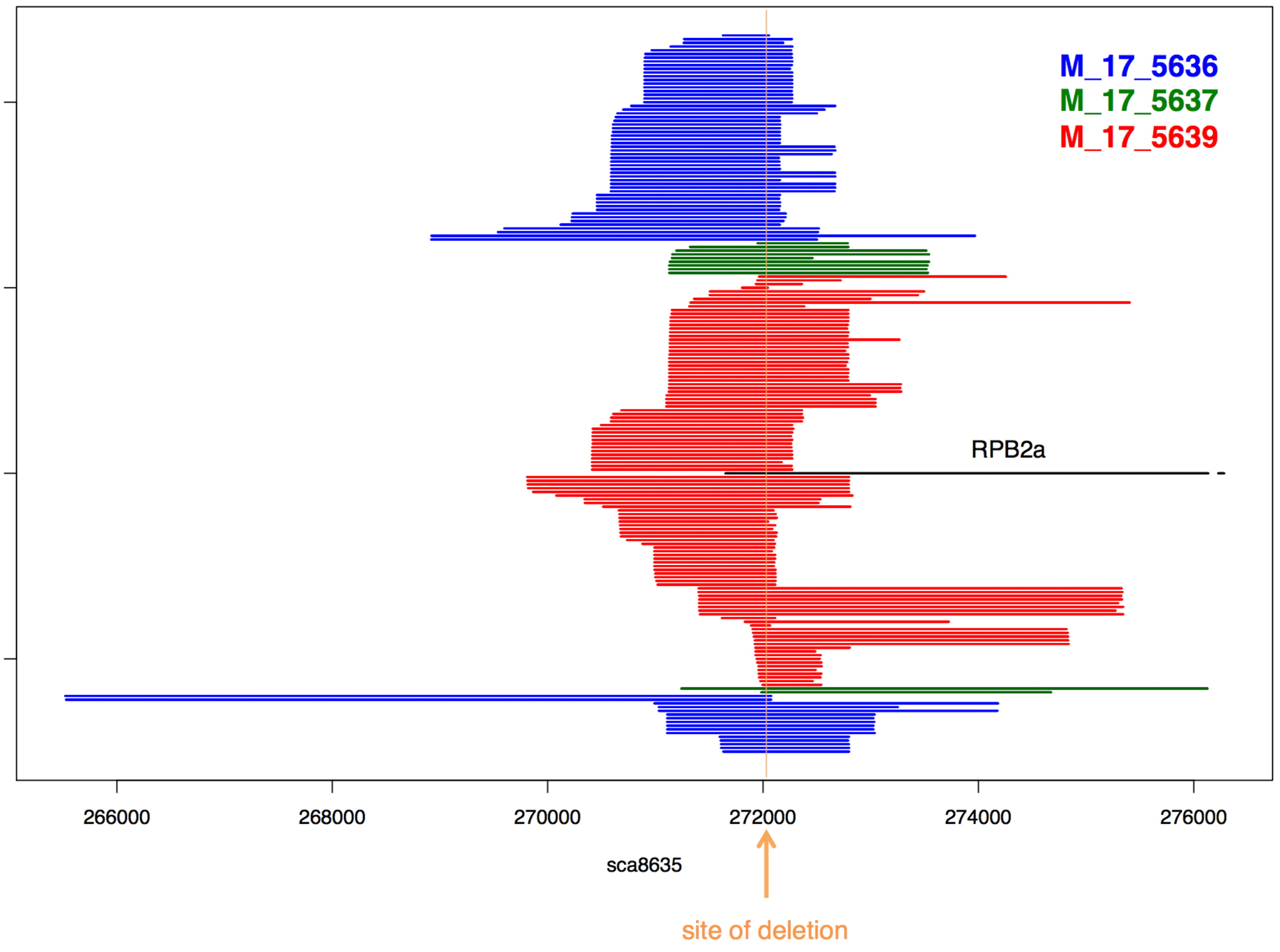

M_17_5636
M_17_5637
M_17_5639
RPB2a
266000
268000
270000
272000
274000
276000
sca8635
site of deletion

Figure S29

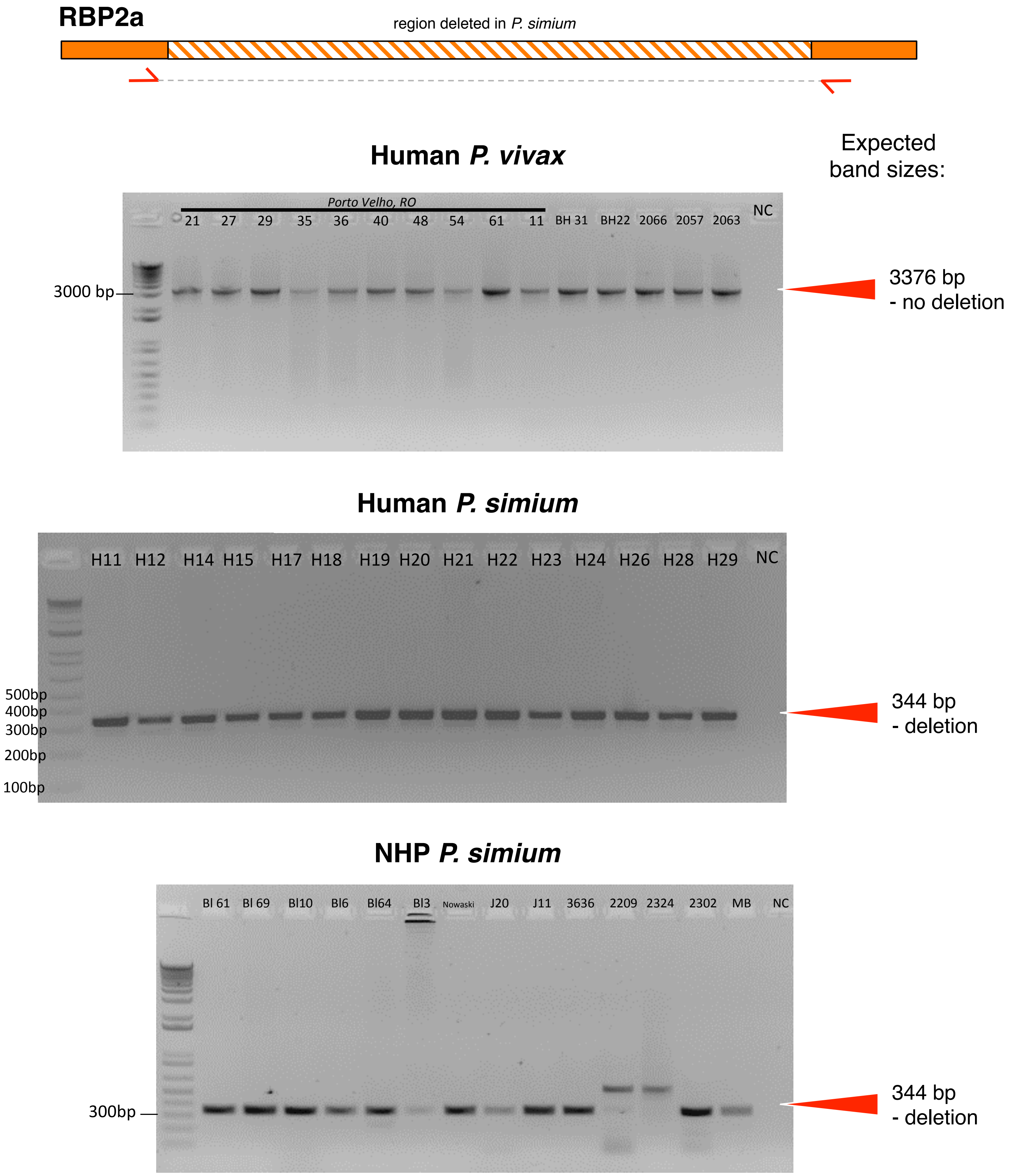


forward primer: AGTGCAAAAGCAGAGGAGG
reverse primer: CCTGCTGCAAAACGGACATT

Figure S30

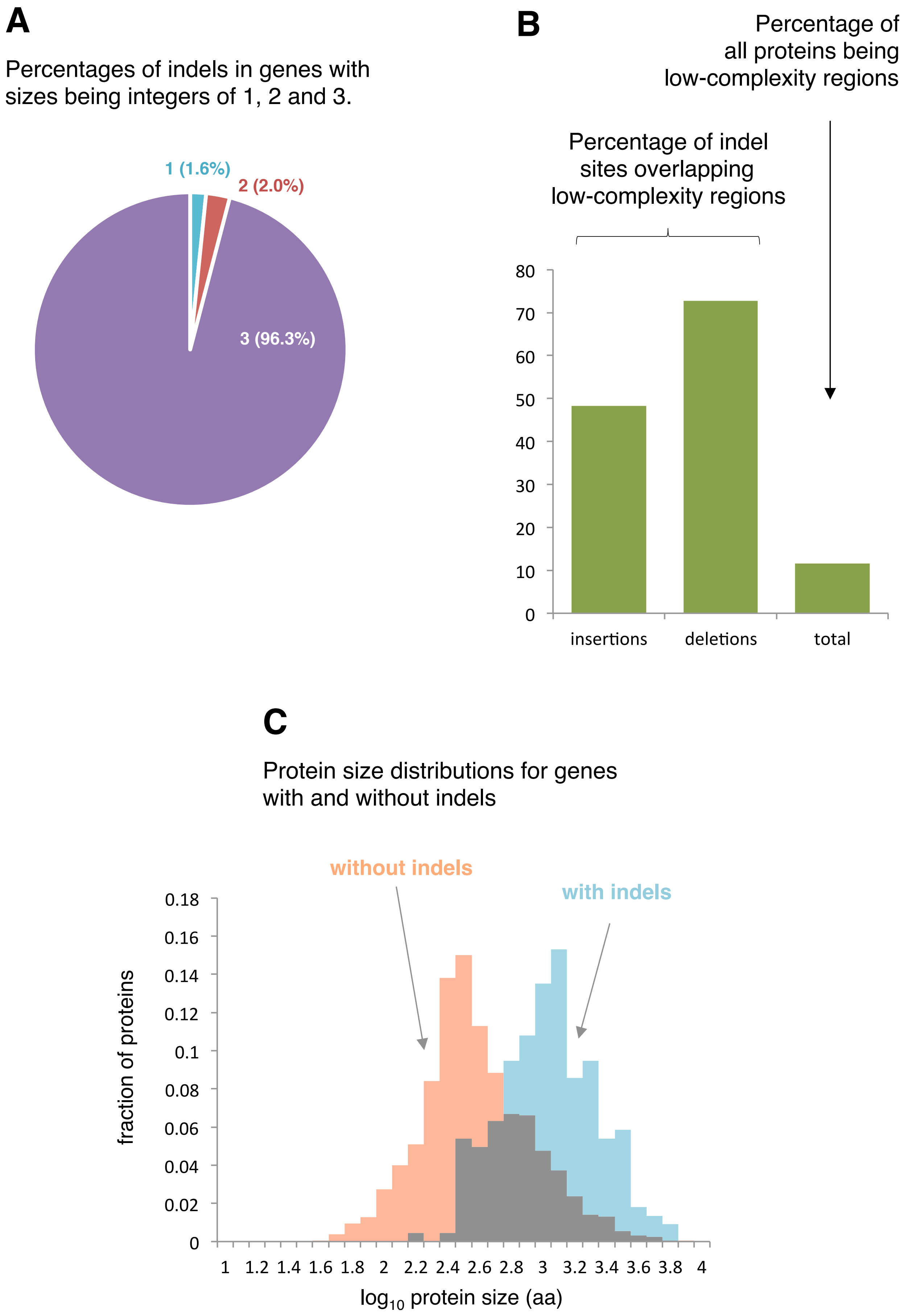
A
Percentages of indels in genes with sizes being integers of 1, 2 and 3.
1 (1.6%)
2 (2.0%)
3 (96.3%)
B
Percentage of all proteins being low-complexity regions
Percentage of indel sites overlapping low-complexity regions
0
10
20
30
40
50
60
70
80
insertions
deletions
total
C
Protein size distributions for genes with and without indels
without indels
with indels
fraction of proteins
0
0.02
0.04
0.06
0.08
0.1
0.12
0.14
0.16
0.18
1
1.2
1.4
1.6
1.8
2
2.2
2.4
2.6
2.8
3
3.2
3.4
3.6
3.8
4
log10 protein size (aa)

Figure S31

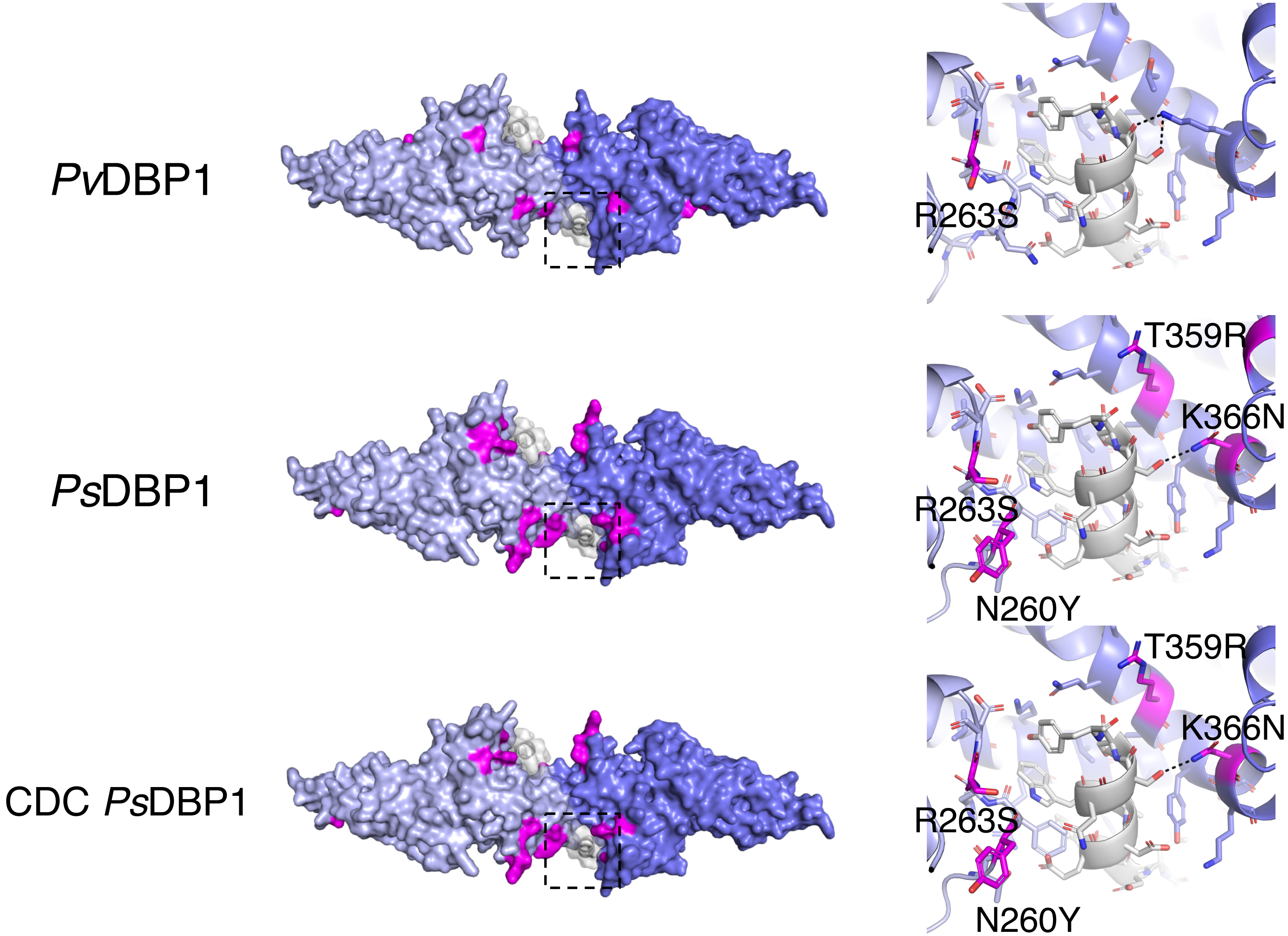
PvDBP1
PsDBP1
CDC PsDBP1
R263S
T359R
K366N
R263S
N260Y
T359R
K366N
R263S
N260Y

Figure S32

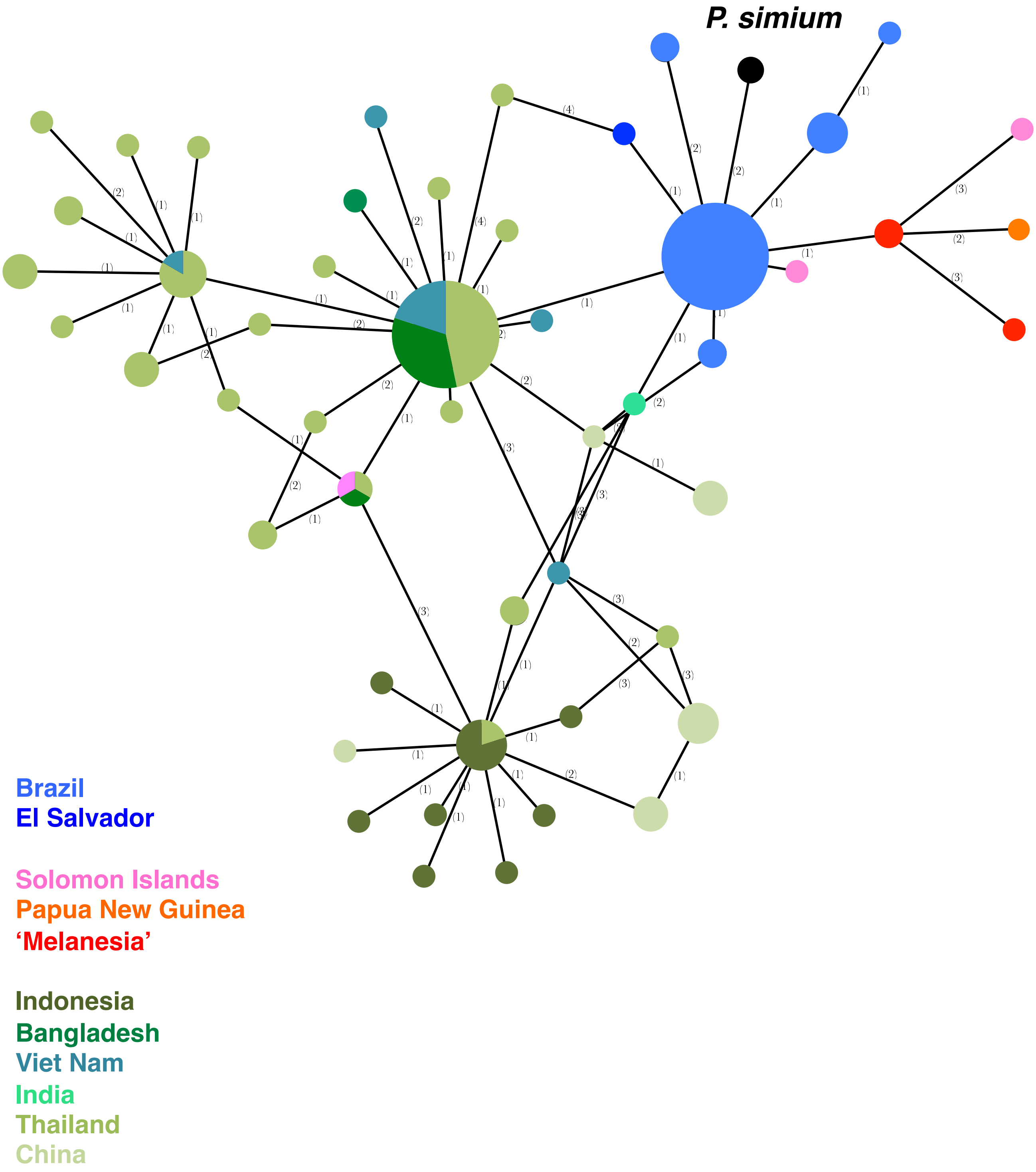

P. simium
Brazil
El Salvador
Solomon Islands
Papua New Guinea
'Melanesia'
Indonesia
Bangladesh
Viet Nam
India
Thailand
China

Figure S33

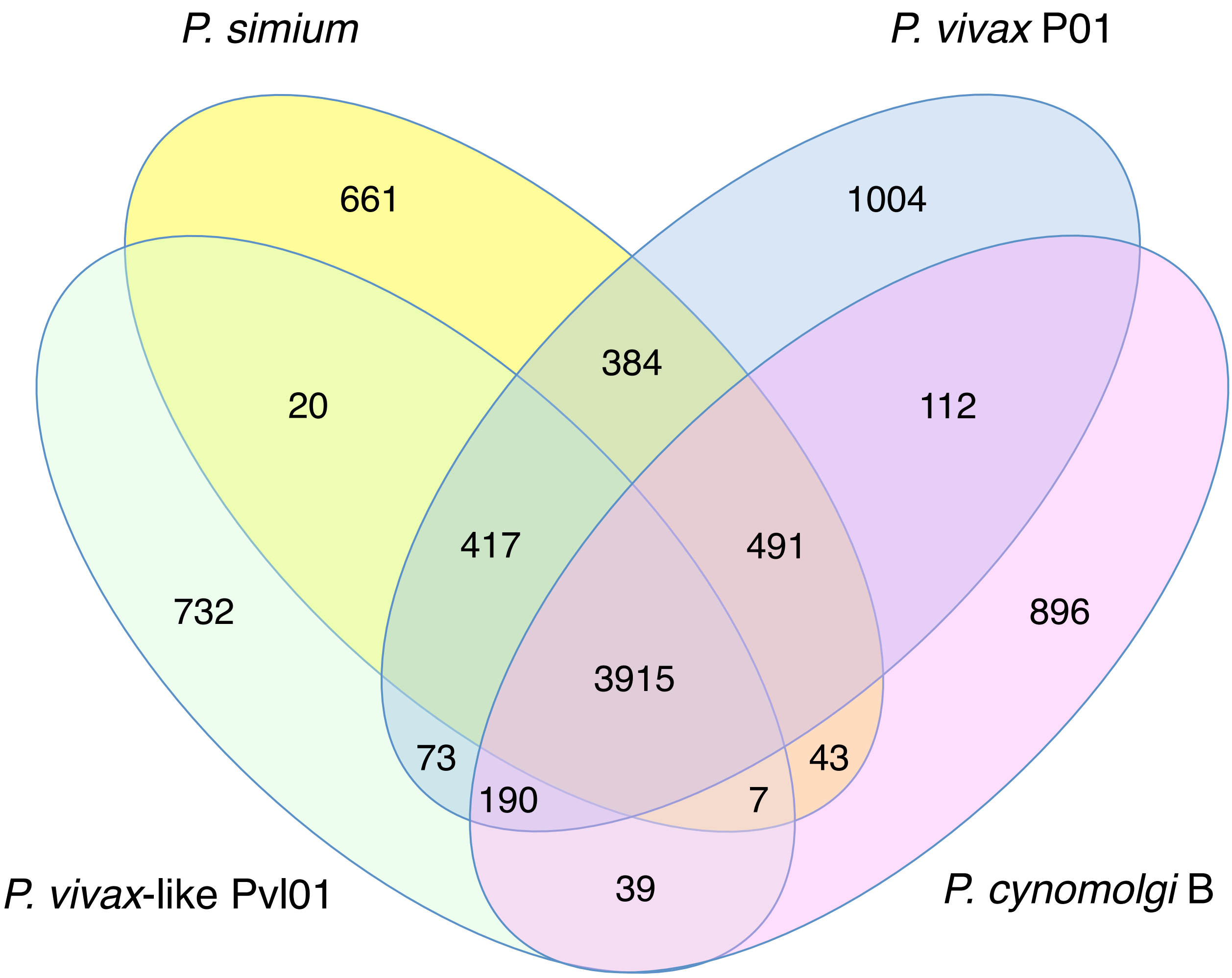

P. simium
P. vivax P01
661
1004
384
20
112
417
491
732
3915
896
73
43
190
7
39
P. vivax-like Pvl01
P. cynomolgi B

Figure S34

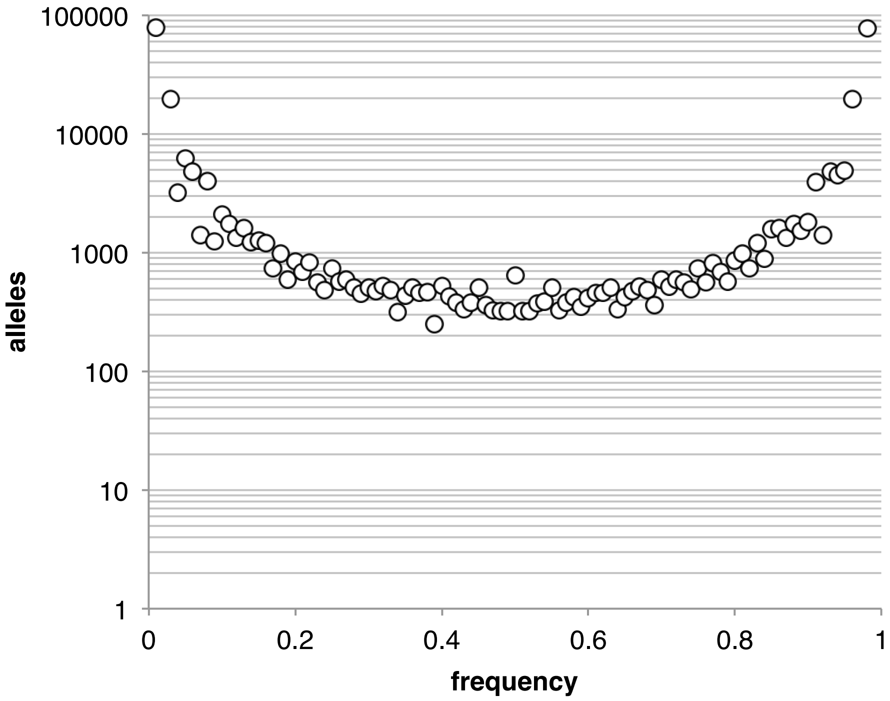

100000
10000
1000
100
10
1
alleles
0
0.2
0.4
0.6
0.8
1
frequency


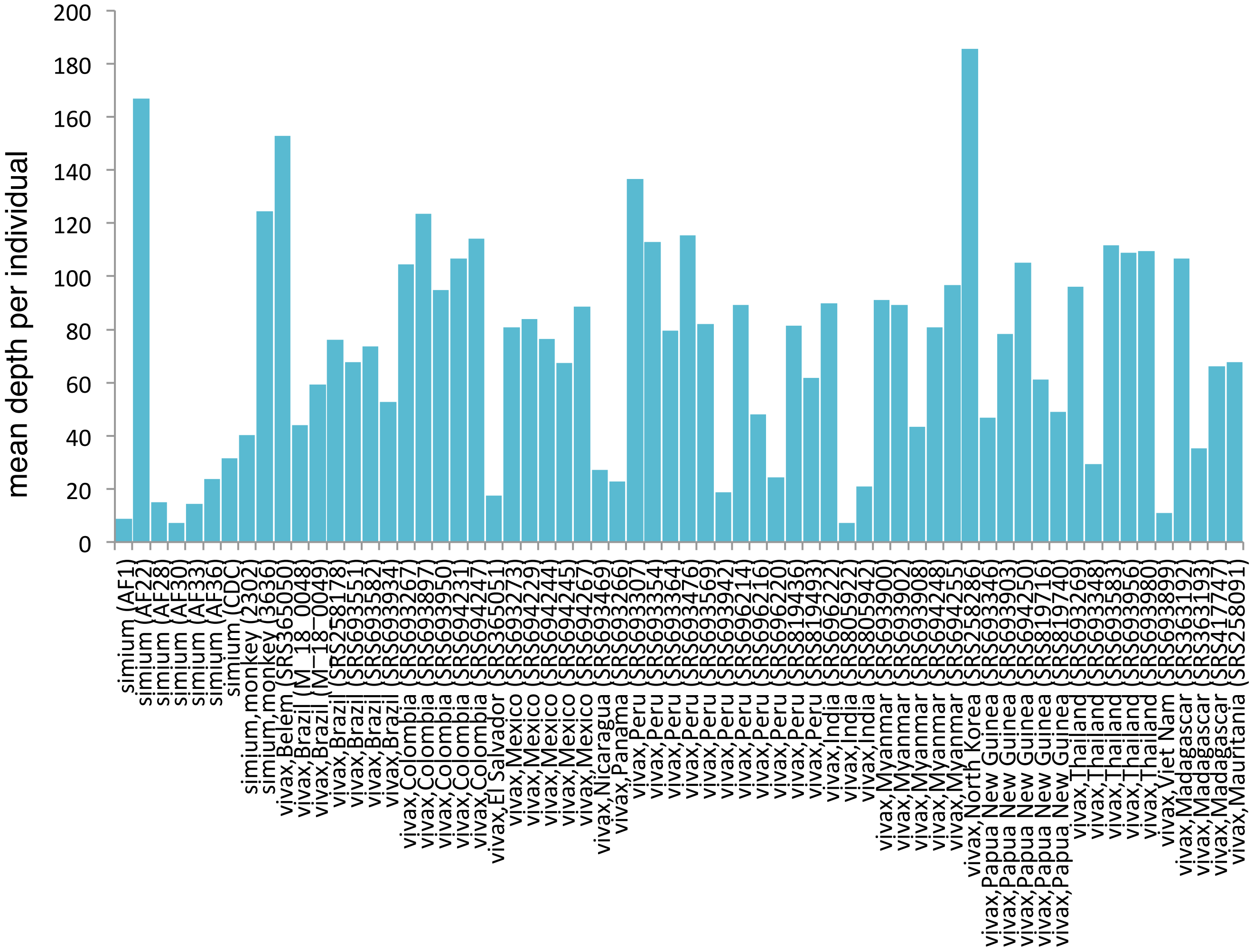

200
180
160
140
120
100
80
60
40
20
0
mean depth per individual
simium (AF1)
simium (AF22)
simium (AF28)
simium (AF30)
simium (AF33)
simium (AF36)
simium (CDC)
simium,monkey (2302)
simium,monkey (3636)
vivax,Belem (SRS365050)
vivax,Brazil (M_18_0048)
vivax,Brazil (M_18_0049)
vivax,Brazil (SRS258178)
vivax,Brazil (SRS693551)
vivax,Brazil (SRS693582)
vivax,Brazil (SRS693934)
vivax,Colombia (SRS693267)
vivax,Colombia (SRS693897)
vivax,Colombia (SRS693950)
vivax,Colombia (SRS694231)
vivax,Colombia (SRS694247)
vivax,El Salvador (SRS365051)
vivax,Mexico (SRS693273)
vivax,Mexico (SRS694229)
vivax,Mexico (SRS694244)
vivax,Mexico (SRS694245)
vivax,Mexico (SRS694267)
vivax,Nicaragua (SRS693469)
vivax,Panama (SRS693266)
vivax,Peru (SRS693307)
vivax,Peru (SRS693354)
vivax,Peru (SRS693364)
vivax,Peru (SRS693476)
vivax,Peru (SRS693569)
vivax,Peru (SRS693942)
vivax,Peru (SRS696214)
vivax,Peru (SRS696216)
vivax,Peru (SRS696220)
vivax,Peru (SRS819436)
vivax,Peru (SRS819493)
vivax,India (SRS696222)
vivax,India (SRS805922)
vivax,India (SRS805942)
vivax,Myanmar (SRS693900)
vivax,Myanmar (SRS693902)
vivax,Myanmar (SRS693908)
vivax,Myanmar (SRS694248)
vivax,Myanmar (SRS694255)
vivax,North Korea (SRS258286)
vivax,Papua New Guinea (SRS693346)
vivax,Papua New Guinea (SRS693903)
vivax,Papua New Guinea (SRS694250)
vivax,Papua New Guinea (SRS819716)
vivax,Papua New Guinea (SRS819740)
vivax,Thailand (SRS693269)
vivax,Thailand (SRS693348)
vivax,Thailand (SRS693583)
vivax,Thailand (SRS693956)
vivax,Thailand (SRS693980)
vivax,Viet Nam (SRS693899)
vivax,Madagascar (SRS363192)
vivax,Madagascar (SRS363193)
vivax,Madagascar (SRS417747)
vivax,Mauritania (SRS258091)
